# Airborne eDNA captures three decades of ecosystem biodiversity

**DOI:** 10.1101/2023.12.06.569882

**Authors:** Alexis R. Sullivan, Edvin Karlsson, Daniel Svensson, Björn Brindefalk, Jose Antonio Villegas, Amanda Mikko, Daniel Bellieny, Abu Bakar Siddique, Anna-Mia Johansson, Håkan Grahn, David Sundell, Anita Norman, Per-Anders Esseen, Andreas Sjödin, Navinder J Singh, Tomas Brodin, Mats Forsman, Per Stenberg

## Abstract

Conserving biodiversity is a global imperative, yet our capacity to quantify and understand species occurrences has been limited. To help address this challenge, we develop a novel monitoring approach based on deep sequencing of airborne eDNA. When applied to a 34-year archive of weekly filters from an aerosol sampling station in northern Sweden, our methods enabled robust detection of over 2,700 genera across all domains of life and estimates of eDNA catchment areas. Reconstructed time series revealed regional biodiversity declines consistent with contemporary, large-scale transformations of forest composition and structure. Our results show airborne eDNA can reliably monitor biodiversity and underscore the immense latent potential in the thousands of aerosol monitoring stations deployed worldwide.

**One-Sentence Summary:** DNA captured from air reveals organisms from all domains of life and their long-term trends.

## Main Text

Humans are driving a global decline in biodiversity (*1, 2*) and the gravity of this crisis remains partially obscured by the difficulty of tracking organisms across time and space. Environmental DNA (eDNA) has emerged as a promising solution to this challenge. Unlike traditional count-based surveys, eDNA can readily detect cryptic taxa (*3*) and archival substrates can grant access to lost or irrevocably altered ecosystems (*4*). These unique features, combined with the logistic demands of traditional monitoring, mean our knowledge of the biodiversity from a given time and place may only extend as far as eDNA permits.

Accumulating evidence from substrates ranging from seawater (*5*) to surface air (*6*–*10*) support eDNA as a source of presence-absence data. More quantitatively, some methods can provide abundance indices (*11, 12*) and diversity estimates congruent with traditional surveys (*13*). In practice, however, eDNA-based applications remain limited due to the stochasticity inherent in ecological processes (*14*) and the errors introduced by existing analytical pipelines, especially false positive detections (*15, 16*).

We demonstrate the potential of airborne eDNA monitoring using a multidecadal archive collected by an aerosol sampling station in northern Sweden. We address some of the most pressing challenges limiting wider adoption of eDNA methods by integrating high-depth metagenomic sequencing with ecological insights. Our approach delineates the spatial footprint of airborne eDNA, robustly determines taxonomic assignments, and uses dynamic models to reconstruct diversity over time. Applied to the filter archive, this allowed us to survey more than 2,700 taxa from all domains of life, recover abundance trends congruent with traditional monitoring, and detect a decline in biodiversity consistent with the effects of contemporaneous forest management.

### Air contains DNA from all types of organisms from a wide range of habitats

#### Airborne eDNA metagenomics

We sequenced near-surface airborne eDNA sampled by a radionuclide monitoring station in the boreal forest of northern Sweden (67.84°N, 20.42°E, see Fig. 1A and supplementary materials). As part of the station’s routine activities, high volumes of surface-level air are continuously pumped (>100,000 m^3^/week) through 0.2 μm glass fiber filters, which are changed weekly and stored long-term in airtight containers. Previously, we found that eDNA can be preserved for decades under these conditions with limited degradation (*6*). We isolated DNA from filters installed during weeks with a mean temperature > 0°C from even-numbered years between 1974 to 2008 for analysis. In total, we generated *ca*. 30 terabases of high-quality metagenomic sequence collected during 380 weeks.

**Fig. 1.**
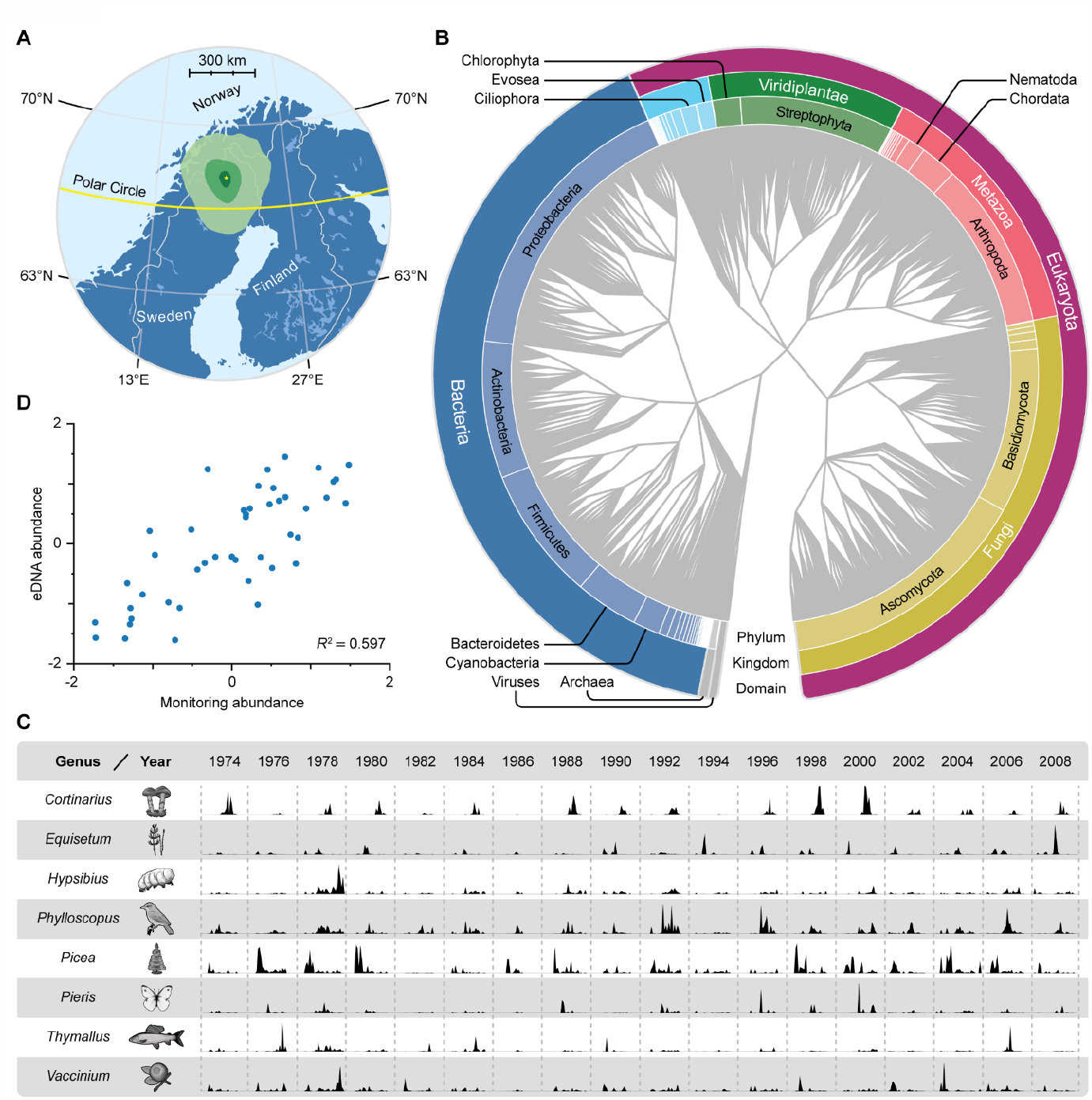
Airborne eDNA provides reliable records of organisms across all domains of life. **A**) Mean modeled origin densities of 22 μm particles in the catchment area during the study period. The differences in intensity of green shading indicate 10-fold differences in density of particles from that area, assuming all areas released the same amount of particles per unit area. The yellow star marks the position of the aerosol monitoring station. **B**) Taxonomic assignments of the 2,739 genera detected in the air filters, according to NCBI taxonomy. **C**) Normalized read proportions from eight genera. **D**) Comparison of taxon abundance estimates from eDNA and point-based surveys estimated for nine bird genera.

#### Accurate taxonomic classifications across all domains of life

Detecting cryptic organisms is a key strength of eDNA, but metagenomic classification methods struggle to balance sensitivity and precision (*17, 18*). We targeted three critical but often neglected steps in a standard pipeline (*19*) for optimization: reference database coverage, parameter choice during read-level classification (*18*), and selection of taxon-level stringency filters (*17, 20*). Combined, these optimizations resulted in a false discovery rate of 4%, a precision of 0.95 and a recall of 0.72 on out-of-sample pseudolabeled test data (supplementary materials). In total, we identified 2,739 high-confidence genera from 69 phyla and 173 classes, in addition to DNA viruses (Fig. 1B, data S5), from a wide range of habitats (Fig. 1C).

The amount of airborne eDNA from a taxon is influenced by their abundance (*11, 21*), habitat (*3, 22*), dispersal mechanisms (*23*), and the source and size of the particles they emit (*22*), among potentially numerous other factors (*12, 22*). Once captured by an air filter, detection probabilities and relative abundances further depend on the eDNA state (*24*), sequencing effort, and the genome sizes and database representation of all organisms contained in an isolate. Wind-dispersed plants, flying insects, and spore-producing fungi are the most abundant taxa in our data, all of which are well-represented in our reference database and on the landscape. However, applying deep sequencing to high-volume air samples also enabled reliable detection of organisms whose particles are less abundant in air, including sixteen genera of fish (Fig. 1C), frogs (*Rana*), moose (*Alces*), reindeer (*Rangifer*) and 41 additional vertebrate genera (Fig. 1B; data S5).

#### Bioaerosol catchments are quantifiable and stable

Spatial footprints, or catchments, for an aerosol sampling station can be estimated from particle sizes and a model of atmospheric conditions. We estimated weekly catchments for three common forest bioaerosols: pine pollen (60 μm), birch pollen (22 μm), and the spores (5 μm) from a typical bracket fungus (Basidiomycota: *Polyporales*) (*25*). Summed over the annual sampling period, these simulations suggested > 50% of 60, 22 and 5 μm particles originate within 20 (± 5.1), 50 (± 17.7) and 310 (± 38.4) km of the aerosol station, respectively (fig. S3). Size distributions for bioaerosols emitted by animals, as well as somatic plant and fungal tissues, are less documented but may fall within the modal 1.0-5 μm fraction ubiquitously documented in bioaerosols (*26*–*28*). More precise catchment estimates require further research (*28*), but we expect airborne eDNA in this study to broadly reflect local plant phenology and landscape-level biodiversity.

Catchment areas for each particle size were broadly elliptical in shape (Fig. 1A) and showed no evidence of systematic changes in size or shape (supplementary materials, data S1). Similarly, we found little indication of seasonal or longer periodicity in wind directions during our annual sampling period 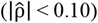 and no support for a relationship between catchment variation and eDNA abundance patterns (data s9, data s10). At local to regional scales, the station’s catchment areas cover a matrix of alpine tundra, montane deciduous forests, open wetlands, and coniferous forests, with smaller components of open water and paved surfaces (supplementary materials, fig. S2). Commercial forest management is extensive at the landscape scale (> 50 km): 1.5% of forests were thinned or felled annually between 1986, the earliest year with reported data, and 2008 in an administrative region roughly congruent with the ≤ 5 μm catchment area (supplementary materials).

#### eDNA abundance indices correlate with traditional surveys

Field experiments in aquatic ecosystems support a strong correlation between abundance estimates from traditional surveys and DNA particle concentrations in natural environments (*11*). Using sequencing data to estimate abundance is considered less promising because read counts provide catch-per-unit-effort (CPUE) data and are always affected by saturation (*29*). As with traditional CPUE surveys, reads can vary proportionally with abundance but how often this holds true for empirical datasets is uncertain (*30*). To test for proportionality, we searched for traditional inventories with sufficient spatial and temporal overlap with the eDNA time series. Data from standardized point-transect surveys for nine bird genera from seven families met this requirement.

Abundance estimates from the traditional surveys explained 60% (p < 0.001) of the variation in log-ratio transformed eDNA abundances (Fig. 1D). Species-specific models could offer further improvements, but the general correlation is already comparable to results from single-species studies in fish using direct DNA quantification (*11*). This shows the potential of using airborne eDNA as an index for population abundances, but we emphasize the need to evaluate each dataset before assuming proportionality.

### Airborne eDNA records seasonal and long-term changes in ecosystem composition

#### Temporal community assemblages

We identified seventeen groups of taxa with similar temporal trends through hierarchical clustering of pairwise log-ratio variances (Fig. 2A, data S5 and S6) (*31*). Seasonal differences divided most organisms along higher taxonomic ranks: eDNA from eukaryotes generally peaked in abundance during a single season, whereas 88% of prokaryote genera were most common during spring and autumn (Fig 2A). A peak consistent with autumn sporulation distinguished most fungi from plants (*32, 33*), and the early spring flowering of trees and dicotyledons separated them from the summer peaks of grasses (*32*) and mosses (Fig. 2A, B). The bimodal seasonality in prokaryotes, however, differed from prior evidence (*33, 34*) and likely results from sequencing effects; that is, organisms with small genomes are most readily sampled when there is little competition in the sequencing pool.

**Fig. 2.**
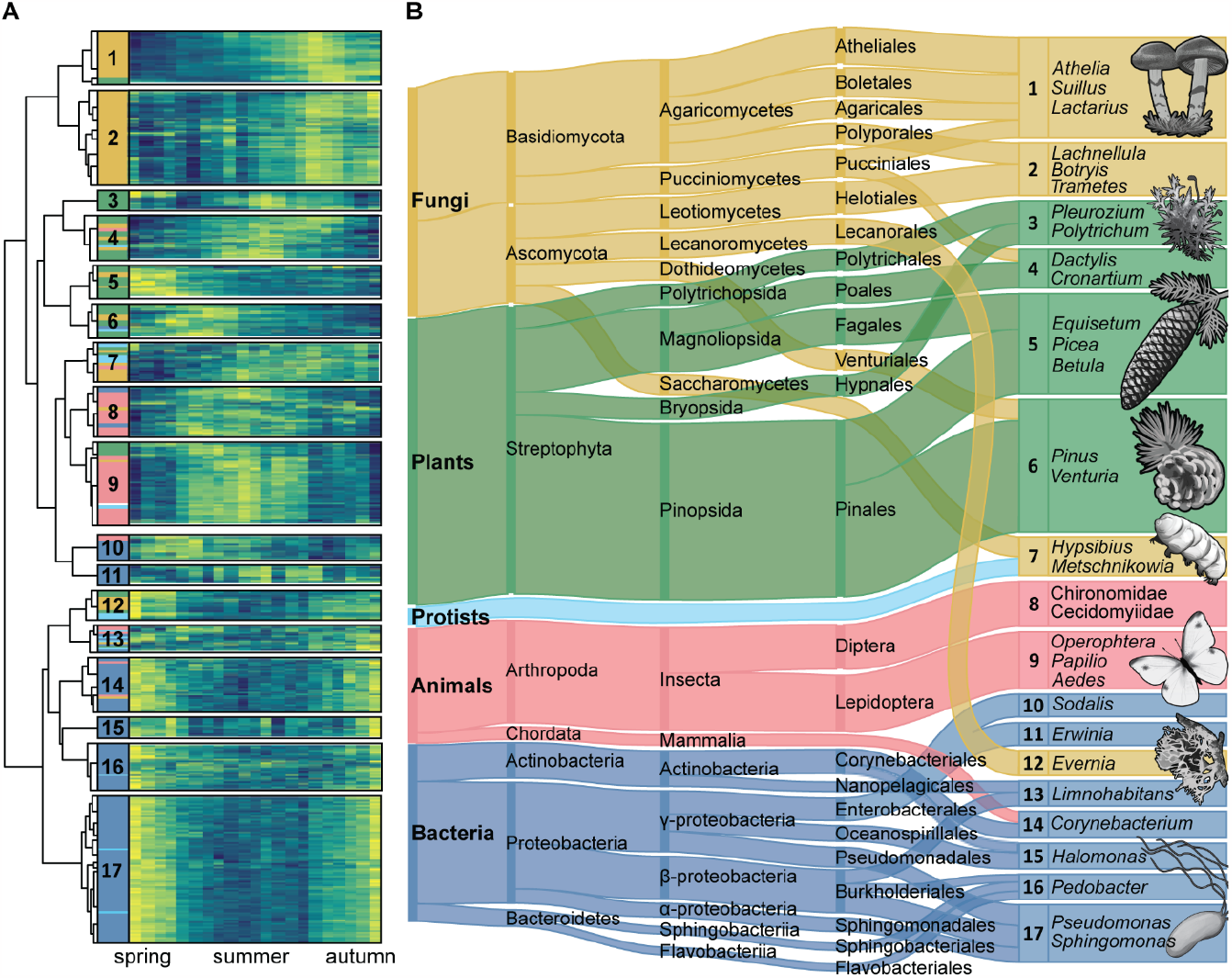
Correlated shifts in abundance reveal temporal assemblages in airborne eDNA. **A**) Hierarchical clustering of the 2,739 genera into 17 temporal clusters by their pairwise log-ratio covariances; stacked bars indicate kingdom membership and the heatmap shows median log-ratio transformed abundances for calendar weeks 21-41 (increasing from dark blue to bright yellow). Cluster sizes are approximately proportional to their taxon richness but note the largest clusters were reduced in size for display. **B)** Taxonomic composition of the clusters from kingdom to order. Protists contains eukaryotes lacking a kingdom classification. Numbered boxes show representative genera for each cluster. Taxonomic groups comprising ≥ 5% of the dataset or a cluster are shown; ribbon and box heights are roughly proportional to rank abundances but the lowest ranks are shown as ties for display.

In addition to phenology, coherent shifts in abundance can result from trophic interactions (*5*). For example, the well-documented endosymbiosis between flies and *Rickettsiales* bacteria (Fig. 2B) and lichenized fungi and algae (Fig 2A) can be detected from their strong temporal covariation (cluster C8 and C12, respectively). This suggests other clusters may reflect undiscovered interactions, such as between putatively endophytic *Venturiales* fungi (*33*) and pine (C6) or the rust fungi and grasses in cluster 4 (Fig. 2B) (*35*). Shared temporal shifts may also indicate a shared response to environmental change (*5*) or aerosolization from a common substrate. A combination may explain the separation between groups of predominantly soil-dwelling (C1) *vs*. endophytic fungi (C2) (*33, 36*) and among bacteria associated with above-ground plant surfaces (C17) (*33, 37*), animal hosts (C14) (*38*), and soils (C16) (*33, 36*). Abiotic conditions, direct trophic interactions, or aerosol emission fluxes that are in turn influenced by the environment (*39*) are all plausible hypothesis for the temporal variation found among microcrustaceans, planktonic bacteria, and other aquatic microbes (C13).

#### Seasonal and long-term cluster dynamics

We partitioned changes in the seventeen clusters into components explained by seasonality, longer-term trends, and environmental parameters using state space models. Ecosystems respond to shifting means, but changes in climatic variability and extremes are expected to be more mechanistically relevant to biota (*40*). To capture some of this complexity, we compared the predictive skill of models using different combinations of latent trend structures and regression matrices, including 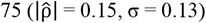 climatic covariates and six comprising a null model of seasonal variation (supplementary materials). The best-performing models predicted 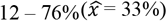 of variation in cluster abundance.

Climatic covariates improved forecasts for eight of the clusters, including all four dominated by plants and three of the four fungal clusters (data S9). Consistent with the timing of pollen and spore release in the boreal region, we found variables related to seasonal transitions to be reliable predictors of fungal and plant eDNA abundance (data S10, data S11). Fungi-dominated clusters generally increased with rain and snow, although eDNA from fungal endophytes (C2) was predictably lower up to 78 weeks after extreme rainfall events (data S10, data S11). Variables related to evapotranspiration were also selected by the models of some plant and fungal clusters, along with the bacterial genera in cluster 11 (data S10, data S11). In general, climatic covariables predicted weekly, seasonal, and cyclic variation but not multiannual or directional trends in abundances (data S10).

After removing the variation predicted by climatic covariates, we found robust evidence of long-term abundance trends in thirteen clusters (Fig. 3A, B; data S10). Most conspicuously, the pine-dominated cluster (C6) increased from 40% of the entire community in the early years of the time series to 80% around 1994 followed by a gradual decline to 60% by 2008 (Fig. 3A). As these are relative abundance trends, a dramatic increase in one component forces declines among the others. However, the trends following this peak indicate a shift in community composition, rather than a saturation artefact driven by a transient spike in pine-associated eDNA. Nine clusters continued to decline even after 1994 and increases in abundance were unequally distributed among the other clusters (Fig. 3A, B). We also detected large abundance changes in some clusters in 2008, the end of our time series, which could indicate nascent trend reversals (Fig. 3B).

**Fig. 3.**
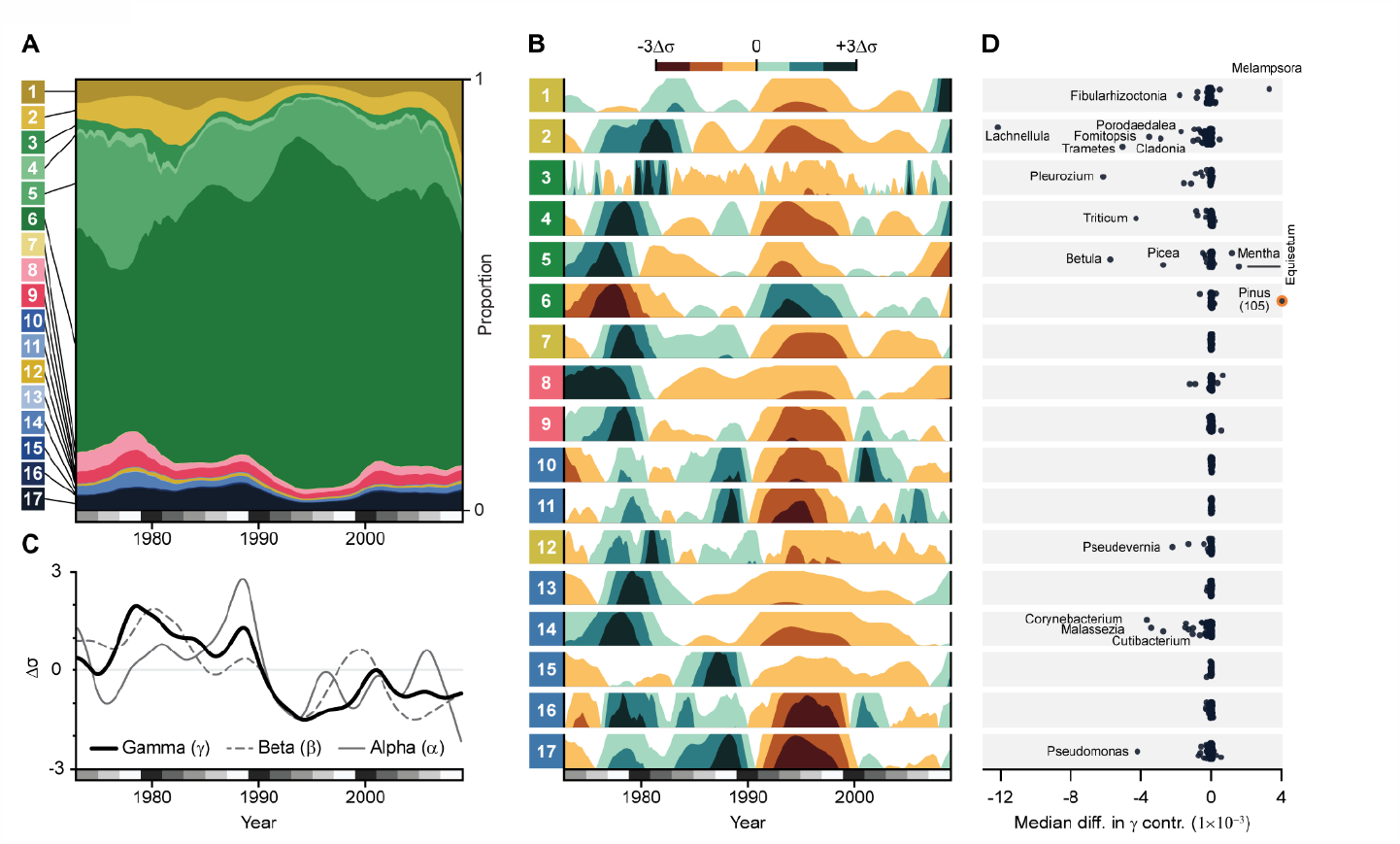
Airborne eDNA records seasonal and long-term changes in ecosystem composition. **A**) Relative abundances of each cluster across the time series. **B**) Centered and scaled relative abundances of each cluster. **C**) Modeled α, β, and γ diversities for the total composition. **D**) Median difference in gamma contributions between 1974-1988 and 1992-2008 for each genus in the 17 clusters. The value of *Pinus* (orange circle) in cluster 6 has been truncated (actual value in parenthesis).

#### Biodiversity loss from declines in forest taxa

We used transformations of the Rényi entropies (*41, 42*) to partition changes in biodiversity into evenness and distinctiveness components (supplementary materials). This framework extends the logic of Hill numbers (*43*) to relative entropy (β) and cross-entropy (γ) to obtain unified families of diversity indices. Higher α diversity indicates a more even relative abundance distribution whereas β increases as taxa are temporally structured. Changes in γ diversity occur through either, or both, of these components and indicate that biodiversity in a broad sense is unevenly distributed across time.

Mean γ diversity declined between 1990 and 1994 (Fig. 3C), concurrently with the rapid increase of the pine cluster. Despite an increase from the mid-1990s, γ diversity averaged 35% lower (95% CI: 31-40%) between 2002-2008 than 1974-1988, a loss equivalent to *ca*. 31 effective taxa. Evenness decreased modestly but consistently over the same period, from 22 to 20 effective taxa (95% CI:17-30 to 15-27), although a steeper decline may have begun in 2008. This means the decline in γ diversity mostly resulted from a change in distinctiveness, with taxa more disproportionately abundant in 1974-1988 than in 2002-2008. Reducing the influence of rarer taxa (q = 2, 3) or restricting the analysis to different taxonomic subsets did not change this pattern of biodiversity loss (data S10).

Diversity metrics are not necessarily positively correlated with ecosystem health. Generalist and invasive taxa can increase diversity (*44*), even though their success often increases with environmental degradation (*45*). We identified the taxonomic drivers of the diversity decline by comparing per-taxon γ contributions from 1978-1988 *vs*. 1992-2008. Consistent with the cluster trends, we found a large increase in in the γ contribution of pine (Wilcoxon signed-rank test, Benjamini-Hochberg adjusted p-value < 0.001) and numerous declines in core taxa like birch (*Betula*; p < 0.05), spruce (*Picea*; p > 0.05), feathermoss (*Pleurozium*; p < 0.001), tree and ground-dwelling lichens, and wood-dwelling fungi (all p < 0.001), among other taxa with uncertain ecologies (Fig. 3D). These genera, and the species within them, occur in different habitats but are all directly affected by forest management (*46*–*48*).

Productive forests (capable of producing > 1 m^3^/ha year^-1^) in Fennoscandia are most frequently clearcut, replanted with seedlings, and thinned multiple times before they are felled again. While effective for timber production, this silvicultural system has converted a structurally-diverse landscape to a mosaic of monocultures. Between 1974 and 2008, primary forests in the region declined by > 50% and more clearcuts occurred within 100 km of the filter station in the 1980s than any earlier period in the 20^th^ century (supplementary materials). These forests were disproportionately replaced by pine, consistent with the long-term increase of pine-associated eDNA. On-the-ground management activities may also create bioaerosol pulses that influence shorter-term eDNA trends: the 1990-2000 maxima in the pine cluster coincides with a period of extensive harvests and reforestation in the region (supplementary materials).

Population declines in taxa dependent on old forests, including both *Porodaedalea* species in the region and *Fomitopsis rosea*, one of the two species in this genus potentially represented in our data, (Fig. 3D) are widely documented in Sweden (*49*). Rare, specialist species like these are naturally vulnerable to environmental changes, but we also detected large γ declines in genera common in young, natural forests: *Pleurozium, Trametes*, and *Fomitopsis pinocola* (Fig. 3D). Field-based studies have more recently emphasized the threats to these and other core genera posed by soil scarification (*50*), insufficient dead wood quantity or quality (*25*), habitat fragmentation (*51, 52*), or the altered light and moisture regimes from high planting densities and fire suppression (*46, 48*). Together, this suggests the largest change in airborne eDNA diversity resulted from commercial forest management across the landscape.

## Conclusions and outlook

We demonstrate the ability of airborne eDNA to detect the contemporary presence of organisms across the tree of life, track shifts in ecosystem composition, and provide quantitative abundance indices. While this marks a notable improvement in the resolution and scope of eDNA biodiversity monitoring, amenability to reanalysis is a key benefit of our dataset. Most (76%) of our reads are unclassified, an unsurprising result given that only a tiny sliver of species have reference sequences (*53*). With more extensive reference databases, future reanalysis of this dataset will continue to provide insights into biodiversity at multiple levels of organization, including the gene pools of individual species (Fig. 4A). We here focused on relative changes between 17 clusters, but relative changes within any given subcomposition can also be investigated (see Fig. 4B).

**Fig. 4.**
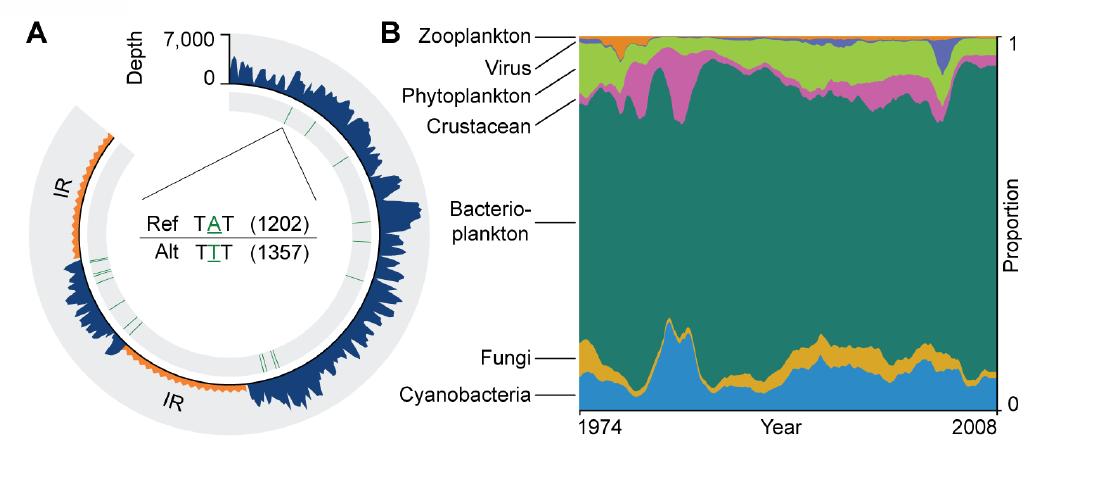
Data support analysis of sub-compositions, individual species and their genetic diversity. **A**) Mapping reads from seven spring weeks in 1998 to the *Betula nana* chloroplast genome (161 Kbp). The y-scale indicates mapping depth and red lines indicate single nucleotide variants relative to the reference genome. In the center, the number of reads supporting two observed sequence variants at one of the positions is shown. IR: inverted repeat regions, where reads cannot be uniquely mapped. **B**) Reclosing the data solely for taxa included in cluster 13 (i.e., holding the total abundance of those taxa constant across time), dominated by aquatic microorganisms, and fitting individual models for those taxa, reveals that changes in abundance of taxa of similar type tend to be more similar (fig. S17). Summation of the relative abundances by type reveals distinct trends for different types of organisms, indicating that ecological interactions could be investigated.

Our study underscores the value of aerosol stations as serendipitous collectors of biodiversity data (*10*). Our results suggest the high flow rates (500-1,500 m^3^ h^-1^) used in radionuclide detection also enable detection of even organisms that do not readily emit bioaerosols. Similarly to air quality networks (*10*), radionuclide stations operate worldwide under standardized protocols. Europe alone hosts more than 400 stations (*54*) and those surveilling for the Comprehensive Nuclear-Test-Ban Treaty Organization (CTBTO) are strategically positioned to maximize global coverage (*55*). Airborne eDNA from these and other already operational networks may provide an unprecedented opportunity to reconstruct ecological history and detect ongoing changes almost in real-time.

## Acknowledgements

We thank Catharina Söderström and Johan Kastlander (CBRN Defence and Security, Swedish Defence Research Agency) for providing access to the air filter archive, and Benedicte Albrectsen and Göran Englund for their feedback on previous versions of this manuscript. We also acknowledge support from the Science for Life Laboratory and the National Genomics Infrastructure (NGI) for providing assistance in massive parallel sequencing. The computations were enabled by resources provided by the National Academic Infrastructure for Supercomputing in Sweden (NAISS) and the Swedish National Infrastructure for Computing (SNIC) at UPPMAX and HPC2N partially funded by the Swedish Research Council through grant agreement nos. 2022-06725 and 2018-05973. Modified Copernicus Climate Change Service information 2020 was used for the catchment area analysis. Neither the European Commission nor European Centre for Medium-Range Weather Forecasts (ECMWF) is responsible for any use that may be made of the Copernicus information or data it contains.

## Author contributions

PS, MF, TB and EK conceived and designed the study; EK and AMJ extracted DNA; DSv constructed the database and performed read classification; EK, ARS, DB, DSv pre-processed the data; ARS designed and implemented the machine learning approach; and HG constructed the particle models. ARS and EK conducted most of the data analysis, with support from DSv, DB, ABS, JAV, AM, DSu, BB, AN, AS, NS, and PAE. EK, ARS, DSv, BB, PS and NS wrote the first draft of the manuscript. All authors contributed intellectual input and approved the final version.

## Data and materials availability

Sequencing data are available through the NCBI Sequence Read Archive under project PRJNA808200.

### Supplementary Materials

Materials and Methods

Figs. S1 to S17

Tables S1 to S5

References (56–131) Data S1 to S11

## Supplementary Materials for

## Materials and Methods

### Summary of supplementary materials and methods

DNA was extracted from weekly air filters sampled in even-numbered years from 1974 to 2008 by a radionuclide aerosol monitoring station in northern Sweden. DNA isolates from each week were shotgun sequenced on their own Illumina NovaSeq 6000 S4 flow cell. Reads were subjected to quality control and then taxonomically classified using a large custom reference database. Classification stringency parameters were optimized for genus-rank using publicly-available species observations made in the vicinity of the aerosol monitoring station. Read counts per genus were log-ratio transformed and detrended to remove potential biases from *e*.*g*., read length variation. The last processing step removed putative false positive taxa using a novel machine learning approach (fig. S1).

The final 2,739 high confidence genera were then clustered based on shared temporal patterns. Time series analysis of the resulting 17 cluster abundances, community diversity components, and of individual genus abundances was performed using Bayesian state space models. The relative predictive power of covariables representing endogenous seasonal patterns, variation in climate and weather-related parameters, and weekly changes in the aerosol station’s catchment area were compared using leave-future out cross validation.

**Fig. S1.**
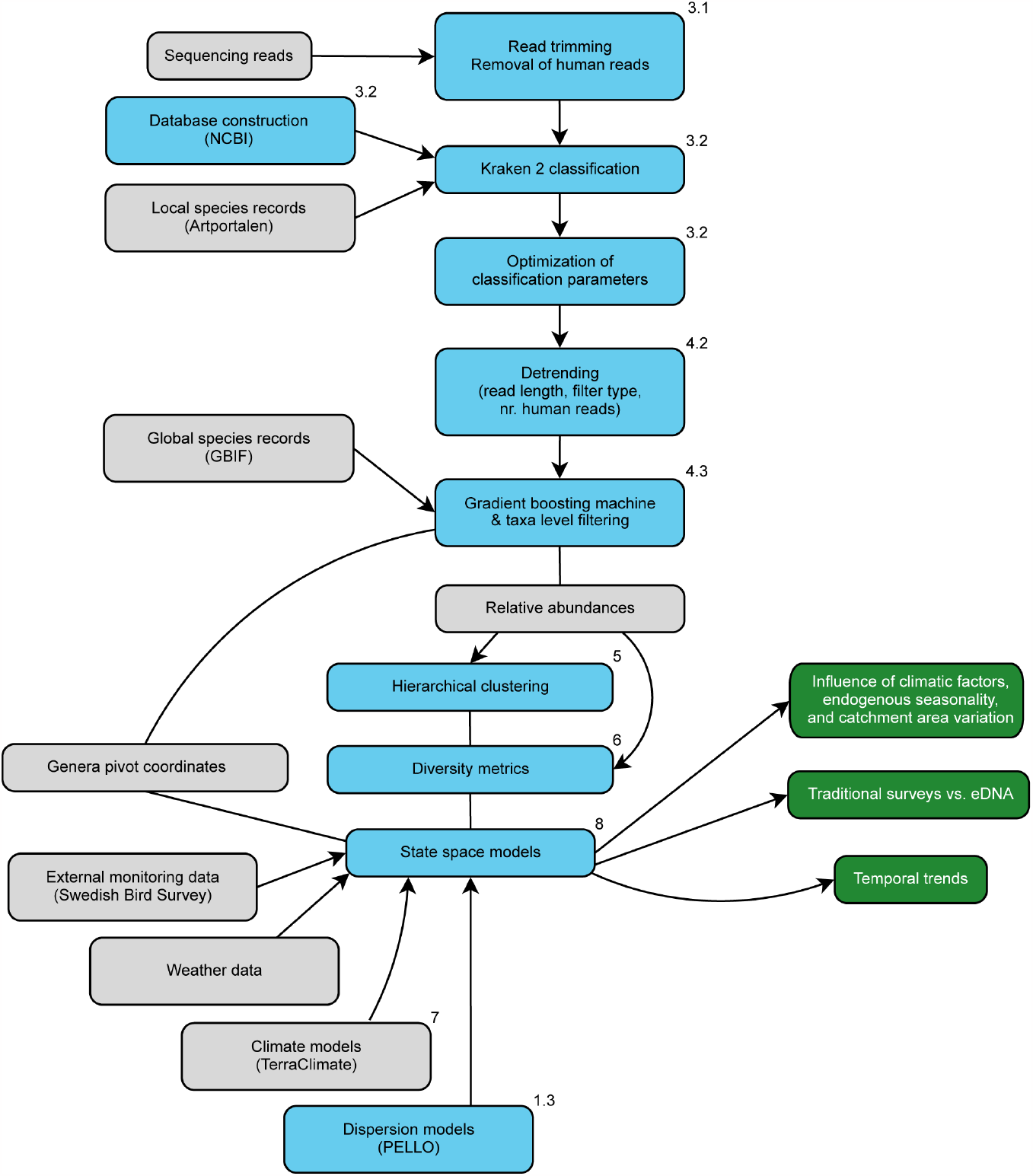
Brief summary of the analysis workflow. For more details of each step, see the indicated section.

## 1. Aerosol sampling station and catchment area

### 1.1 Ecological context

The aerosol sampling station is located in the province of Norrbotten, the northernmost in Sweden, *ca*. 9 km east of the mining town of Kiruna (population 23,000, 67.84°N, 20.42°E) in the northern boreal zone. Data on land cover were extracted from the Swedish National Land Cover Database (*56*), mapped in 2017-2019. The data consist of a base map with 25 thematic classes in three hierarchical levels and has a raster format with 10 m pixel size. Using ArcGIS *v*. 10.3, we extracted the area and proportion of land cover classes within 50, 20, 5, 0.5, and 0.1 km radius of the aerosol sampling station. Thematic classes were aggregated into nine classes. Forests outside and on wetlands were not separated.

Land cover within 50 km was dominated by vegetated open land (39%; mainly low and middle alpine belts), open wetland (20%), coniferous forests (15%), deciduous forests (11%), mixed forests (6%), and water (6%; fig. S2). Minor classes included temporarily deforested land (1.8%; clearcuts), open land without vegetation (0.8%; mainly high alpine belt), and artificial vegetation-free surfaces (0.7%; e.g. mining areas, building, and road/railway). Agriculture was uncommon. Forested area increased from 32% at the 50 km scale to 64% at the 0.5 km scale with a commensurate decrease in open habitats. Land cover within 0.1 km of the aerosol sampling station was composed of 75% forest, 12% open wetland, 8% artificial surfaces, and 5% other land cover.

In 2020, we inventoried a total of eleven 10 m radius plots located at 25 (four plots) and 100 m (seven plots) distance from the station. We recorded the diameter at breast height (DBH; 1.3 m) and species of trees with DBH ≥ 10 cm and calculated basal area per hectare. At this scale, the forests were dominated by pine (71% of basal area), followed by spruce (18%), and birch (11%). They were old, multi-layered, and semi-open (mean basal area 15 m^2^ ha^-1^). The forests had a semi-natural character, with a few old stumps indicating past selective logging. Understory vegetation was dominated by dwarf shrubs (*Empetrum nigrum* ssp. *hermaphroditum, Vaccinium myrtillus, V. vitis-idaea*) and bryophytes (*Hylocomium splendens, Pleurozium schreberi*), with patches of terricolous lichens (e.g., *Cladonia* spp., *Nephroma arcticum, Peltigera* spp.).

**Fig. S2.**
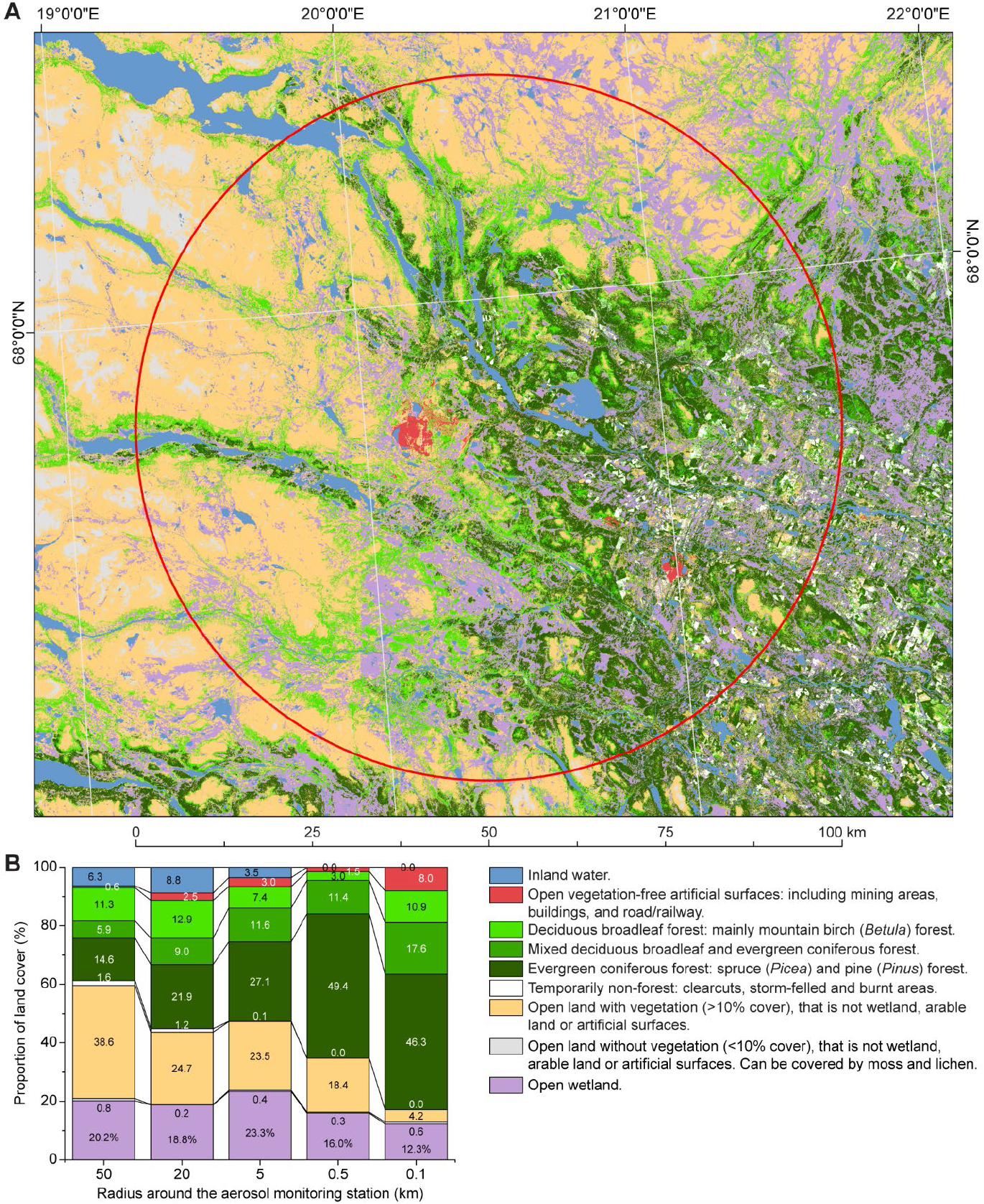
Land cover around the aerosol sampling station. **A**) Map of nine land cover classes in a 50 km buffer around the aerosol monitoring station. **B**) Relative composition (%) of land cover at five different spatial scales (50 km to 0.1 km). Based on land cover data with 10 x 10 m pixel size.

### 1.2 Aerosol sampling

Air filters were collected once a week between 1974 and 2008 by the Swedish Defense Research Agency (FOI) to monitor radioisotopes in surface level aerosols (*57*). The filters belong to a larger collection spanning the five decades of continuous and ongoing radionuclide surveillance at stations across Sweden. Filters are made of glass fiber with a pore size of 0.2 μm and filter more than 100,000 m^3^ of air each week. The manufacturer changed in 1996 (from Camfil type CS 5.0, Camfil Svenska AB, to HB5773, Hollingsworth & Vose Company Ltd.), but the new filters were produced with the same specifications. We detrended the sequence data (section 4.2 Detrending) to account for potential effects of the filter manufacturer change. From 1976-1984, filters were stored in rectangular plastic containers and in cylinder shaped containers in all other years. We selected weekly air samples from every other year between 1974 and 2008. We attempted DNA extraction from filters installed during weeks with a mean temperature > 0°C because aerosol DNA concentrations are low during freezing conditions (*6*). The air filters were randomized and coded prior to DNA extraction.

### 1.3 Catchment area estimation

Bioaerosols are airborne particles released into the atmosphere such as fungal spores, bacteria, pollen, and shed cells. During their journey to an aerosol sampler, bioaerosols undergo processes such as deposition and coagulation and interact with atmospheric moisture as they are carried by complex and chaotic wind patterns. These processes determine the spatial extent of sources sampled by the aerosol station, which we refer to as catchment areas.

We employed PELLO (*58*), a random displacement Lagrangian particle model validated (*59*) and applied in several studies (*9*–*11*), to estimate catchment areas. PELLO is normally used in applications with some basic knowledge of the source (*i*.*e*., position and characteristics of the pollutant released in the atmosphere), but we lacked two important source properties: position and time. The straightforward solution to this problem is to define a large number of sources covering the entire calculation domain both in time and space and then keep track of all aerosols that enter the filter station. This was unfeasible in our scenario due to the large number of sources we would have to define, and hence the large number of model particles to handle, to cover the region of interest in time and space. Our approach was therefore to use an adjoint version of PELLO where model particles advected with wind and dispersed due to turbulence backward in time from the aerosol station to their origin. This backward simulation let us define only one source, but spread in time, which shortened the computation time by several orders of magnitude.

PELLO models particle transport with data from numerical weather predictions (NWP) from the European Centre for Medium-Range Weather Forecasts (ECMWF). For this study, we used the ERA-5 dataset (*60, 61*) (1980-2008, except 1994 as the data for that year could not be retrieved) with a 1.0 x 1.0° horizontal resolution and a vertical resolution of 79 hybrid sigma pressure levels (in ERA-5, this is the lowest 16 km of the atmosphere). We used a 6 and 12 hour forecast step starting at 06:00 and 18:00, resulting in four forecast fields per day. The spatial domain of the weather data covered Europe, including western part of Russia and Northern Africa. Aerosol dry and wet deposition were modeled but no other biological or chemical particle properties were incorporated.

As a source for the adjoint dispersion, we used particles with diameters of 5, 22 and 60 μm and a density of 800 kg/m^3^ (*62*), representing smaller fungal spores or larger bacterial cells, birch pollen, and pine pollen, respectively. The spatial domain of the release of bioaerosol for the adjoint dispersion was defined with a horizontal domain of 30 x 30 m and a vertical domain stretching from 0-300 m, roughly corresponding to the planetary boundary layer in a neutral atmosphere. The source domain represents a ground source on the regional scale where the bulk of the bioaerosols are well mixed in the planetary boundary layer. Although we only modeled three different particle diameters, we expect it to provide a rough estimation of the catchment area within this regional context.

We summarized the spatial extent of the catchment areas and the proportion of particles originating from eight cardinal directions. We calculated the particle mass originating from different distances from the aerosol sampling station (2, 5, 10, 20, 31, 50, 100, 180, 310, 520, and 860 km) for each week in the even-numbered years from 1980 to 2008 (except 1994 as previously described) for the 22 μm particle and for each week in 1988 for the 5 and 60 μm particles (data S1). For each particle size, we calculated the cumulative mass within each radius scaled by the sum within the 860 km radius area (fig. S3). Yearly averages of the proportion of particle mass originating from each cardinal direction were also calculated for each distance (data S1). We used these weekly sums for the 22 μm particle size as regression covariates in the time series analysis in Section 8. This allowed us to assess the potential influence of changes in bioaerosol sources on weekly eDNA compositions.

**Fig. S3.**
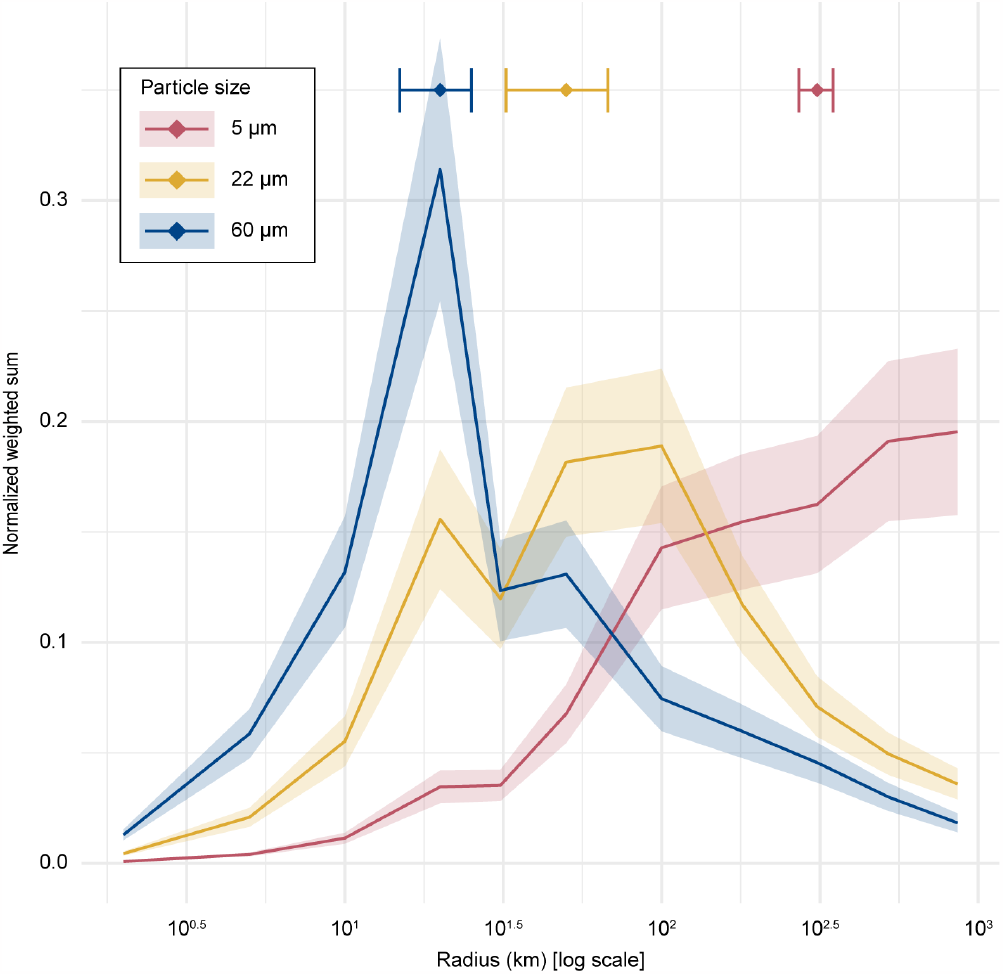
Particle dispersion bootstrapping and Monte Carlo simulation results. Normalized weighted sums (*i*.*e*., contributions from various distances closed to one) plotted against the distance (log-scale) from the aerosol sampling station, color-coded according to particle size. Shaded areas correspond to the normalized standard error obtained from the bootstrap procedure for each particle size. Horizontal error bars (standard deviation) and data-points at top of plot correspond to the results of the Monte Carlo simulation equal to 50% of cumulative particle mass from all directions, color-coded according to particle size, using the block bootstrapping as input.

To assess the range of particle dispersion and its associated uncertainty, block bootstrapping with *R* package ‘boot’ *v*. 1.3-28.1 (*63, 64*) was employed. Each bootstrap replicate consisted of 1,000 resamples with a block size of four weeks, approximating a lag of one month. The bootstrapped data were then normalized using weighted sums (fig. S3). To identify the 50% cumulative particle mass originating from all directions, a weighted Monte Carlo simulation was conducted, using the normalized weighted sums and their standard errors as input parameters over 1,000 draws.

To evaluate the year-to-year variation in the shape of the catchment area for the even years between 1980 and 2008, a linear mixed-effects model was implemented using the R package ‘nlme’ *v*. 3.1-163 (*65, 66*). The dependent variable was the scaled particle mass value, normalized to sum to one. Fixed effects included the year and the cardinal direction, as well as their interaction. A random intercept for the year was included to account for repeated measures, along with a first-order autoregressive correlation term to handle autocorrelation. The mixed-effects model indicated no significant year-to-year variation in the shape of the catchment area across the studied period (table S1).

**Table S1.** Catchment area linear mixed-effect model results.

| effect | estimate | std. error | df | t-value | p-value |
| --- | --- | --- | --- | --- | --- |
| intercept | 0 | 0.86 | 1204 | 0.00 | 1.00 |
| year (fixed effect) | 0 | 0.00 | 12 | 0.01 | 0.99 |
| cardinal directions (fixed) | not shown individually |  |  |  | n.s. |
| year × cardinal direction | not shown individually |  |  |  | n.s. |

## 2 DNA sequencing

### 2.1 Extraction

The DNA extraction protocol was adopted from (*6, 67, 68*) with a few modifications. For each air filter, three punches were punched out within a sterile plastic bag using a biopsy punch (Ø8 mm, Integra Miltex, Plainsboro, NJ, USA) and collected in three separate 2.0 mL screw cap tubes containing 1.0 g of 0.1 mm zirconia/silica beads and 0.5 g of 1.0 mm zirconia/silica beads (BioSpec, Bartlesville, OK, USA). Prior to extraction, lysis and binding solutions were UV-radiated for a minimum of 90 min. A volume of 1.0 mL lysis buffer was then added to each tube (0.5 M EDTA, pH 8.0 (Thermo Fisher Scientific, Waltham, MA, USA), 0.5% Tween-20 (Sigma-Aldrich, Saint Louis, MO, USA) and 20 mg/mL Proteinase K (Thermo Fisher Scientific) and briefly agitated in a FastPrep-24 instrument (MP Biomedicals, Santa Ana, CA, USA) for 10 s at 4.0 m/s. The samples were then incubated at 37°C overnight. The next morning the samples were agitated for the same duration and speed and then centrifuged 15 min at 16,000 g. The supernatants (3 x 0.5 mL) belonging to the same air filter were pooled in a 50 mL screw cap tube (Sarstedt, Newton, NC, USA). An additional 0.5 mL buffer (0.5 M EDTA, 0.5% Tween-20) was added to each filter punch, agitated for 30 s at 5.0 m/s and centrifuged for 15 min at 16,000 g. The supernatants were collected and added to the corresponding 50 mL tube. This procedure was repeated once more with a 30 s, 6 m/s agitation, and a 5 min centrifugation step.

To each 50 mL tube, 8.8 volumes of binding buffer were added (5M GuHCl, (≥ 99%, Sigma-Aldrich), 40% Isopropanol (Thermo Fisher Scientific), 90 mM NaAc (pH 5.2, Sigma-Aldrich), 0.05% Tween-20 (Sigma-Aldrich), Nuclease free water (Qiagen, Hilden, Germany), followed by 10 s vortexing. Using a QIAvac 24 Plus vacuum manifold (Qiagen), the solution was then passed through a Zymo-Spin IIICG column (Zymo Research, Irvine, CA, USA) mounted with conical reservoirs (Zymo Research). The column was washed once with 0.75 mL binding buffer and twice with 0.75 mL 80% Ethanol (Thermo Fisher Scientific). The column was dried by centrifugation for 2 min at 13,000 g. The column was then moved to a DNA LowBind tube (Sarstedt) and 60 μL EB buffer (Qiagen) was added to the column. The column was then left for 5 min before the DNA was eluted by centrifugation for 1 min at 13,000 g. The eluted DNA was further cleaned using DNeasy PowerClean pro (Qiagen) and repaired using NEBNext FFPE DNA Repair Mix (New England Biolabs) as per manufacturers’ protocol. The final DNA concentrations were measured using Qubit Fluorometric Quantification and the Qubit 1X dsDNA HS Assay Kit (Thermo Fisher Scientific).

### 2.2 Sequencing

Libraries were prepared from isolates with a minimum of ∼10 ng DNA at the Swedish National Genomics Infrastructure (SciLifeLab, SNP&SEQ, Uppsala) using the Thruplex DNA-Seq kit (Takara, Kusatsu, Shiga, Japan) with 8 PCR cycles according to the manufacturer’s protocol. Libraries were sequenced on Illumina NovaSeq 6000 S4 flow cells using 2 x 150 bp output (Illumina, San Diego, CA, USA). Read numbers for sequenced weeks are shown in fig. S4. Sequencing data are available through the NCBI Sequence Read Archive under project PRJNA808200. The files are named according to the following format Ki-YYYY-WW-RandID, where Ki is short for Kiruna station, YYYY and WW are the ISO year and week, respectively, and RandID is the randomized ID that determined the order of DNA extraction and sequencing.

**Fig. S4.**
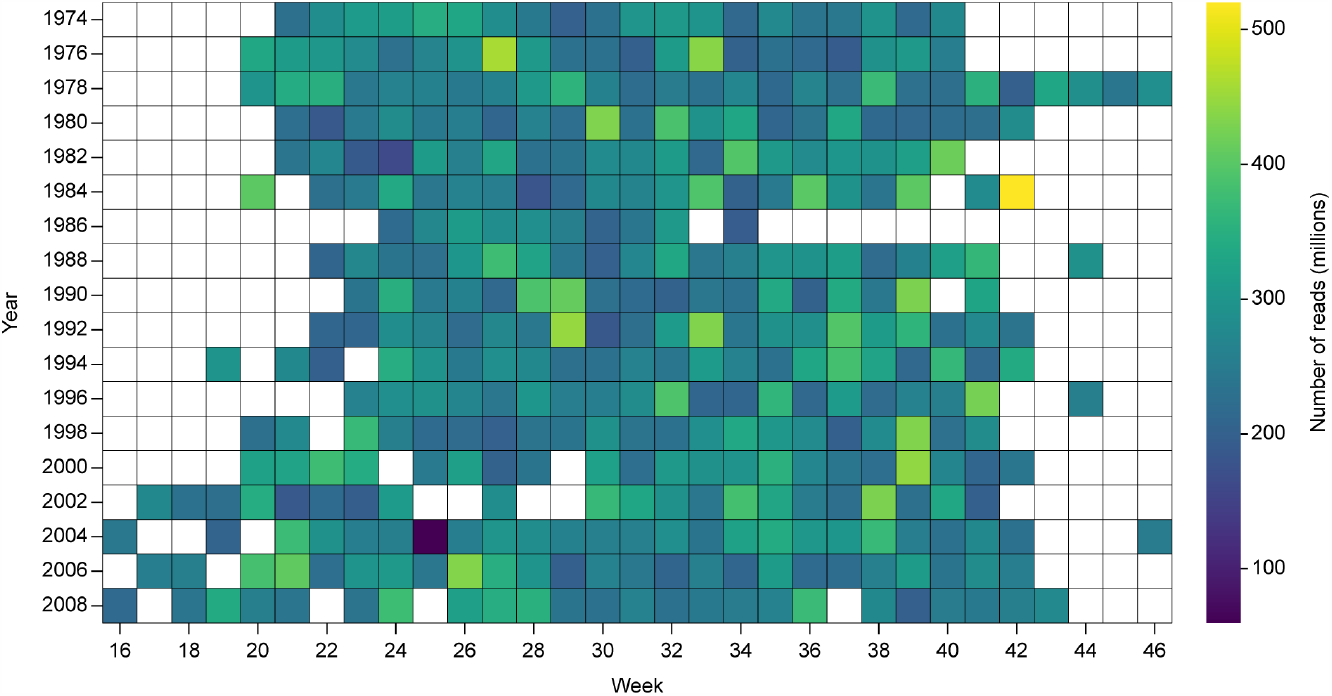
Number of paired-end reads sequenced from each weekly air filter. White cells indicate weeks without data. The consecutive run of missing data in the end of 1986 was due to air filters missing from the archive.

## 3. Bioinformatics pipeline

### 3.1 Read preprocessing and filtering

We first trimmed adapter sequences using Cutadapt *v*. 2.0 (*69*) and retained reads with length ≥ 50 bp. Air filters are replaced at the aerosol sampling station by hand. Therefore, we removed reads mapping to the human reference genome hg19 using BBMap *v*. 38.69 with the following parameters: `minid: 0.95 maxindel: 3 minhits: 2 bandwidthratio: 0.16 bandwidth: 12 qtrim: “rl” trimq: 10 quickmatch: “quickmatch” fast: “fast” untrim: “untrim”`. The proportion of human reads detected and removed from the weekly sequence data are displayed in fig. S5.

### 3.2 Taxonomic read classification

We used a version of Kraken 2 *v*. 2.0.8-beta (*70*) that we forked^1^ to report the number of minimizer hit groups in the standard output and StringMeUp,^2^ a post-processing python script developed in-house. StringMeUp allows reclassification of reads based on a user-specified confidence score stringency and/or minimum minimizer hit groups cutoff. It only requires the output from Kraken 2 and the taxonomy used to build the database. In short, StringMeUp processes each read by evaluating the confidence score at the currently assigned node. If the confidence score is less than the user-specified cutoff, the read is reclassified to the parent of the current node and the confidence score is recalculated as outlined in the manual of Kraken 2.^3^ This continues until the confidence score requirement is satisfied. If the current node is the root and the confidence score is less than the cutoff, the read is deemed unclassified.

**Fig. S5.**
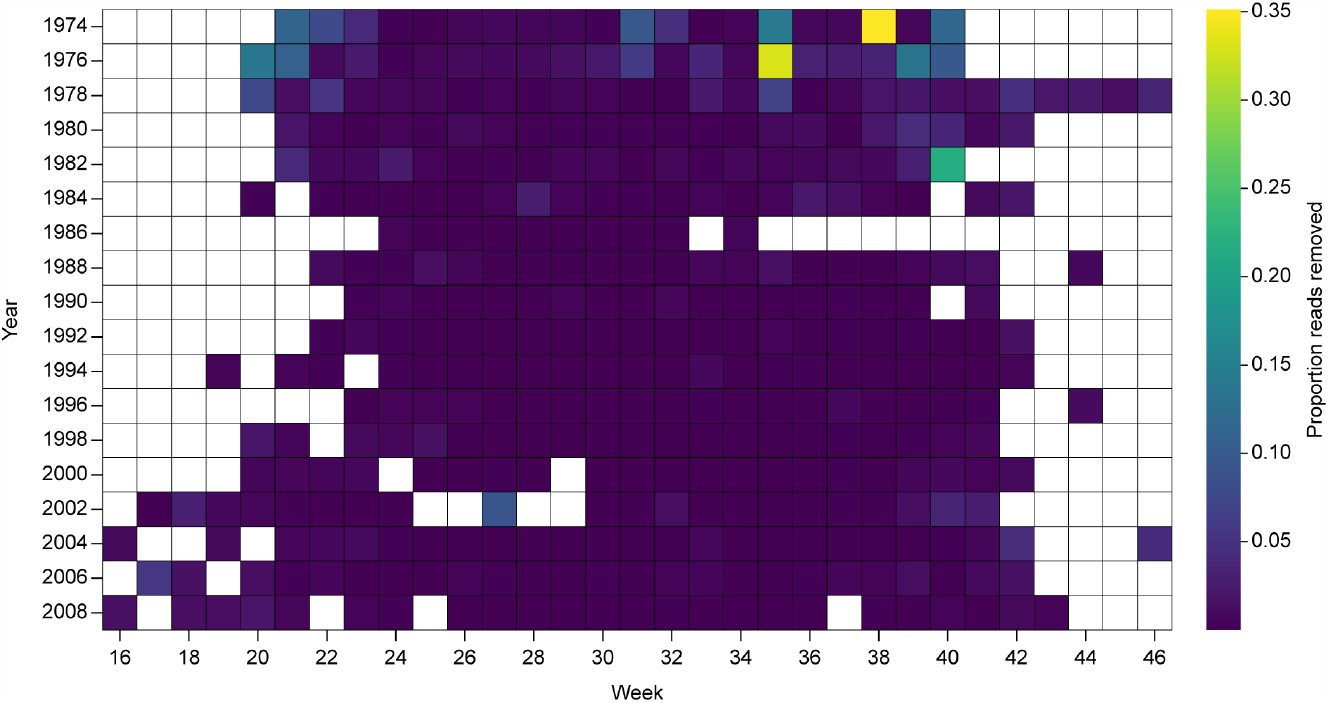
Human read removal. Proportion of paired-end reads from each week that mapped to the human reference genome and were removed prior to further analysis.

#### 3.2.1 Custom Kraken 2 database

Input data for the Kraken2 database comprised nucleotide sequences from the 1) NCBI non-redundant nucleotide (nt), 2) NCBI RefSeq genomic, and 3) GenBank whole genome shotgun (WGS) databases. The nt fasta file contained 256 GB of sequence data and was downloaded^4^ using the Kraken 2 command `--download-library`. The RefSeq genomic blast database was downloaded^5^ from the NCBI ftp,^6^ converted to a 1.6 terabyte (TB) fasta file using the NCBI blast+ package (*71*) application `blastdbcmd`, and staged for inclusion in the Kraken 2 database with the Kraken 2 command `--add-to-library`.

The WGS assemblies were selected in a multi-step process. First, a list of available WGS projects was acquired through the NCBI Sequence Set Browser^7^ and WGS projects (at the species rank) non-redundant with the nt or RefSeq genomic databases were identified. Projects with unannotated (UNA) or environmental (ENV) sequences or that lacked a biosample or taxonomic ID were excluded, leaving 13,731 projects from 4,809 unique species and 2.4 TB of sequence data. Fasta files were downloaded using `fastq-dump`, part of the SRA toolkit,^8^ and subsequently staged for inclusion in the Kraken 2 database in the same way as the RefSeq genomic fasta file.

Input for the Kraken 2 database build summed to 4.2 TB and included sequence data from 1,740,636 taxa from 89,168 named genera (data S2). From this, a 2.2 TB hash table (database) was built using 72 threads with a wall time of 75 hours. Minimizer and *k*-mer size settings were kept at their defaults.

#### 3.2.2 Kraken 2 classification and filtering with StringMeUp

Sequences from the 380 weeks were classified using the Kraken 2 database (section 3.2.1) using 72 threads with a mean wall time of 1.96 hours per sample. Classifications were made under minimal stringency settings, *i*.*e*., `--confidence 0` and `--minimum-hit-groups 1`. The reads were classified in this way so that StringMeUp could be applied on the output and stringency settings freely selected from a wide range. We found that 76,521 genera had at least one classified read under the minimum stringency threshold.

**Fig. S6.**
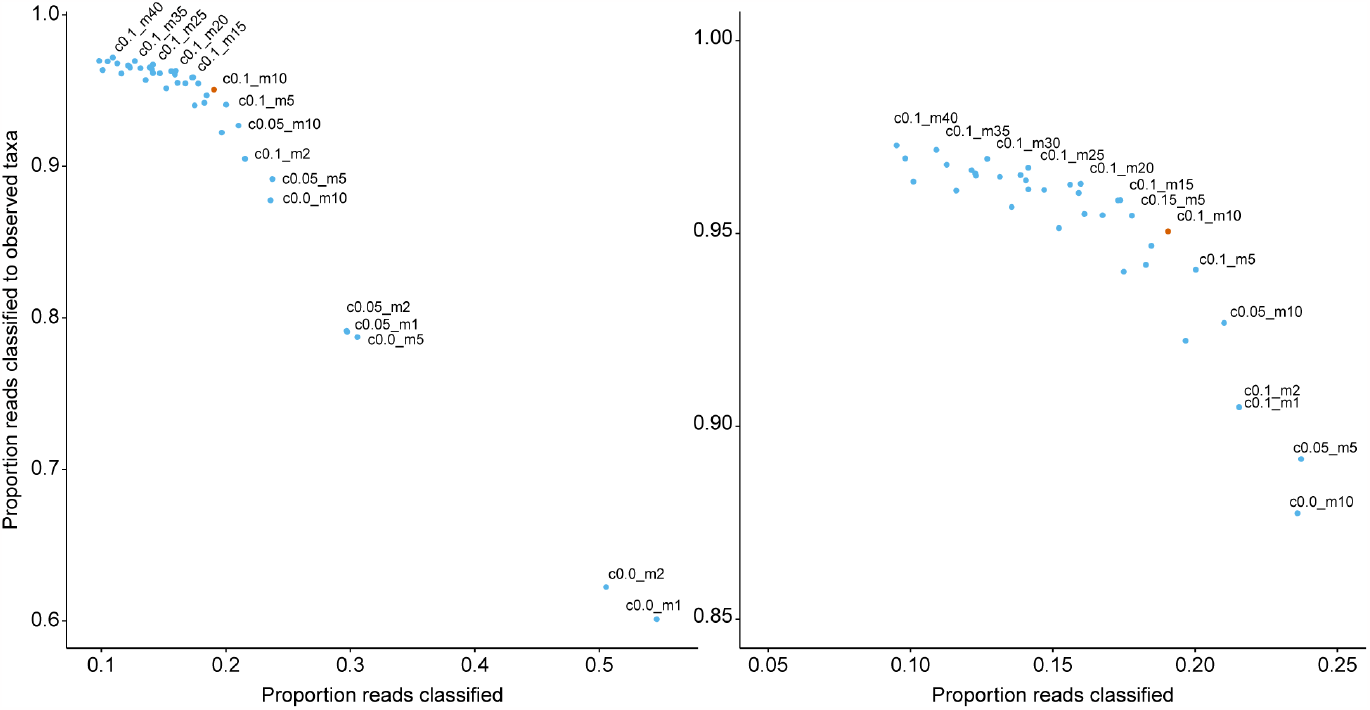
A larger fraction of reads are classified to taxa reported in Torne lappmark whereas classification success decreases with increasing stringency cutoffs in Kraken 2. Stringency was altered by varying the cutoffs for minimum confidence scores and hit groups (*e*.*g*., c0.1_m2 denotes a confidence score of 0.1 and 2 minimum number of hit groups). The combination we used is marked in red (c0.1_m10). Parameter combinations that have a lower proportion of reads assigned to observed taxa at a comparable level of total assigned reads are unlabeled. The left panel shows all tested combinations and the right shows a detailed view of the more stringent parameter settings.

The penultimate step in the read classification pipeline was to select confidence score and hit group threshold. We randomly selected two weeks from each year (*n* = 36), subset the reads assigned to the most abundant genera (> 25^th^ percentile), and then calculated the fraction of reads assigned to a taxonomic family observed in Torne lappmark^9^ out of all assigned reads over a grid of cutoff combinations. Taxa observations were retrieved from the Swedish Species Observation System database^10^ (*72*) (data S3). Confidence scores were evaluated at 0, 0.05, 0.1, 0.15, 0.2, 0.25, and 0.3. Minimum hit groups were evaluated at 1, 2, 5, and 10 for all confidence scores and at 15, 20, 25, 30, 35, and 40 for confidence scores 0, 0.05, and 0.1. The parameter space was extended until no improvement (in proportion of reads assigned to Torne lappmark taxa) was observed.

We considered a minimum confidence of 0.1 with a minimum of 10 hit groups to be a good trade-off between the fraction of reads assigned to taxa plausibly present near the aerosol sampling station and the total number of classified reads (fig. S6). Using this level of stringency, 40,034 genera had at least one classified read. More stringent cutoffs marginally increased the Torne lappmark fraction but the total number of classified reads continued to decrease almost linearly. A less stringent cutoff combined with the machine classifier in Section 4.3 may have increased the sensitivity of our assignments, but we preferred this more conservative approach for the ecosystem-level biodiversity analyses. Finally, we removed taxa that did not have > 10 classified reads in any of the weekly samples, leaving 15,672 genera.

## 4. Relative abundance transformations and detrending

### 4.1 Removal of zero inflated taxa and log-ratio transformations

Metagenomic datasets are a type of compositional data because the maximum number of reads is constrained by the sequencing instrument. In our dataset, classified reads for a given week comprise a *D*-part composition, where *D* is the number of genera. The sample space of a *D*-part composition is a subset of ℝ^*D*^ known as the simplex, 𝕊^*D*−1^ (*73*). Because a composition is only free to vary in 𝕊^D−1^, operations defined on ℝ^*D*^ are invalid. More simply, compositions cannot be added together or multiplied by a scalar and methods based on the covariance matrix cannot be expected to give sensible results. This challenge can be addressed by using the Aitchison geometry to define a Euclidean vector space on the simplex and using log-ratio transformations to express compositions as coordinates in ℝ with respect this geometry (*74*).

We performed most subsequent analyses on log-ratio transformed data, which requires addressing zero count data first. An observation of zero reads from an organism may be due to its true absence from the catchment area,but we assume zeros from regularly detected taxa are artifacts of limited, stochastic sampling. We removed 9,380 genera with zero counts in ≥ 2/3 of the weeks and imputed zeros for the remaining 6,292 using geometric Bayesian multiplicative replacement (*75*) as implemented by the `cmultRepl` function in the *R* package ‘zCompositions’ *v*. 1.4.0-1 (*76*). This method replaces zeros with estimates drawn from a multinomial distribution and preserves the sum and correlation structure of the composition.

The centered log-ratio (CLR) transformation maps a composition from the simplex 𝕊^*D*−1^ to the unconstrained space of ℝ^*D*^. The CLR transformation is an isometry, meaning the Euclidean distances between two parts of a composition in ℝ^*D*^ is equivalent to the Aitchison distance in 𝕊^*D*−1^. The CLR also provides a one-to-one transformation of all features, which makes interpretability easier but always results in singular covariance matrices.

A composition *x* ∈ 𝕊^*D*−1^ can be CLR transformed through:

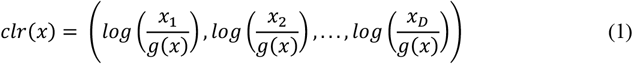

where *g*(*x*) is the geometric mean of the composition *x* and *D* is the number of parts in the composition *x*.

An alternative is the isometric log-ratio (ILR) transformation, which assigns coordinates in ℝ^*D*−1^ with respect to an orthonormal basis in 𝕊^*D*−1^. This transformation can be done according to the formulae:

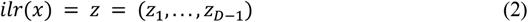

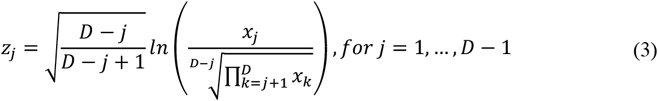

where *D* is the number of parts in the composition.

The ILR transformation is also an isometry and additionally matches the dimension of the simplex in ℝ and therefore does not result in singular covariance matrices. The tradeoff is the ILR transformation losses interpretability because matching the dimensionality of 𝕊 means there cannot be a one-to-one correspondence of the *D* compositional parts. Due to this, the ILR transformation is preferable to CLR when one wishes to analyze the composition as a whole, rather than a subset of components (*77*).

When using the ILR transformation, all information about *x*_1_ is contained the first coordinate *z*_1_. The same cannot be said about the other coordinates since *e*.*g*., *x*_2_ is used in the calculation of *z*_1_ and *z*_2_. Calculating the first coordinate for each *x* = (*x*_1_, …, *x*_*D*_) results in *D* pivot coordinate systems, which measures the relative dominance of each part in the composition (*74*). The pivot coordinate log-ratio (PLR) transformation is a pragmatic solution when univariate analysis or visualization of compositional parts is desired or necessary. PLR transformations were made with the *R* package ‘robCompositions’ *v*. 2.3.1 (*74, 78*).

### 4.2 Detrending

We identified three confounding factors that could bias eDNA abundance estimates: 1) a change in air filter manufacturer in 1996, 2) potentially more human contamination earlier in the time series (fig. S7), and 3) read length variation due to partial DNA degradation (fig. S7).

**Fig. S7.**
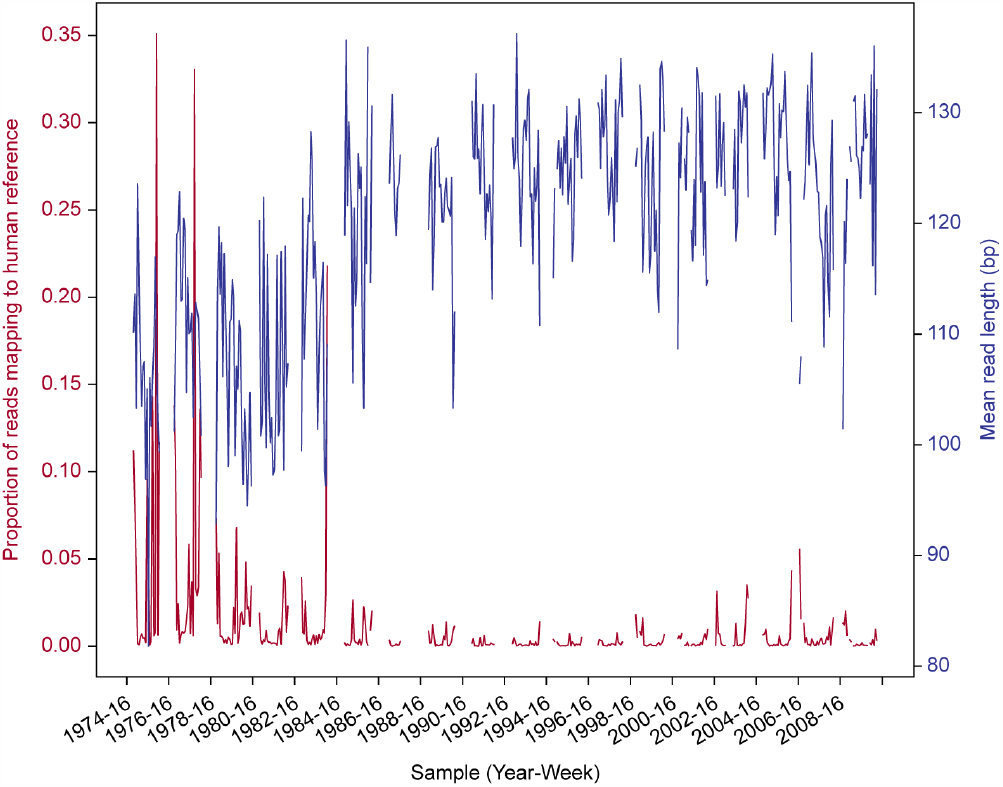
Proportion of human reads and mean read length. The proportion of reads that mapped to the human reference genome (red) and the mean read length in base pairs (blue).

We addressed read length variation by removing trends between genera abundances and their weekly mean read length. First, we applied the ILR transformation to both read lengths and relative abundances prior to detrending using the *R* package ‘compositions’ *v*. 2.0-6 (*79*). We modeled the weekly abundance of a given ILR component as a function of mean read length using generalized linear models (GLM). GLMs for each component were fit using the python module ‘statsmodels’ *v*. 0.11.1 (*80*) with the log, identity, and inverse link functions. The best fit was inferred using the Akaike information criterion (*81*). Weeks with a zero read count for a given component were not included in the models, leaving their imputed zero values unchanged. Sample means were re-added to the residuals, which were inversely transformed to relative abundances using the ‘compositions’ package. Redundancy analysis (RDA) was applied to the relative abundance matrix conditioned on air filter type and human read count proportion using the *R* package ‘vegan’ *v*. 2.6-4 (*82*) and the residuals were then used for subsequent analysis. For a comparison of the data before and after detrending, see fig. S8.

**Fig. S8.**
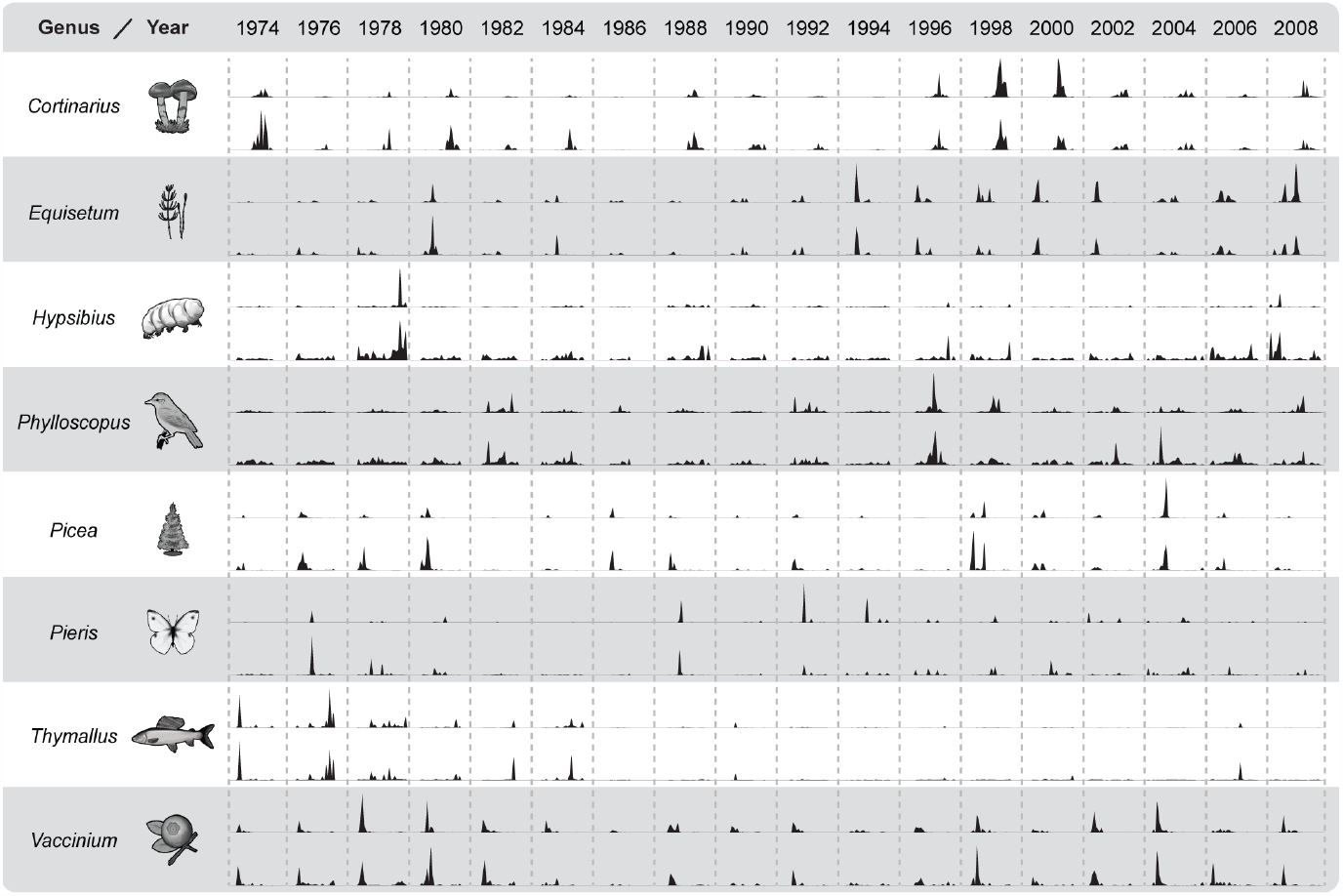
Comparison of relative abundances before and after detrending. For each genus two data tracks are shown. The top track shows relative abundances before detrending and the bottom after detrending. Abundances are scaled between 0 and 1.

### 4.3 Classification refinement with gradient boosting

The 6,292 genera putatively captured by the air filters included unlikely taxa such as the white rhinoceros (*Ceratotherium simum*). Besides initial misclassification due to read quality, redundant *k*-mers, and low sequence abundance, false positives may also arise from contaminants and other issues in the reference genomes and the unique computational burden imposed by any given database (*17*). To address this problem, we developed a gradient boosting machine (GBM) to distinguish between taxa likely to be true and false positives based on their classification metrics and abundance patterns throughout the time series.

#### 4.3.1 Feature engineering

Based on the known limitations of the classification pipeline (*17*) and the behavior of a few conspicuous false positives (see also Section 4.4), we hypothesized that false positive genera would have lower abundances; be detected rarely, or alternatively, with unusual consistency; occur more frequently in lineages with more sequence data and/or larger genomes, and have distinct per-read Kraken 2 classification quality metric profiles. We defined 31 features (parentheses correspond to column names in data S4) from these expectations and calculated them for each genus:

1) mean abundance (abundance_mean) and 2) its square (abundance_mean_squared),
3) median abundance (abundance_median) and 4) its square (abundance_median_squared),
5) 5^th^ percentile of weekly abundances (abundance_percentile_5th) and 6) its square (abundance_percentile_5th_squared),
7) 95^th^ percentile of weekly abundances (abundance_percentile_95th) and 8) its square (abundance_percentile_95th_squared),
9) number of weeks with relative abundance > 0 (weeks_present) and 10) its square (weeks_present_squared),
11) standard deviation of relative abundance (abundance_sd) and 12) its square (abundance_sd_squared),
13) abundance coefficient of variation (CV) and 14) its square (CV_squared),
15) number of minimizers per clade (minimizers_clade),
16) number of minimizers per taxon (minimizers_taxon),
17) total sequences per clade (total_sequence_clade),
18) total sequences per taxon (total_sequence_taxon),
19) ratio of mean abundance to number of clade minimizers (abundance_mean_minimizerC_ratio),
20) ratio of median abundance to number of clade minimizers (abundance_median_minimizerC_ratio),
21) ratio of mean abundance to number of taxon minimizers (abundance_mean_minimizerT_ratio),
22) ratio of median abundance to number of taxon minimizers (abundance_median_minimizerT_ratio),
23) Kraken 2 confidence score: total number of *k*-mers classified to a given genus divided by the total number of *k*-mers from the corresponding reads (confidence_original),
24) alternative confidence score: same as feature 22 but without unclassified *k*-mers in the denominator (confidence_classified),
25) other *k*-mers lineage ratio: number of *k*-mers classified to any node leading to the assigned genus divided by the total number of *k*-mers, excluding those classified to the genus (other_kmers_lineage_ratio),
26) other *k*-mers root ratio; the ratio of *k*-mers classified to the root node to the total number of *k-*mers, excluding those classified to the genus (other_kmers_root_ratio),
27) other *k*-mers classified ratio: the ratio of classified to unclassified *k*-mers, excluding those classified to the genus (other_kmers_classified_ratio),
28) other *k*-mers distance: the average taxonomic distance (number of intervening edges) between the nodes that the *k*-mers are classified to (other than the genus) and the genus that the corresponding reads are classified to (other_kmers_distance),
29) other *k*-mers distance lineage excluded: as in feature 27, but excluding *k*-mers classified to any rank in the lineage leading to the genus (other_kmers_distance_lineage_excluded),
30) total *k*-mers: sum of *k*-mers classified to the genus clade across the time series (total_kmers), and
31) number minimizer hit groups per *k*-mer: the sum of minimizer hit groups from reads classified to a genus divided by the total number of *k*-mers (mhg_per_kmer).

We also considered the possibility that one or more weeks could be enriched for false positives by including the weekly abundance of each genus as features. Finally, we one-hot encoded kingdom-rank assignments to allow these features to differ in their utility and probability distributions. All features were calculated from PLR transformed data (Section 4.1).

#### 4.3.2 Training data acquisition

As we lacked labeled (*i*.*e*., empirically known) training data, we used species occurrence records to create two groups that we expect to be enriched for true and false positive taxa, respectively. As pseudopostives, we used genera registered in the Swedish Species Observation System^11^ with > 3 observations reported from ≤ 40 km of the aerosol sampling station between 1974-2008 (*83*). We also included humans, dogs, *Aedes*, and 33 bacterial genera identified in soil and water samples from a similar ecosystem^12^ as pseudopositives, yielding 317 in total. For pseudonegative taxa, we identified 379 taxa that 1) have no reported occurrences in the Global Biodiversity Information Facility online database (GBIF) within 5,000 km of the aerosol sampling station (*84*), and 2) are not closely related to any European taxa lacking a reference genome. For example, Glossinidae, containing the *Glossina* tsetse flies, is in the same superfamily as the Hippoboscidae, which occur in Europe and lack a representative genome, so *Glossina* was not considered a pseudonegative genus. These criteria presumably exclude many actual false positives (*i*.*e*., where the classification does not result from shared ancestry) from the training data, but we wanted to allow genera poorly represented in the reference database to be captured at higher taxonomic ranks (see also Section 4.4). Prior to model training, we randomly selected and set aside 15% (*n* = 91) of the pseudopresences and absences as test data. The full list of pseudolabeled taxa and their feature data is provided as data S4 and their taxonomic composition is summarized in table S2.

**Table S2.** Taxonomic composition of pseudolabeled data. Taxa are divided by kingdom into positive and negative and test and training fractions. Orders with more than 15 labeled taxa are shown; the remaining taxa in each kingdom are summed as ‘others’.

| taxon | training |  | test |  | total |  |
| --- | --- | --- | --- | --- | --- | --- |
|  | neg. | pos. | neg. | pos. | neg. | pos. |
| *Bacteria* | 0 | 29 | 0 | 4 | 0 | 33 |
| *Metazoa* | 212 | 70 | 35 | 12 | 247 | 82 |
| Mammalia | 110 | 11 | 18 | 0 | 128 | 11 |
| Aves | 27 | 31 | 8 | 4 | 35 | 35 |
| Insecta | 4 | 25 | 0 | 6 | 4 | 31 |
| Actinopteri | 40 | 0 | 6 | 1 | 46 | 1 |
| others | 31 | 3 | 3 | 1 | 34 | 4 |
| *Viridiplantae* | 90 | 99 | 15 | 16 | 105 | 115 |
| Magnoliopsida | 33 | 60 | 5 | 10 | 38 | 70 |
| Pinopsida | 26 | 3 | 4 | 0 | 30 | 3 |
| Polypodiopsida | 24 | 7 | 6 | 0 | 30 | 7 |
| Bryopsida | 0 | 16 | 0 | 3 | 0 | 19 |
| others | 7 | 13 | 0 | 3 | 7 | 16 |
| *Fungi* | 24 | 81 | 3 | 6 | 27 | 87 |
| Agaricomycetes | 16 | 52 | 1 | 2 | 17 | 54 |
| Lecanoromycetes | 0 | 14 | 0 | 3 | 0 | 17 |
| others | 8 | 15 | 2 | 1 | 10 | 16 |
| **total** | **326** | **279** | **53** | **38** | **379** | **317** |

#### 4.3.3 Parameter tuning and classification

We trained the GBM using the *R* interface for xgboost *v*. 1.7.5.1 (*85*). We iteratively performed grid searches with 5-fold cross validation over a total of 6,561 hyperparameter combinations to identify a set approaching the smallest binary classification error rate. First, we fixed the learning rate (eta) to 0.3 and explored regularization and tree-specific parameters over the grid:

~~~
max_depth = c(1, 3, 5, 7, 9),
min_child_weight = c(1, 3, 5, 7, 9),
gamma = c(0.0, 0.01, 0.1, 0.3, 0.5, 1.0),
subsample = c(0.4, 0.6, 0.8),
colsample_bytree = c(0.4, 0.6, 0.8),
reg_alpha = c(1e-5, 1e-2, 0.1, 1, 100),
reg_lambda = c(1.0, 1.5, 2.0, 3.0, 4.5).
~~~

We defined successively narrower ranges over six tuning rounds and, in the final round of tuning, tested `eta = c(0.1, 0.15, 0.2, 0.25, 0.3)` with the remaining parameters fixed. The final trained model used: `eta = 0.3, max_depth = 5, min_child_weight = 2, subsample = 0.7, colsample_bytree = 0.4, reg_alpha = 1e-05, gamma = 0.3, reg_lamba = 1.5.`

**Table S3.** Gradient boosting machine (GBM) classification performance. False discovery rate (FDR), precision, and recall are reported for the n = 91 test dataset using predictive probabilities from 0.50 to 0.95 as the cutoff for a positive classification. ‘#negative’ and ‘#positive’ denote the number of genera below or above a given cutoff, respectively, out of the 6,292 genera dataset.

| cutoff | FDR | precision | recall | #negative | #positive |
| --- | --- | --- | --- | --- | --- |
| 0.50 | 0.09 | 0.89 | 0.74 | 2,830 | 3,462 |
| 0.55 | 0.08 | 0.91 | 0.74 | 2,941 | 3,351 |
| 0.60 | 0.08 | 0.91 | 0.74 | 3,083 | 3,209 |
| 0.65 | 0.08 | 0.91 | 0.74 | 3,225 | 3,067 |
| 0.70 | 0.06 | 0.93 | 0.74 | 3,369 | 2,923 |
| 0.75 | 0.04 | 0.95 | 0.71 | 3,553 | 2,739 |
| 0.80 | 0.04 | 0.95 | 0.71 | 3,737 | 2,555 |
| 0.85 | 0.04 | 0.94 | 0.61 | 3,960 | 2,332 |
| 0.90 | 0.02 | 0.96 | 0.50 | 4,267 | 2,025 |
| 0.95 | 0.00 | 1.00 | 0.47 | 4,795 | 1,497 |

We compared the false discovery rate (FDR), precision, and recall for the test data over a range of predicted classification probabilities (table S3). We emphasize that these are based on estimated labels and do not necessarily indicate the error rates of the full dataset. Nevertheless, the key features identified by the trained GBM are mostly derived from the *k*-mer classification patterns, a result expected only if the pseudolabeled training are enriched for real positive and negatives. Four features comprised 45% of the binary classification error improvement: the original Kraken 2 confidence score (25%; feature 22 in Section 4.3.1); other *k*-mers distance (13%; feature 27), relative abundance standard deviation (4%; feature 10), and the classified confidence score (3%; feature 23). Pseudonegative genera tended to have a larger other *k*-mers distance, a smaller standard deviation, and lower confidence scores than pseudopostives (fig. S9). This suggests false positive genera are likely to show limited variation in abundance over the time series and that reads with *k*-mers assigned to false positives tend to also contain *k-*mers assigned to taxonomically-distant clades. In particular, *k*-mer distances > 20 result if a read contains *k*-mers classified to both eukaryotes and prokaryotes, which can occur from reference genome contamination (Section 4.4).

**Fig. S9.**
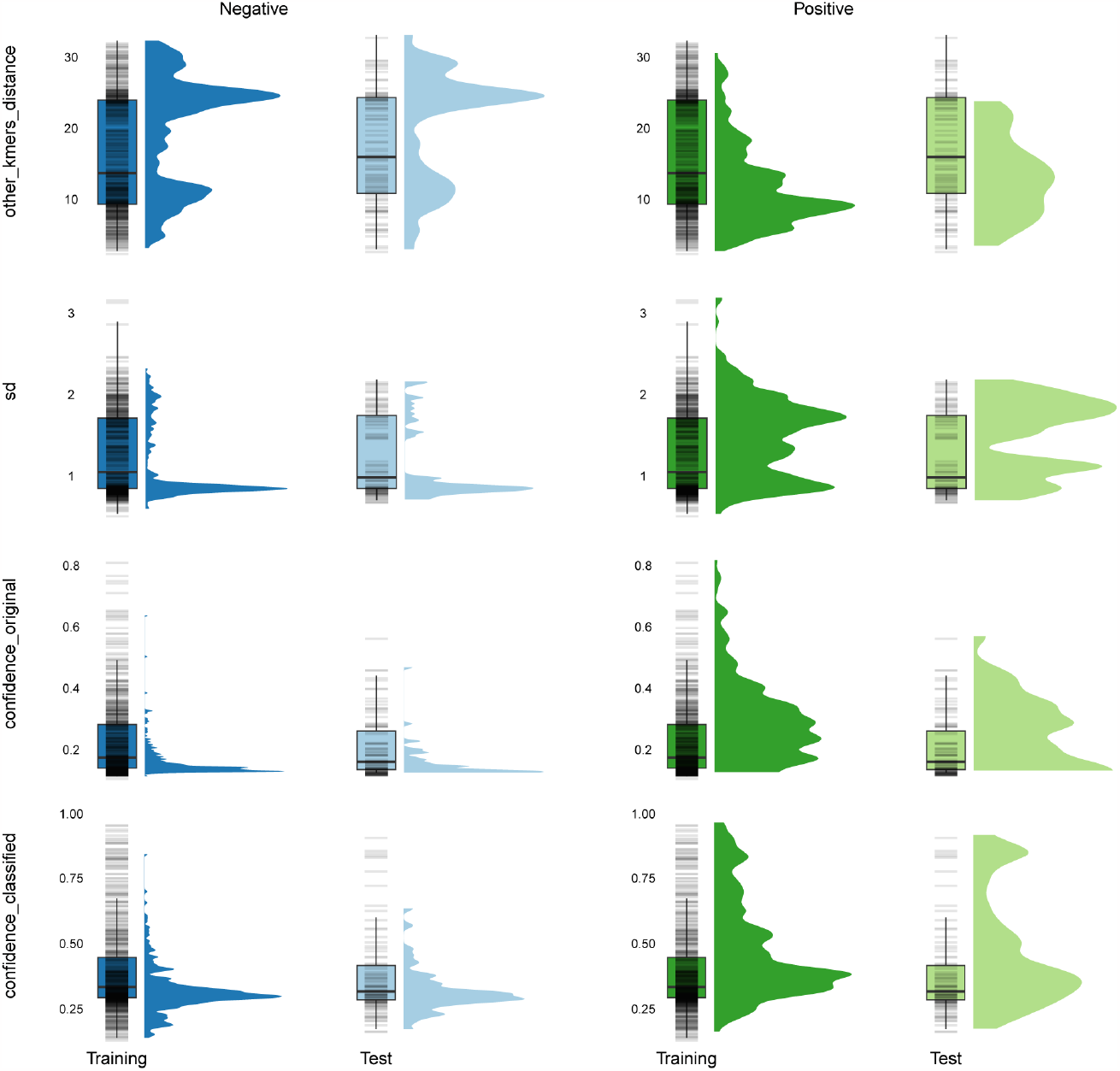
Distributions of the four most influential features in the gradient boosting machine classification model. Each row corresponds to a feature. The first two columns (blues) show values for the pseudolabeled negative genera and the last two columns (greens) show pseudolabeled positive genera. Training data are shown in a darker shade and test data in a lighter shade.

For the final classification of the 6,292 genera dataset, we selected the 0.75 probability threshold, which classified 2,739 as true occurrences (weekly relative proportions of these taxa are given in data S5). Given the taxonomic composition of the pseudolabeled training data (table S2), we expect genera-rank classifications to be most accurate for mammals, birds, and fish, followed by common seed plants and agaricomycote fungi. Genera-rank assignments for insects and microbial taxa are probably the least accurate but we demonstrate a method in Section 4.4 for determining if questionable assignments result from shared ancestry with the assigned genus.

### 4.4 Read mapping analysis of classified taxa

Accurate taxonomic classification highly depends on the reference sequences in the database library. Organisms lacking a reference can be misclassified *e*.*g*., to close relatives or contaminated reference genomes. After Kraken 2 classification we found reads assigned to organisms that are vanishingly unlikely to be present near the aerosol sampling station, such as the white rhinoceros (*Ceratotherium simum*). To understand the source of this signal, we mapped reads classified by Kraken 2 to white rhinoceros back to northern white rhinoceros’ genome.^13^ Reads from week 37 in 1996 were selected for this analysis due to high number classified to white rhinoceros. Reads were aligned using Hisat2 *v*. 2.2.1 (*86*) using default parameters. Reads from the air filter mapped to only 213 out of the 942,426 contigs in the draft assembly. We blasted (blast *v*. 2.10.1+) 15 of the contigs with most hits against the nt database using the following parameters (unspecified parameters kept as default):

~~~
    -task megablast
    -db nt_v5
    -outfmt “6 qseqid staxids bitscore std sscinames sskingdoms stitle”
    -num_threads 10
    -max_target_seqs 10
    -evalue 1e-25
    -max_hsps
~~~

The blast matches showed > 80% identity with sequences from *Pseudomonas* species. From this, we conclude the signal from white rhinoceros is a false positive caused by reference genome contamination.

After performing genus-level classification refinement (Section 4.3), we detected two potential misclassifications among the 100 most abundant taxa: the forest tree *Larix* (Pinales: Pinaceae) and an insect endemic to Antarctica, *Belgica* (Diptera: Chironomidae). Reads assigned to *Larix* are surprisingly abundant (9% of reads assigned to positive classified taxa), given that the nearest natural populations are located *ca*. 1,000 km east in Arkhangelsk oblast, Russia or 2,000 km south in the northern Carpathians. This is comparable to *Picea* (9% of reads), which is the dominant tree along with *Pinus* (30%) in the region. Interestingly, we found an almost perfect correlation between the PLR coordinates of *Larix* and *Pinus* (fig. S10A). *Larix* flowers *ca*. 2 months earlier than *Pinus* in central Europe (*87*) and about month earlier in Siberia (*88*), which suggests they would also differ in phenology if present together in the aerosol station’s catchment area.

We mapped reads classified to *Pinus* and *Larix* back to their reference sequences included in our Kraken 2 database (masked for low-complexity sequences). We extracted reads from the week with the highest abundance in each year for *Larix*: 1974:25, 1976:27, 1978:26, 1980:26, 1982:28, 1984:23, 1986:26, 1988:21, 1990:27, 1992:25, 1994:27, 1996:28, 2000:28, 2002:23, and 2004:28. We only mapped reads from weeks 1980:26, 1990:27, and 2004:28 for *Pinus* due to the extremely high number of *Pinus*-classified reads in our dataset (> 10^8^ PE reads during flowering weeks). Extracted reads were mapped back using BBMap *v*. 38.98 with the following parameters: `pairedonly = t ambiguous = best killbadpairs = f minid = 0.97` (other parameters set as default). For a true positive signal, we expect aligned reads to be randomly distributed across the non-repetitive parts of the genome. Thus, we expect a positive relationship between the number of reads aligned and the contig length. To compare the last between *Larix* and *Pinus*, we used the mapping results of the 1,000 longest contigs from each genus. We see the expected positive relationship between the aligned reads and contig length for *Pinus* (fig. S10B) but not for *Larix* (fig. S10C). Our results suggest *Larix* is a false positive, unlike the *Pinus* signal, potentially driven by cross-classification of *Pinus* reads.

We investigated *Belgica* using the same false-true positive reasoning that we used for *Larix* and *Pinus*. We mapped *Belgica*-classified reads from weeks 1974:26, 1976:39, 1978:27, 1980:23, 1982:27, 1984:26, 1986:28, 1988:30, 1990:24, 1992:27, 1994:31, 1996:31, 1998:32, 2000:30, 2002:27, 2004:34, 2006:23, and 2008:35 back to its reference sequences in the Kraken 2 database using the same method as for *Larix* and *Pinus*. For *Belgica*, we found a positive relationship between the number of aligned reads and contig length, as in *Pinus* (fig. S10D). From this, we conclude that the *Belgica* signal most likely originates from a relative absent from the reference database. From the perspective of our GBM classifier, *Belgica* would then be correctly classified as a positive occurrence, even though the genera-rank assignment is extremely unlikely to be correct.

**Fig. S10.**
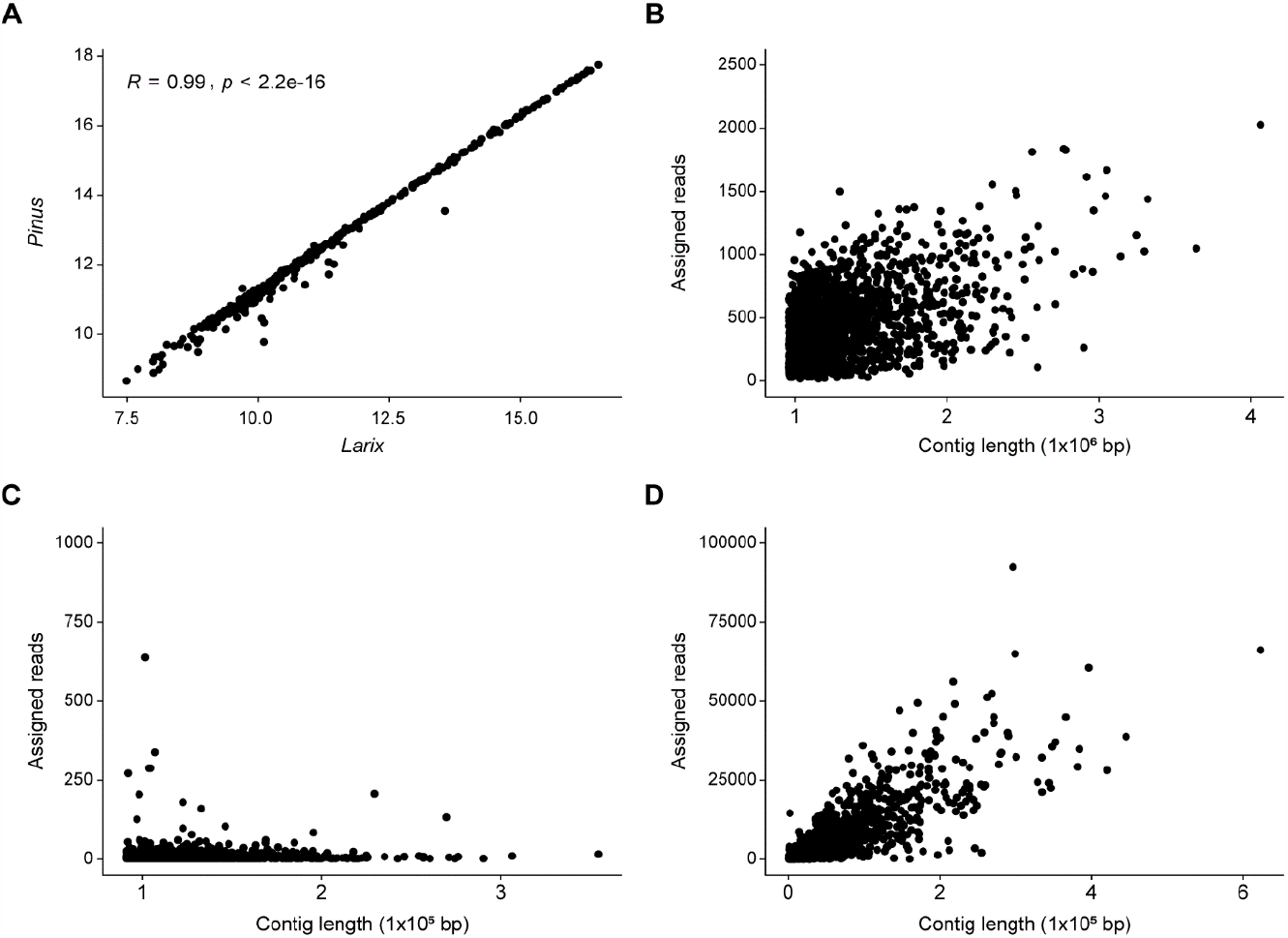
Correlation of *Pinus* and *Larix* abundances and relationship between the number of mapped reads and contig length for *Pinus, Larix* and *Belgica*. **A**) Correlation between *Pinus* and *Larix* PLR coordinates (ρ = 0.99, *p* < 0.001). Number of mapped reads per contig *vs*. contig length for **B**) *Pinus*, **C**) *Larix* and **D**) *Belgica*.

## 5. Dimensionality reduction and clustering

### 5.1 Taxa-based clustering and ordination

Standard measures of correlation and distance are inappropriate for compositional data due to their constrained covariance structure. Therefore, we employed an analogue of dissimilarity calculated from the pairwise variance between CLR transformed abundances (*31, 89*):

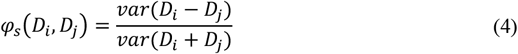

We then performed hierarchical clustering of the 2,739 genera based on their pairwise *φ*_*s*_ using Ward’s method. This method was considered one of the most feasible options based on a benchmarking routine that evaluated various clustering methods, including Gaussian mixture models (GMM), DB-SCAN, k-Means, and hierarchical clustering. The clustering performance was assessed through a combination of silhouette (*90*) and Calinski-Harabasz (*91*) indices. The cluster membership of each genus at *k* = 17 is included in data S5 and the taxonomic composition of each cluster is summarized in data S6.

## 6. Diversity metrics

Our data can be thought of as a metacommunity, where the sequences from each weekly filter sample is a subcommunity. This is illustrated in the matrix *P* below, where each column contains the sequences from week *w*_1_, *w*_2_, …, *w*_*N*_ and each row contains the relative abundance of genus *p*_1_, *p*_2_, …, *p*_*S*_:

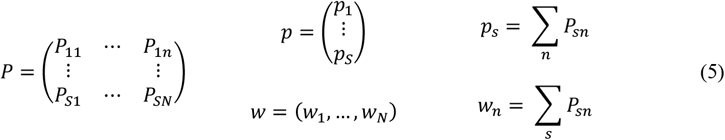

where ∑_*n*_ *w*_*n*_ = ∑_*S*_ *p*_*S*_ = ∑_*s,n*_ *P*_*sn*_ = 1. This is equivalent to considering *P*_*SN*_ as a probability distribution ∈ {1, …, *S*} × {1, …, *N*} with marginal distributions *p* and *w*. For our data, *S* = 2,739 positive-classified genera (Section 4.3) and *N* = 378 sequence compositions from calendar weeks 21-41 in even-numbered years from 1974-2008.

We partitioned the diversity observed in week *l* into alpha (α), beta (β), and gamma (γ) diversity components following the framework of (*41*) and (*42*). As in the Hill numbers and Shannon entropy, α-diversity here quantifies the evenness, or average rarity, of *P*_.*n*_ independently from the rest of the time series. In contrast, β- and γ- diversity relate *P*_.*n*_ to *p*, the vector of marginal relative abundances. β-diversity scales *p* by *w*_*n*_, the size of the community in week *l* to measure the distinctiveness of the composition. Scaling by *w*_*n*_ allows comparison of changes in compositional uniqueness that are conditionally independent of α-diversity. γ-diversity measures the average rarity of taxa in week *l* with respect to the entire metacommunity, that is, γ = β + α.

α-diversity is Hill diversity and equal to the exponential of Shannon entropy when *q* = 1:

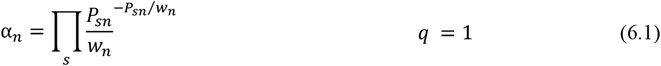

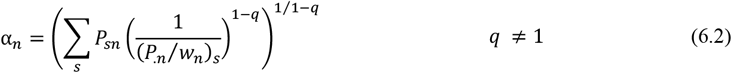

Larger values of *q* increasingly emphasize dominant over rare taxa; ^*q*=0^*α* is taxon richness and ^*q*=2^*α* is also known as Simpson’s concentration index. Higher α-diversity (for *q* > 0) indicates a more even abundance distribution, that is, a larger number of effective taxa. α-diversity obtains its maximum α = *S* if all 1, …, *S* taxa are present in equal relative abundances.

β-diversity is the exponential of Rényi’s relative entropy and equal to the exponential of Kullback-Leilber divergence for *q* = 1:

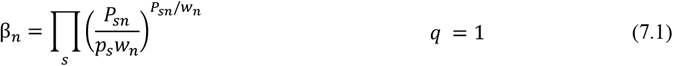

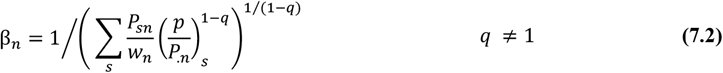

β-diversity measures the distinctiveness of the genera abundance distribution of the *n*^th^ week relative to the entire metacommunity. β-diversity is 1 if the composition in a week is identical to the whole metacommunity (*i*.*e*., perfectly representative) and increases as genera are more overrepresented in week *n* relative to *p*_*s*_*w*_*n*_ to a maximum of β_*n*_ = 1/*w*_*n*_.

γ-diversity is the exponential of Rényi’s cross entropy:

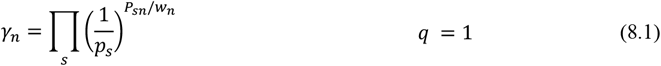

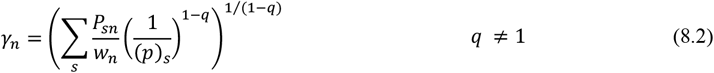

This measures the average rarity of genera in week *n* relative to the metacommunity. This means that if the composition of a week is identical to the marginal distribution *p*, its γ-diversity would equal the α-diversity of the metacommunity. γ-diversity increases with evenness, as in α-diversity, and as genera are more common in week *n* compared to their overall rarity, up to a maximum of *γ*_*n*_ = *S*/w _*n*_.

### 6.1 Per-taxon γ-diversity contributions

We tested for significant differences in the weekly γ-diversity contributions from each genus, *i*.*e*., the multiplicand in Equation 8.1, in matched calendar weeks between the early and late years of the time series using the two-sided Wilcoxon rank sum test. We initially assessed the sensitivity of the results to the years used as the ‘early’ and ‘late’ periods using comparisons between ‘74-’80 vs. ‘02-’08, ‘74-’82 *vs*. ‘00-’08, ‘74-’84 *vs*. ‘98-’08, ‘74-’86 *vs*. ‘96-’08, and ‘74-’88 *vs*. ‘94-’08. We avoided comparisons including ‘90 and ‘92 because these years correspond to the temporary peak in *Pinus* abundance and the lowest γ-diversity. With the exception of *Picea*, we found no difference in the significance of Benjamini-Hochberg adjusted *p*-values (FDR°=°0.05) or the direction of change for the genera with the largest differences in γ-diversity contributions (those in Fig. 3C in the main text). *Picea* changed both signs and significance depending on the weeks used in the comparison, likely because pollen production is irregular in Norway spruce. We therefore used ‘74-’88 *vs*. ‘94-’08 for the analysis. The median per-genus difference in γ-diversity contribution, 95% confidence intervals, and Benjamini-Hochberg adjusted *p*-values are given in data S7.

## 7. Climatic variables

### 7.1 Data sources and construction

We used observations from a weather station^14^ located *ca*. 3 km from the aerosol sampling station (*92*) and 1/24° gridded daily estimates (*93*) to construct 24 base variables capturing changes in the mean, variance, skewness, and kurtosis of local precipitation and temperature. Fifteen follow the ETCCDI climate extreme indices (*94*), including inhomogeneity adjustments (*95*), but we estimate their values over multiple rolling intervals. We derived 20 variables describing water and energy available for primary production from the monthly values in the 1/24° TerraClimate dataset (*96*). Given the frequency of the eDNA samples, we disaggregated the Terraclim variables to weekly intervals using cubic spline interpolation such that monthly means (or sums, if applicable) remained unchanged. Similarly, we interpolated weekly values from the monthly indices of the North Atlantic (*97*) and Atlantic Multidecadal (*98*) oscillations, which influence regional temperature and precipitation. Weekly values for the Arctic oscillation were calculated from daily indices (*99*). All 56 base variables and their data sources are summarized in table S4.

The duration of exposure to thermal and moisture variability can modulate vital rates and phenological patterns. For example, accumulated temperature is a key signal of bud burst and insect emergence and the balance between duration and intensity influences the ability of organisms to acclimate to stressful conditions (*100*). To incorporate some of this complexity into our models, we applied summary statistics to each base variable over rolling windows covering up to the previous 78 weeks. Intervals were selected to reflect local seasonal patterns between 1961 and 2009: four and eight weeks cover the period between the first (last) days consistently > 0°C and ≥ 5°C (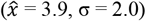, σ = 2.0); 13, 17, and 26 weeks connect the current week to conditions during the prior spring thaw (week number 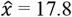, σ = 1.5), snow melt (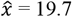, σ = 1.1), and start of the 5°C growing season (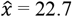, σ = 1.4); and the 52 and 78 windows include the influence of the prior growing and dormant season, with the latter including the two previous dormant seasons. Additionally, we included one- and two-week windows for daily counts of 0°C temperatures and mean daily temperature and precipitation. For disaggregated variables, we considered standard deviations over ≥ 8-week intervals due to their original monthly resolution. Note that observations are equally weighted within windows and do not incorporate time-lagged effects *per se* but values of TNN_52,78_ and TXN_52,78_ are determined by the previous year’s winter temperatures and TXX_52,78_ and TNX_52,78_ by summer.

**Table S4.** Summary of climatic covariables.

| name | base description | $f(x)$ | windows (weeks) |
| --- | --- | --- | --- |
| AET<sup>a</sup> | Actual Evapotranspiration; weekly total water extracted from plants and soil | $\bar{x}$ | 4,8,13,17,26,52,78 |
| | | $\sigma$ | 8,13,17,26,52,78 |
| AMOI<sup>b</sup> | Atlantic Multidecadal Oscillation Index; weekly mean | $\bar{x}$ | 4,8,13,17,26,52,78 |
| | | $\sigma$ | 8,13,17,26,52,78 |
| AMOI.LP<sup>b</sup> | 10-yr low-pass Atlantic Multidecadal Oscillation Index; weekly mean | $\bar{x}$ | 4,8,13,17,26,52,78 |
| | | $\sigma$ | 8,13,17,26,52,78 |
| AOI<sup>c</sup> | Arctic Oscillation Index: daily Hurrell station-based value | $\bar{x}$ | 4,8,13,17,26,52,78 |
| | | $\sigma$ | 8,13,17,26,52,78 |
| DS<sup>d</sup> | Dry spells; $\geq 6$ consecutive days with $< 1$ mm precipitation | $\Sigma$ | 4,8,13,17,26,52,78 |
| CSD<sup>e</sup> | Cold spell duration; $\geq 6$ consecutive days where $T_{\text{MIN}} < 10^{\text{th}}$ percentile<sup>†</sup> | $\Sigma$ | 4,8,13,17,26,52,78 |
| WS<sup>d</sup> | Wet spells; $\geq 6$ consecutive days with $\geq 1$ mm precipitation | $\Sigma$ | 4,8,13,17,26,52,78 |
| deficit<sup>a</sup> | Deficit: difference between weekly PET and AET totals | $\bar{x}$ | 4,8,13,17,26,52,78 |
| | | $\sigma$ | 8,13,17,26,52,78 |
| DTR<sup>e</sup> | Diurnal temperature range; difference between daily $T_{\text{MIN}}$ and $T_{\text{MAX}}$<sup>†</sup> | $\bar{x}$ | 4,8,13,17,26,52,78 |
| FCF<sup>e</sup> | Frost change frequency; days where $T_{\text{MIN}} < 0^{\circ}\text{C}$ and $T_{\text{MAX}} > 0^{\circ}\text{C}$ | % | 1,2,4,8,13,17,26,52,78 |
| FD<sup>e</sup> | Frost days; $T_{\text{MIN}} < 0^{\circ}\text{C}$<sup>†</sup> | $\Sigma$ | 1,2,4,8,13,17,26,52,78 |
| ID<sup>e</sup> | Ice days; $T_{\text{MAX}} < 0^{\circ}\text{C}$<sup>†</sup> | $\Sigma$ | 1,2,4,8,13,17,26,52,78 |
| NAOI<sup>f</sup> | North Atlantic Oscillation Index; weekly mean Hurrell station-based value | $\bar{x}$ | 4,8,13,17,26,52,78 |
| | | $\sigma$ | 8,13,17,26,52,78 |
| PD<sup>a</sup> | Potential deficit; difference between weekly precipitation and PET totals | $\bar{x}$ | 4,8,13,17,26,52,78 |
| | | $\sigma$ | 8,13,17,26,52,78 |
| PDSI<sup>a</sup> | Palmer Drought Severity Index; weekly mean | $\bar{x}$ | 4,8,13,17,26,52,78 |
| | | $\sigma$ | 8,13,17,26,52,78 |
| PET<sup>a</sup> | Potential evapotranspiration; weekly total Penman-Montieth reference evapotranspiration | $\bar{x}$ | 4,8,13,17,26,52,78 |
| | | $\sigma$ | 8,13,17,26,52,78 |
| precip<sup>d</sup> | Total daily precipitation | $\bar{x}$<br>$\sigma$ | 1,2,4,8,13,17,26,52,78 |
| pressure<sup>e</sup> | Daily mean atmospheric pressure | $\bar{x}$<br>$\sigma$ | 4,8,13,17,26,52,78<br>8,13,17,26,52,78 |
| radiation<sup>a</sup> | Weekly total downward surface shortwave solar radiation | $\bar{x}$<br>$\sigma$ | 4,8,13,17,26,52,78<br>8,13,17,26,52,78 |
| RM10<sup>d</sup> | Days with $\geq 10$ mm precipitation<sup>†</sup> | $\Sigma$ | 4,8,13,17,26,52,78 |
| runoff<sup>a</sup> | Weekly total precipitation and snowmelt exceeding PET and soil recharge | $\bar{x}$<br>$\sigma$ | 4,8,13,17,26,52,78<br>8,13,17,26,52,78 |
| RX1day<sup>d</sup> | Maximum 1-day precipitation<sup>†</sup> | $\Sigma$ | 4,8,13,17,26,52,78 |
| soil<sup>a</sup> | Weekly total soil column moisture | $\bar{x}$<br>$\sigma$ | 4,8,13,17,26,52,78<br>8,13,17,26,52,78 |
| SWE<sup>a</sup> | Snow water equivalent; amount of liquid water in snow pack | $\bar{x}$<br>$\sigma$ | 4,8,13,17,26,52,78<br>8,13,17,26,52,78 |
| TAVG<sup>d</sup> | Daily mean temperature | $\bar{x}$<br>$\sigma$ | 1,2,4,8,13,17,26,52,78 |
| TMAX<sup>e</sup> | Daily maximum temperature | $\bar{x}$ | 1,2,4,8,13,17,26,52,78 |
| TMIN<sup>e</sup> | Daily minimum temperature | $\bar{x}$ | 1,2,4,8,13,17,26,52,78 |
| TN10p<sup>e</sup> | Cool nights; days where $T_{\text{MIN}} < 10^{\text{th}}$ percentile<sup>†</sup> | % | 4,8,13,17,26,52,78 |
| TN90p<sup>e</sup> | Warm nights; days where $T_{\text{MIN}} > 90^{\text{th}}$ percentile<sup>†</sup> | % | 4,8,13,17,26,52,78 |
| TNN<sup>e</sup> | Minimum daily $T_{\text{MIN}}$<sup>†</sup> | min | 4,8,13,17,26,52,78 |
| TNX<sup>e</sup> | Maximum daily $T_{\text{MIN}}$<sup>†</sup> | max | 4,8,13,17,26,52,78 |
| TX10p<sup>e</sup> | Cool days; days where $T_{\text{MAX}} < 10^{\text{th}}$ percentile<sup>†</sup> | % | 4,8,13,17,26,52,78 |
| TX90p<sup>e</sup> | Warm days; days where $T_{\text{MAX}} > 90^{\text{th}}$ percentile<sup>†</sup> | % | 4,8,13,17,26,52,78 |
| TXN<sup>e</sup> | Minimum daily $T_{\text{MAX}}$<sup>†</sup> | min | 4,8,13,17,26,52,78 |
| TXX<sup>e</sup> | Maximum daily $T_{\text{MAX}}$<sup>†</sup> | max | 4,8,13,17,26,52,78 |
| VP<sup>a</sup> | Vapor pressure; weekly mean atmospheric pressure exerted by water vapor | $\bar{x}$<br>$\sigma$ | 4,8,13,17,26,52,78<br>8,13,17,26,52,78 |
| VPD<sup>a</sup> | Vapor pressure deficit; weekly mean difference between saturated vapor pressure and actual vapor pressure | $\bar{x}$<br>$\sigma$ | 4,8,13,17,26,52,78<br>8,13,17,26,52,78 |
| WSD<sup>e</sup> | Warm spell duration; $\geq 6$ consecutive days where $T_{\text{MAX}} > 90^{\text{th}}$ percentile<sup>†</sup> | $\Sigma$ | 4,8,13,17,26,52,78 |

a – TerraClimate (96); b – Trenberth and Shea (98); c – Climate Prediction Center, NOAA (99); d – PTHBV v. 3.0 (93); e – Menne et al. 2012 (92), station code: SWE00140904; f – Hurrell (97); <sup>†</sup>ETCCDI index (94).

### 7.2 Variable selection

We first excluded variables with > 50% zero-valued observations during the aerosol sampling period (weeks 21-41), which removed 16 related to cold spells, ice days, frost days, and consecutive wet days. Then, we used the `findCorrelation` function in the *R* package ‘caret’ *v*. 6.0-93 to identify the largest subset with all pairwise 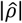 < 0.70. We curated this subset to include variables with potentially greater mechanistic importance or clearer interpretations over those that simply maximized the size of the regressor matrix (e.g., VPD over PDSI, FCF_17_ over runoff_sd_13_). The final regressor matrix comprised 75 variables with pairwise 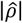 = 0.15 (*σ* = 0.13) and is provided in data S8.

### 7.3 Missing year interpolation

Daily measurements for TMIN, TMAX and air pressure were not reported from 1993-1995 by the nearest weather station.^15^ In practice, this resulted in 21 missing observations for their derived variables. We initially considered using other nearby weather stations (*92*) to supplement the observations but they either also lacked these measurements or their temporal coverage did not overlap sufficiently to assess potential inhomogeneity. Therefore, we interpolated values for 1994 for the 18 affected variables: pressure_4,8,26_, DTR_4,13,52_, FCF_17,26,78_, FD_365_, TN10p_4,13_, TN90p_4,26_, TNX_52_, TXN_52_, and TXX_26,52_. We followed the state space model framework described in section 8, with the following modifications: 1) we used the entirety of the reported data from 1959-2008 to inform parameter estimation, 2) only trigonometric seasonal dummy variables were included in the regressor matrix, and 3) we considered the model with the lowest cumulative one-step-ahead forecast errors to be the best prediction. We examined the rank-transformed time series and considered the imputed 1994 estimates to be plausible, especially for variables calculated over longer periods or with long-term trends or cycles (fig. S11).

**Fig. S11.**
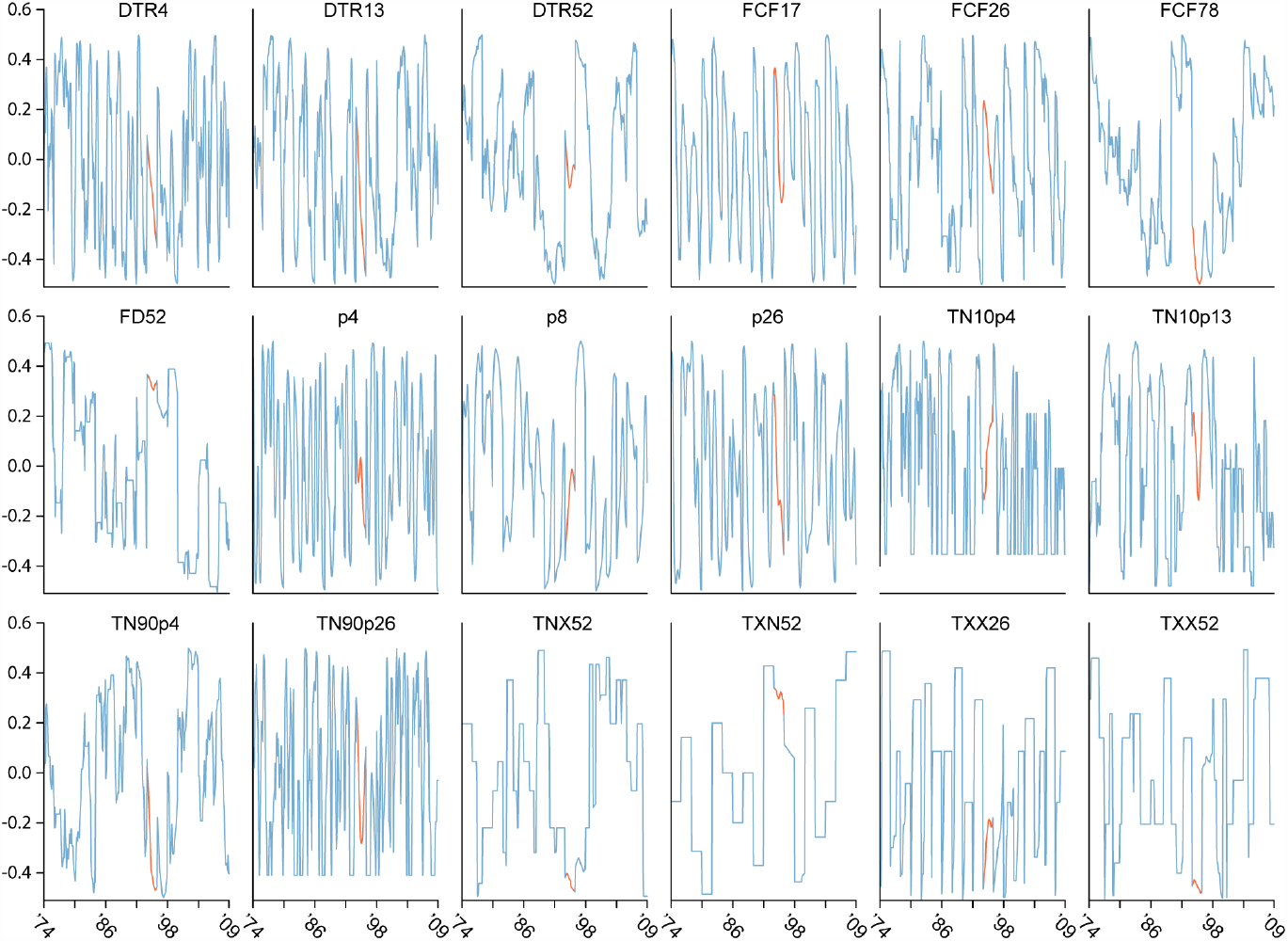
Climatic covariables with imputed values for 1994. Rank-transformed observed data is shown in blue and imputed values in orange.

### 7.4 Variable clustering and categorization

We related each of the original 393 variables to their larger ‘climatic’ neighborhood using densMAP^16^ (*101*) combined with hdbscan^17^ (*102*) with ‘densvis’ *v*. 1.8.1 (*101*) and *‘*dbscan’ 1.1-11 (*103*) for *R*, respectively. Informally, variables within a neighborhood describe the same, or a similar, climatic feature while those in distant neighborhoods are more likely generated by a different latent process.

Both densMAP and hdbscan are sensitive to hyperparameter choices. In the absence of a more objective cost function, we considered hyperparameter combinations with higher classification rates to be better summaries of the climatic data. We conducted a grid search over the densUMAP and hdbscan parameters: n_neighbors = c(10, 15, 20, 25), n_components = c(10, 20, 30, 40, 60), lambda = c(0.05, 0.1, 0.15), metric = c(“correlation”, “cosine”, “manhattan”, “euclidean”), min_samples = c(10, 11, 12, 13, 14, 15, 16, 17, 18). Cluster number varied by `min_samples`, which directly specifies the smallest permitted cluster size but no other hyperparameter had a clear individual effect, nor did any independently influence the classification rate. Combinations with classification rates above the 75^th^ percentile (*n* = 32, mean = 95.53%) most frequently resolved 3 and 7 clusters (*n* = 16 and 9, respectively). We compared the climatic variable assignments at *k* = 3 and *k* = 7 to assess their stability.

Assignments differed primarily in resolution and in identity of unclassified variables, although the Manhattan distance differed in both cases and additionally produced hierarchically incompatible *k* = 3 and *k* = 7 assignments. The remaining three *k* = 7 assignments differed by only a single successfully classified variable and were consistent with the *k* = 3 results.

We considered the *k* = 7 assignments as the best estimate of high-dimensional neighborhood space and examined each group to identify common features. Based on this, we suggest our climatic variables can be summarized as aspects of seven latent axes:

1. precipitation, which includes precipitation variables with < 52-week intervals;
2. water storage, comprising most runoff and soil moisture variables with ≥ 8-week intervals, running means of the PDSI, and precipitation variables with ≥ 52-week intervals;
3. snow accumulation, inferred from the inclusion of ≥ 52-week snow water variables and running means of the NAO and AO indices;
4. warming trend, based on the inclusion of most temperature-derived variables with ≥ 52-week intervals and all estimates of TN90p and TX90p;
5. seasonal transitions, which consists of variables delimiting the potential vegetative growth period, including the recent number of frost and ice days, temperature variability, and short window estimates of runoff, snow cover, radiation, PET, and AET;
6. evapotranspiration, a group with similar base variables as seasonal transitions but with ≥ 8-week windows, in addition to most sub-annual temperature variables and estimates of water deficit and soil moisture variability; and
7. the Atlantic Multidecadal Oscillation, which simply consists of running means of both AMO indices.

Like the climatic variables themselves, these categories are an abstraction intended to represent the environment experienced by a hypothetical organism. However, we use them as a heuristic device because they help clarify the kind of variation represented by abstruse regressors (*e*.*g*., the standard deviation of the NAOI falls on the ‘seasonal transitions’ axis), and they emphasize the relationship between trends and potentially more proximate factors, rather than a single index.

## 8. Time series analysis

### 8.1 Introduction to state-space models

We used linear state-space models (SSMs) to analyze eDNA and traditional count-based time series. Such models consider time series data to be the result of two connected stochastic systems: 1) a hidden, or latent, process that generates variation across time, and 2) a measurement process that allows discrepancies between the latent state and observed data. The relationship between sequenced eDNA from a given taxon *y* and the true DNA abundance in the catchment area *μ*, for example, can be written:

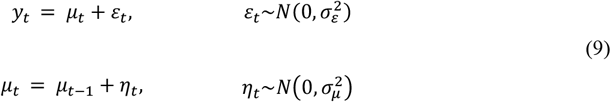

where

° *y* is the vector of eDNA abundance at time steps *t* = 1…*T*,
° ε is measurement error with variance 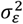,
° *μ* is the corresponding latent population size,
° and η represents variation in μ with variance 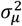.

Recursive algorithms, most commonly the Kalman filter (*104*), solve Equation 9 by formalizing the intuition that the historic performance of a model can be used to refine future predictions. The filter computes 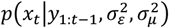 and uses the prediction errors *v*_t_ = *x*_t_ – *y*_t_ and their variance *F*_t_ to obtain minimum-variance unbiased estimates of *x*_t_ and the system parameters, in this case, 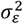 and 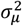 (*105*). SSMs fit by a Kalman filter can be framed in maximum-likelihood or Bayesian terms, and we employ both as a matter of accessibility given the available implementations suitable for ecological time series.

### 8.2 eDNA abundance and diversity trends

#### 8.2.1 Structural time series models

We modeled eDNA abundances observed in calendar weeks 21-41 of each year using the *R* package ‘bsts’ *v*. 0.9.9 (*106, 107*). Here, the simple model in Equation 9 is extended to include a second latent state, *δ*_t_, to allow a stochastic directional trend:

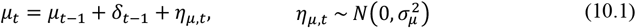

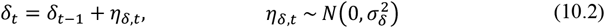

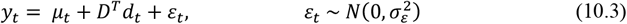

This is known as a local linear trend (LLT) or ‘random walk with drift’ model. If 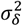 approaches zero but 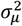 does not, the model reduces to the local level (LL) in Equation 9 and indicates that μ is equally likely to increase as decrease at each time step. Conversely, a relatively large 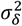 with 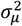 approaching zero results in an integrated random walk (IRW) model, where *μ* changes according to a stochastic but directional trend (*108*).

We tested for potential responses to climatic (Section 7) and aerosol dispersion-related (Section 1.3) variation by comparing predictive power of Equation 10 using three different sets of *d*_t_ covariables:

1. six variables representing generic seasonal patterns, defined by the trigonometric function (*105*):

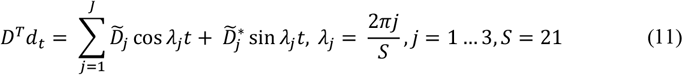

where λ denotes the *j*^th^ harmonic and *S* specifies the length of the season;
2. these combined with the 75 climatic variables described in Section 7;
3. or the trigonometric seasonality combined with the particle dispersion variables described in Section 1.3.

Our SSMs are limited to linear Gaussian cases, but ecology theory predicts unimodal or skewed responses to environmental variation (*109*). Therefore, we tested five transformations (Yeo-Johnson, exponential, minmax, ranks, and standard scores) of the covariate matrices in a regression model for each of the 17 cluster abundances. We compared their forecast errors using the diagnostic tests in Section 8.2.3 to identify which transformation best conformed with model assumptions for the majority of the clusters. Rank transformation was most consistently adequate for the climatic regressors whereas all transformations performed well with the particle dispersion variables. For better comparability between the models, we applied the rank transformation to both regressor matrices.

#### 8.2.2 Prior distribution specifications

Completing the model in Equation 10 requires specifying prior distributions on the estimated parameters 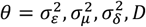. In ‘bsts,’ variance terms are drawn from the gamma distribution:

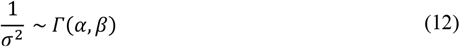

with mean *α*/*β* and variance *α*/*β*^2^. A hierarchical spike-and-slab prior is placed on the vector of regression coefficients *D*, where ζ is a Bernoulli distributed variable determining if *D* = 0 for each of the 1…*K* covariates:

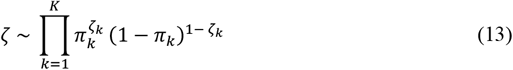

or is otherwise drawn from

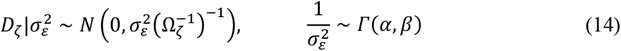

where

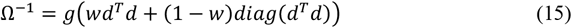

and 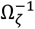 denotes the rows and columns of Ω^-1^ where ζ = 1. Equation 15 reduces to Zellner’s *g* prior when the diagonal shrinkage parameter *w* is zero. More simply, Ω^-1^ conveniently scales the prior distribution on *D*_*ζ*_ based on the covariance structure of the subset of covariates sampled in a particular draw.

We defined the priors on 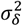 and 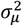 to enforce two cases of Equation 10: the local linear trend (LLT) and the integrated random walk (IRW). We compared these models explicitly because we found that in practice, LLT models simplified to an LL process when 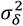 was negligible but not to an IRW process unless both 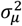 and 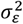 approached zero, an unlikely scenario for eDNA time series. This result is not surprising given the difficulties of estimating process error when measurement error is high. As we considered an IRW process with high measurement error and a small slope to be a plausible alternative to the LLT, we chose to enforce this outcome by fixing 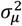 to an arbitrarily small value.

In a set of pilot runs on clusters 17 (prokaryotes), 8 (insects), and 5 (plants), posterior estimates were generally insensitive to priors on and over the unique combinations of *α* = {0.01, 0.05, 0.1, 0.5, 1, 2} and 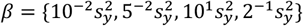, where 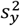 is the sample variance of the time series. However, MCMC diagnostics (Section 8.2.3) favored *α* = 1. We then selected *β* with the expectation that measurement error is the largest source of variance, followed by 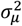 and 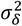, respectively. Prior distributions used in the production models are given in table S5.

**Table S5.** Prior distributions used in the production models.

| model | regressors | abbreviation | $\sigma_\mu^2$ | $\sigma_\delta^2$ | $\Omega^{-1}$ | $\pi_k$ | $\sigma_\epsilon^2$ |
| --- | --- | --- | --- | --- | --- | --- | --- |
| local<br>linear<br>trend | harmonics | LLT-base | $\alpha = 1$<br>$\beta = 10^{-1} s_y^2$<br>$\sigma_\mu^2 \leq s_y^2$ | $\alpha = 1$<br>$\beta = 10^{-2} s_y^2$<br>$\sigma_\delta^2 \leq 0.5 s_y^2$ | $g = 1$<br>$w = 0.5$ | $\frac{2}{7}$ | $\alpha = 1$<br>$\beta = 0.5 s_y^2$<br>$\sigma_\epsilon^2 \leq s_y^2$ |
| | harmonics<br>&<br>climate | LLT-climate | | | | $\frac{3}{41}$ | $\alpha = 1$<br>$\beta = 0.75 s_y^2$<br>$\sigma_\epsilon^2 \leq s_y^2$ |
| | harmonics<br>&<br>catchment | LLT-particle | | | | $\frac{2}{27}$ | $\alpha = 1$<br>$\beta = 0.75 s_y^2$<br>$\sigma_\epsilon^2 \leq s_y^2$ |
| integrated<br>random<br>walk | harmonics | IRW-base | $10^{-4}$ | $\alpha = 1$<br>$\beta = 10^{-2} s_y^2$<br>$\sigma_\delta^2 \leq 0.5 s_y^2$ | $g = 1$<br>$w = 0.5$ | $\frac{2}{7}$ | $\alpha = 1$<br>$\beta = 0.5 s_y^2$<br>$\sigma_\epsilon^2 \leq s_y^2$ |
| | harmonics<br>&<br>climate | IRW-climate | | | | $\frac{3}{41}$ | $\alpha = 1$<br>$\beta = 0.75 s_y^2$<br>$\sigma_\epsilon^2 \leq s_y^2$ |
| | harmonics<br>&<br>catchment | IRW-particle | | | | $\frac{2}{27}$ | $\alpha = 1$<br>$\beta = 0.75 s_y^2$<br>$\sigma_\epsilon^2 \leq s_y^2$ |

#### 8.2.3 Model fit and convergence diagnostics

For pilot runs exploring data transformations and prior specifications, we relied primarily on omnibus tests applied to posterior mean forecast errors to efficiently compare dozens of models. Following Commandeur and Koopman (*110*) and Durbin and Koopman (*105*), we used the *F* variance ratio between the first (*t* = 2, …, 121) and last (*t* = 254, …, 378) thirds of the time series, the magnitude and significance of autocorrelation in the first 42 lags, and Kolomogorov-Smirnov’s *d* to test for heteroscedasticity, serial dependence, and non-normality, respectively. Convergence was evaluated by calculating effective sample sizes (ESS) for each parameter, Geweke’s convergence diagnostic (*111*), Raftery and Lewis’s diagnostic (*112*) with the *R* package ‘coda’ *v*. 0.19-4 (*113*) and through visual inspection of parameter trace plots. For final model runs, we verified these summary statistics using diagnostic plots of the posterior forecast error and latent state distributions and assessed identifiability by plotting univariate prior and posterior distributions, likelihood profiles, and joint posterior distributions. Pilot models were run for 10^5^ and final models for 10^6^ MCMC iterations, with 10% discarded as burn-in.

Some models, primarily of bacteria-dominated cluster abundances, exhibited substantial evidence against normal, identical, and independent (IID) errors in all tested combinations of priors, time series models, and covariate transformations. Poor model performance was most likely caused by a small number of extreme observations in all cases. We removed data points more extreme than 1.5 × the interquartile range in log space for both abundances and diversity metrics. Forecast errors were approximately IID after removing these outliers. Diagnostic summary statistics and the transformations applied, if any, for all production runs in data S9.

#### 8.2.4 Leave-future-out cross validation

We compared the predictive accuracy of models with different trend and regression specifications using the exact expected log pointwise predictive density (ELPD) estimated by leave-future-out cross validation (LFO) (*114, 115*). Time series models that more accurately predict the next *M* future observations conditioned on data from *t* = 1 … *t*_M−1_ are more likely to be well-specified and have generalizable parameter estimates. Thus, we computed the expected log-predictive densities *p*(*y*_*t*+1:*M*_|*y*_1:*t*_) for each *t* ∈ {*L*, …, *N* − *M*}, where *L* is the minimum number of observations considered before making predictions ahead, *N* the sample size, and *M* the number of future observations:

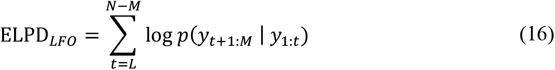

We used 21-step-ahead predictions (the number of time points in a year in our dataset) for the last 57 weeks (15%) of the time series, *i*.*e*., *M* = 21, *L* = 300 and *N* = 378. This process refits the time series model for each *t* ∈ {*L*, …, *N* − *M*} and uses *S* random draws 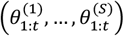 from the posterior distribution *p*(*θ*|*y*_1:*t*_) to calculate the log likelihood of *p*(*y*_*t*+1:*M*_|*y*_1:*t*_):

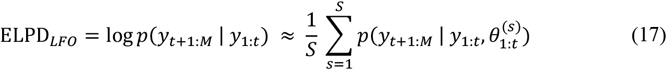

We used *S* = 3.6 × 10^5^ (*i*.*e*., 4 × 10^5^ iterations with the first 10% discarded as burn in) for cross-validation. Obtaining 57 forecasts for each model is still time consuming, but each model only needs to be fit to the full dataset once because the Kalman recursions can be re-filtered to obtain *p*(*y*_*t*+1:*M*_ | *y*_1:*t*_) at each *t* ∈ {*L*, …, *N* − *M*}. We considered a model to be the best among the candidates if the ELPD difference divided by the standard error of the difference was > 2 in all pairwise comparisons (*116*). If one or more models were similarly supported, we preferred the model with the fewest number of parameters and/or the smaller regressor matrix.

### 8.3 Abundance trends from traditional monitoring data

#### 8.3.1 Data acquisition

We conducted an extensive search of publicly-available data and consulted with government authorities to identify monitoring surveys within 100 km of the aerosol sampling station with at least seven years of data between 1973-2008. We found two programs meeting these initial requirements: the Swedish Bird Survey^18^ (*117*) and the Swedish Electrofishing Register.^19^ However, electrofishing data for the river closest to the aerosol sampling station, the Torne (< 5 km), was only available for four years after 2003. Our initial models indicated different population trajectories among and within river catchments, which suggests that the electrofishing data may not adequately represent the area closest to the aerosol sampling station. We excluded fish from further consideration, leaving birds for comparison.

The Swedish Bird Survey comprises point observations collected by volunteers according to a standardized protocol along predefined routes. We narrowed our search to routes surveyed for ≥ 5 years and with ≥ 10 total counts of a genus represented in the filter sequences. This resulted in nine genera: *Anas* (Anseriformes: Anatidae), *Corvus* (Passeriformes: Corvidae), *Cuculus* (Cuculiformes: Cuculidae), *Ficedula* (Passeriformes: Muscicapidae), *Gavia* (Gaviiformes: Gaviidae), *Lagopus* (Galiformes: Phasianidae), *Parus* (Passeriformes: Paridae), *Phylloscopus* (Passeriformes: Phylloscopidae), and *Saxicola* (Passeriformes: Muscicapidae). For the three genera with multiple species in the Kiruna region (*Corvus corax* and *carone, Anas crecca* and *penelope*, and *Lagopus lagopus* and *muta*), we summed the counts and analyzed them as a single genus.

#### 8.3.2 State space models

We modeled abundance trends from the count data using SSMs as implemented in the *R* package ‘MARSS’ *v*. 3.11.4 (*118*). For each genus, we considered survey routes as observers of the same latent population trend but with potentially different autoregressive (AR) errors (*119*). This allows each survey route to be influenced by local conditions and have different random error rates. In MARSS notation, this model is written:

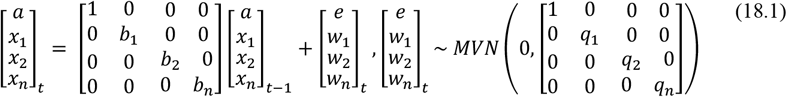

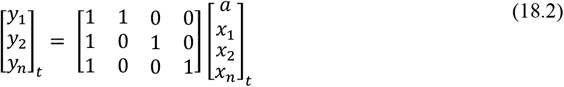

where

° *a* is the latent trend observed by all routes at time *t*;
° *b*_*n*_ is the AR(1) parameter (φ) for the 1… *l*^*th*^ route;
° *x*_*n*_ is the AR(1) trend for each route at time *t*;
° and *w*_*n*_ is the observation error for the 1… *n*^*th*^ route at time *t* with variance *q*_*n*_.

We fit maximum-likelihood models for each genus via the EM algorithm with the options: `minit = 500, maxint = 2000, abstol = 1e-6, conv.test.slope.tol = 1e-6`. Because the variance of the estimated shared trend was high for most genera (fig. S12), we calculated the two-year centered moving average before extracting even-numbered years between 2000-2008 for comparison with the eDNA estimates. Our results are similar to estimates from the Norrbotten Country Board^20^ (*120*), which suggests trends estimated from the Swedish Bird Survey’s standardized routes are robust to analysis method.

**Fig. S12.**
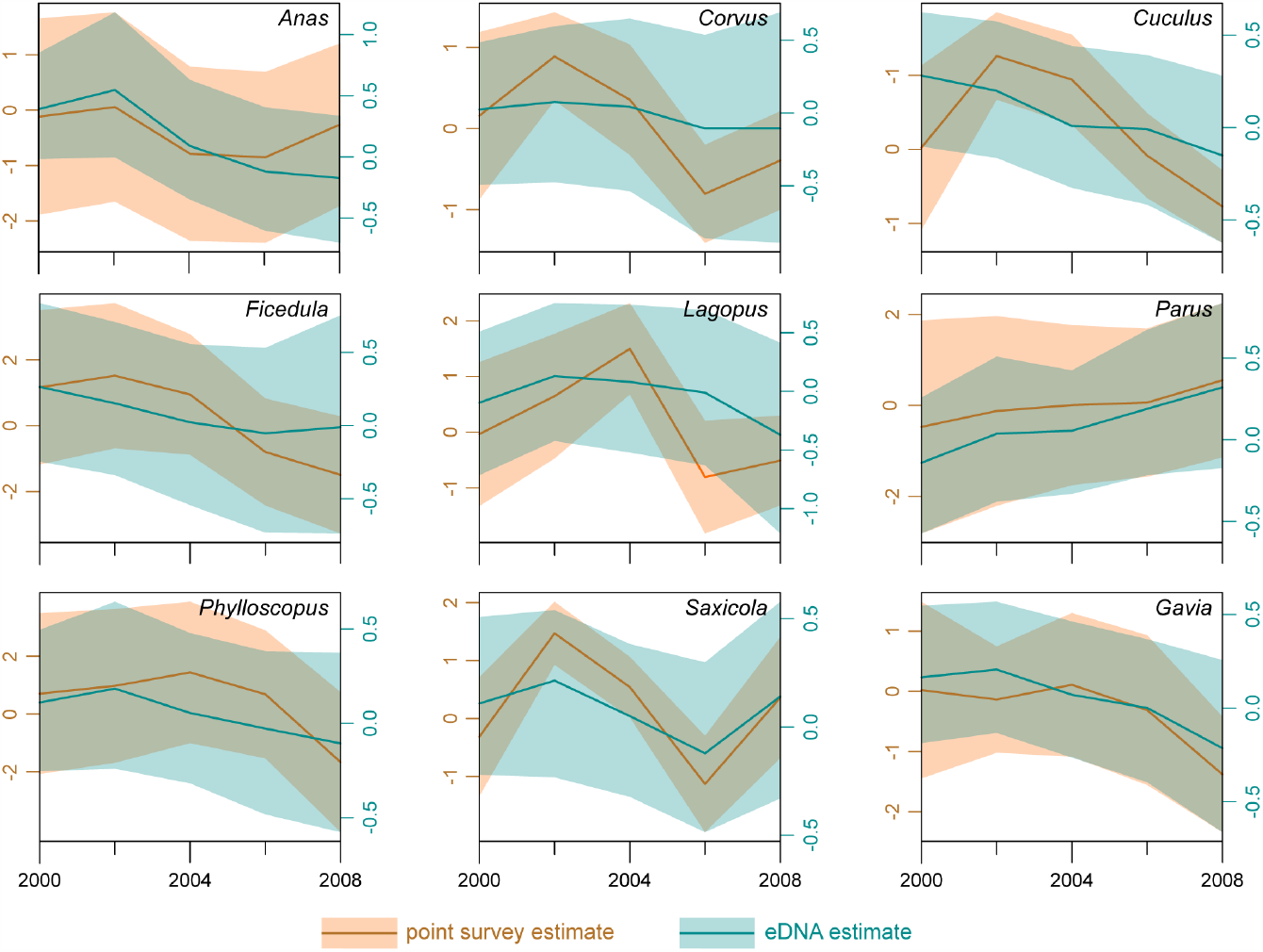
Scaled annual abundances indices for nine bird genera estimated from point surveys (orange) and PLR transformed eDNA (blue). Shaded regions show 95% confidence and credible intervals for point surveys and eDNA estimates, respectively. Note different *y*-scales are used for the two data sources.

Models of PLR-transformed eDNA abundances were estimated following the methods in Section 8. Models using the ‘climate’ regressors produced the best 1-year-ahead forecasts according to ELPD differences (Section 8.2.4) for five genera and were tied for the top rank for all genera (data S9). The LLT and IRW trend models performed similarly, but MCMC diagnostics (8.2.3) suggested the LLT models had convergence issues for some genera (data S9). We therefore used the ‘irw climate’ models for all genera and calculated annual averages from the posterior median state (fig. S12). We *z-*transformed the averaged eDNA and count estimates and estimated their correlation with ordinary least squares regression.

## 9. Land use and forest history

**Fig. S13.**
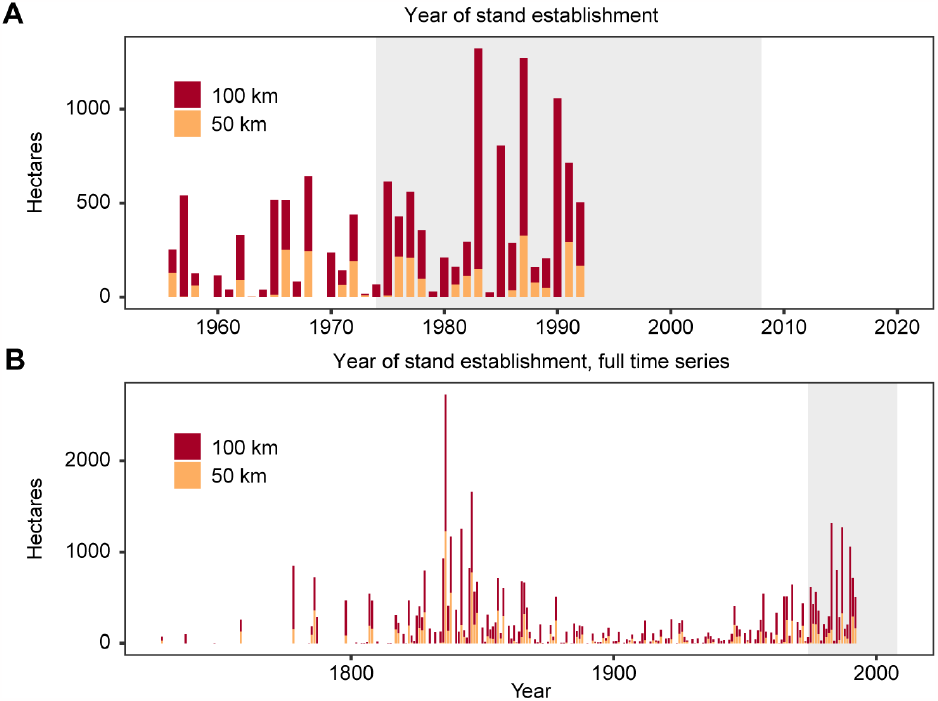
Forest harvests outside formally protected areas in northern Sweden. **A**) year of stand establishment within 50 and 100 km of the aerosol sampling station from 1955 to 1993, and **B**) with all available records, 1793-1993. Shaded areas in each panel denote years overlapping with the eDNA time series.

Forests have been a dominant, continuous presence in northern Fennoscandia since shortly after the last glacial maximum. Until the 19^th^ century, the indigenous Sami people were the majority inhabitants of the region and primarily engaged in reindeer pastoralism, hunting, fishing, and low-intensity agriculture (*121, 122*). Colonization by Swedish and Finnish-speaking agriculturalists began by the early 17^th^ century, but demographic and land use changes occurred slowly until the second half of the 19^th^ century (*121*) or as recently as the 1880s around Kiruna (*123*). We used stand age data from the Comprehensive Forest Inventory^21^ to estimate the timing and spatial extent of stand-replacing disturbances within 50 and 100 km of the aerosol sampling station (*124*). Consistent with the broader forest history in northern Fennoscandia (*122*), we found two peak periods of canopy conversions, first in the mid-1800s and later in the 1980s (fig. S13). Note that the inventory was conducted from 1982-1993 and stand establishments are likely underestimated during this period. Fire may have caused some portion of the canopy loss in the 1700 and 1800s, but clearcuts have been the dominant stand-replacing disturbance since the 1900s.

Contemporary land use comprises commercial forestry, nature-oriented tourism, reindeer husbandry by the Sami people, and large-scale mining operations. The town of Kiruna (population 23,000), inactive open-pit mines (1900-1960), and a large underground iron mine (1960-present) lie 10 km west of the aerosol station. Another iron mine located near the town of Gällivare *ca*. 90 km southwest operated open-pit until the 1960s, and the nearby Aitik open-pit copper mine was established in 1968. A smaller open-pit mine 35 km to the southeast operated from 1965-1983. A large, contiguous network of formally-protected nature reserves spans much of the subalpine zone west of the aerosol sampling station (fig. S14).

Forestry outside the subalpine zone is intensive and extensive relative to other boreal regions. For example, the Swedish National Forest Inventory^22^ (NFI) (*125*) reports *ca*. 14% of the total forested area in northern Norrland^23^ was felled between 1986-2016 (*126*), compared to 10% of the eastern boreal shield region (roughly, Ontario and further east) and 4% of the western shield (*127*). From 1982 to 2008, *ca*. 33% of northern Norrland forests received at least one silvicultural treatment (*126*).^24^ The Swedish National Land Cover Database, constructed from 2017-2019 using satellite and LiDAR data, classified 16.5% of forests within 350 km of the aerosol station (19.5% in Norbotten county) as ‘temporarily non-forested’, that is, regrowing stands with a canopy height < 5 m (*56*). Given the site indices^25^ typical of the region (*128*), these stands were likely younger than 20-40 at the time of the database construction.

**Fig. S14.**
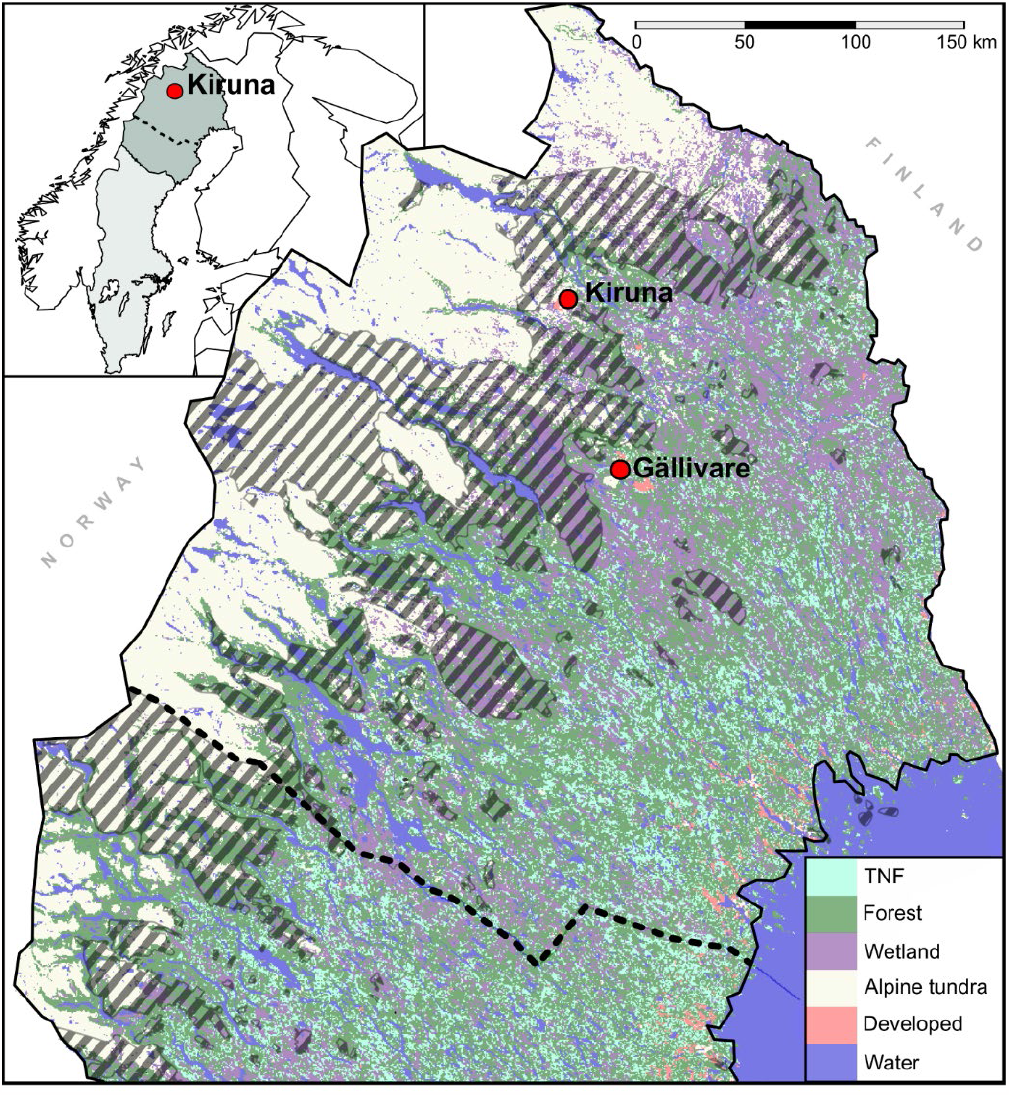
Map of land cover from the Swedish National Land Cover Database (NMD) and formally protected areas in Sweden within 350 km of the aerosol station. Land cover classes aggregated from the original 21 thematic classes: ‘TNF’ are ‘temporarily non-forested’ with regrowing trees that are < 5 m tall; ‘forests’ designates areas with > 10% crown cover and > 5 m canopy height; ‘wetlands’ denotes non-forested areas where water covers the soil most of the year; ‘alpine tundra’ refers to non-wetland areas incapable of supporting forests but may be covered by vascular plants, bryophytes, or lichens; ‘developed’ includes permanent construction, roads, railways, and a small amount of cultivated land (< 1%); and ‘water’ includes all permanent water bodies. Inset shows the position of the town of Kiruna and the two northernmost provinces of Sweden (darker and lighter shading, respectively) within Fennoscandia; the dotted line indicates the border between Norrbotten and Västerbotten.

Data from the NFI for northern Norrland (*126*) indicate the extent of > 100 year old forests declined by *ca*. 35% during the years concurrent with the eDNA time series (fig. S15A). Forests older than 160 years decreased by 55% between 1974 and 1995, or an 80% decline since 1955 (fig. S15A). The oldest forest fraction increased modestly after the mid-1990s minima (fig. S15A); these likely established from advance regeneration left by early high-grading (*129*) and some may be functionally ‘old-growth’ forests (*130*). Forest biomass (in forest cubic meters,^26^ m^3^sk) and density have generally increased since 1955, but pine has increased the most by far (fig. S15B and C).

**Fig. S15.**
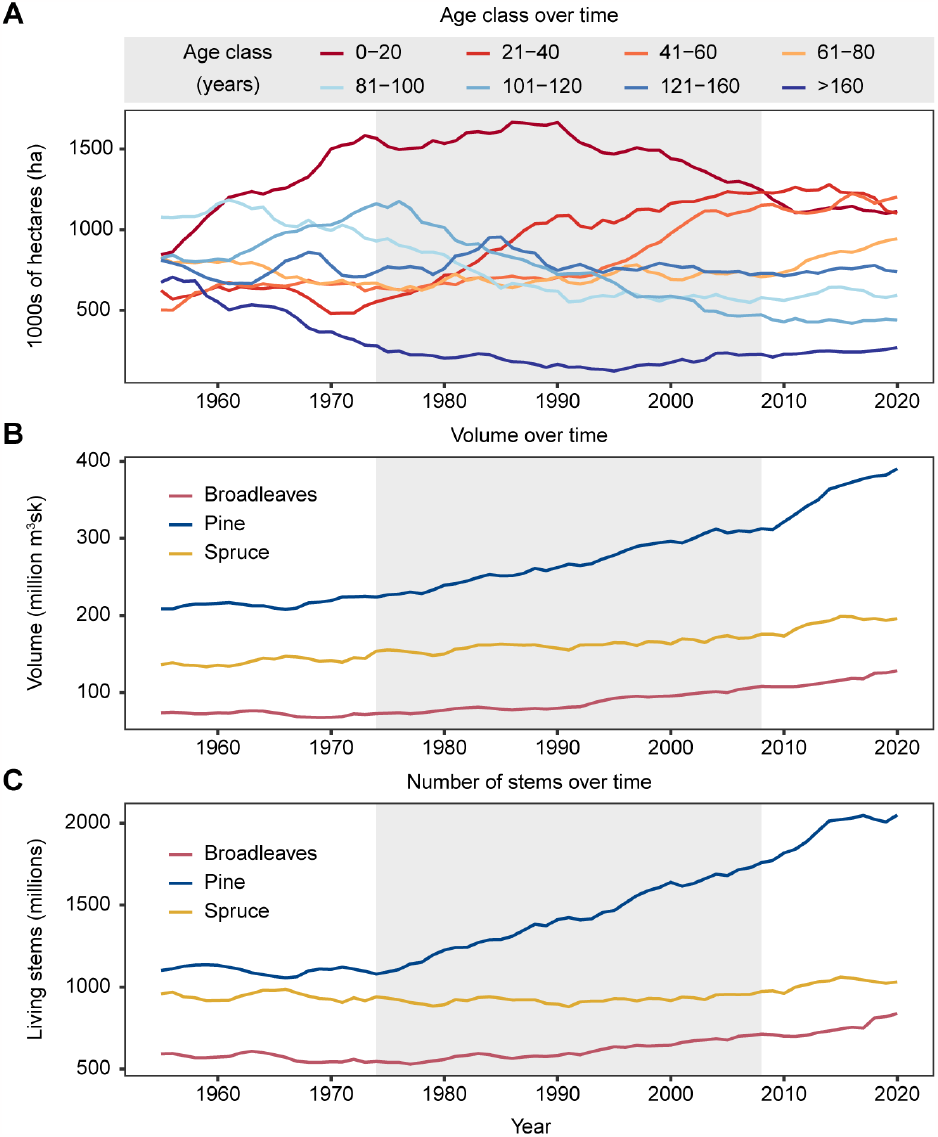
Forest age and standing biomass in northern Sweden. **A**) Productive forest area by age class, **B**) total standing volume in millions of forest cubic meters (m^3^sk) by species across all land use classes. **C**) Total number of living stems (≥ 10 cm) in all land use classes.

Higher resolution forest history data, especially integrated in a spatiotemporal framework with eDNA data, could help identify how specific silvicultural treatments or conservation interventions impact (or not) regional biodiversity. Conversely, historic reconstructions informed by archaeological datasets (*e*.*g*., (*131*)) would help verify and calibrate eDNA time series when contemporary remote sensing and monitoring data are lacking, as is the case here. For example, the peak in pine-associated eDNA we found during the mid-1990s coincides with a period of rapid change in the area covered by pine, which suggests aerosols emitted by harvest and afforestation activities may have also influenced this trend (fig. S16).

**Fig. S16.**
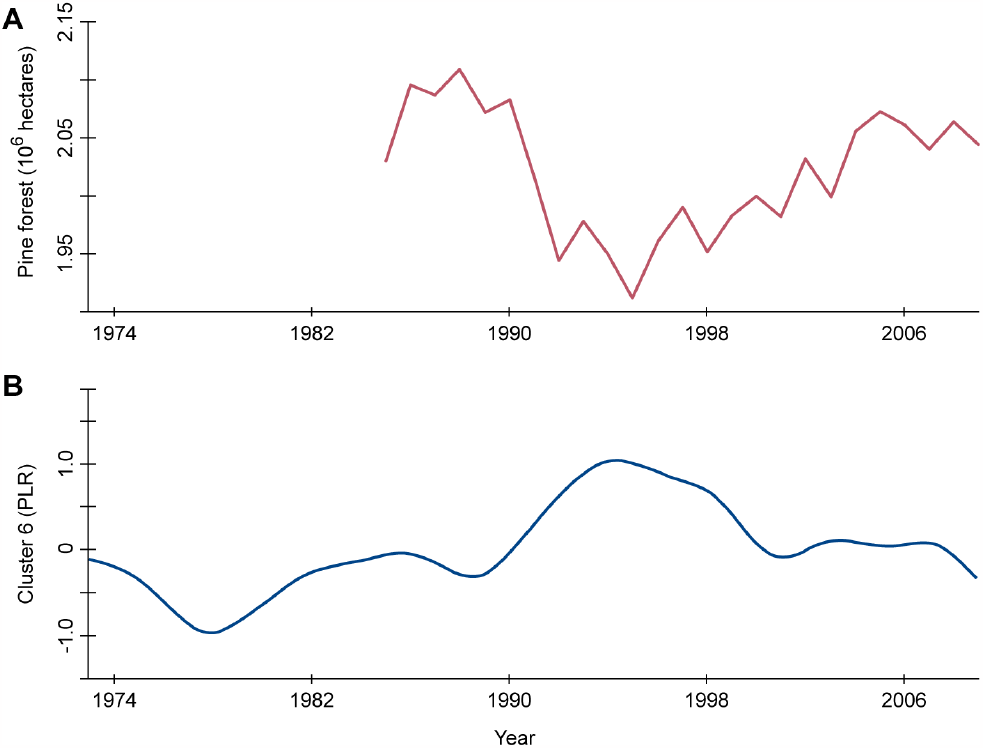
Comparison of trends in pine forest cover and pine-associated eDNA abundance. **A**) spatial extent of pine forests in Norrbotten province from 1985 to 2008 from the National Forest Inventory and **B**) weekly trend estimates of the PLR-transformed relative abundance of cluster 6, the pine-dominated cluster.

## 10. Read alignment to the *Betula* nana chloroplast genome

Weekly samples with high *Betula* relative abundance were selected to study species classification and genetic variation using the complete *Betula nana* chloroplast sequence.^27^ Air filter samples from weeks 20-26 of 1998 were selected for this analysis. Reads were merged and aligned using BBMap *v*. 38.61b with: `pairedonly = t ambiguous = ‘toss’` (other parameters set as default). The SAM output was then filtered with a custom script selecting only complete reads (150 bp) that contain up to one mismatch in the alignment. To annotate observation counts of SNPs, a naïve variant calling was performed using freebayes *v*. 1.1.0-60-gc15b070 with: `-haplotype-length = 0 -min-alternate-count = 1 -min-alternate-fraction = 0 -pooled-continuous` (unnamed parameters set as default).

## 11. Cluster 13 individual genera models

We estimated trends within Cluster 13 using the pivot coordinate transformed data for each genus using a simplified version of the state space models in Section 8: 1) we considered only the LLT model (Equation 9) for the time series component and 2) only used the trigonometric seasonal dummy variables for the regression component (Equation 11) We back-transformed the pivot coordinates to relative abundances, with the genera within the cluster summing to one each week, for display. The relative abundances for each genus in cluster 13 are displayed in fig S17.

**Fig. S17.**
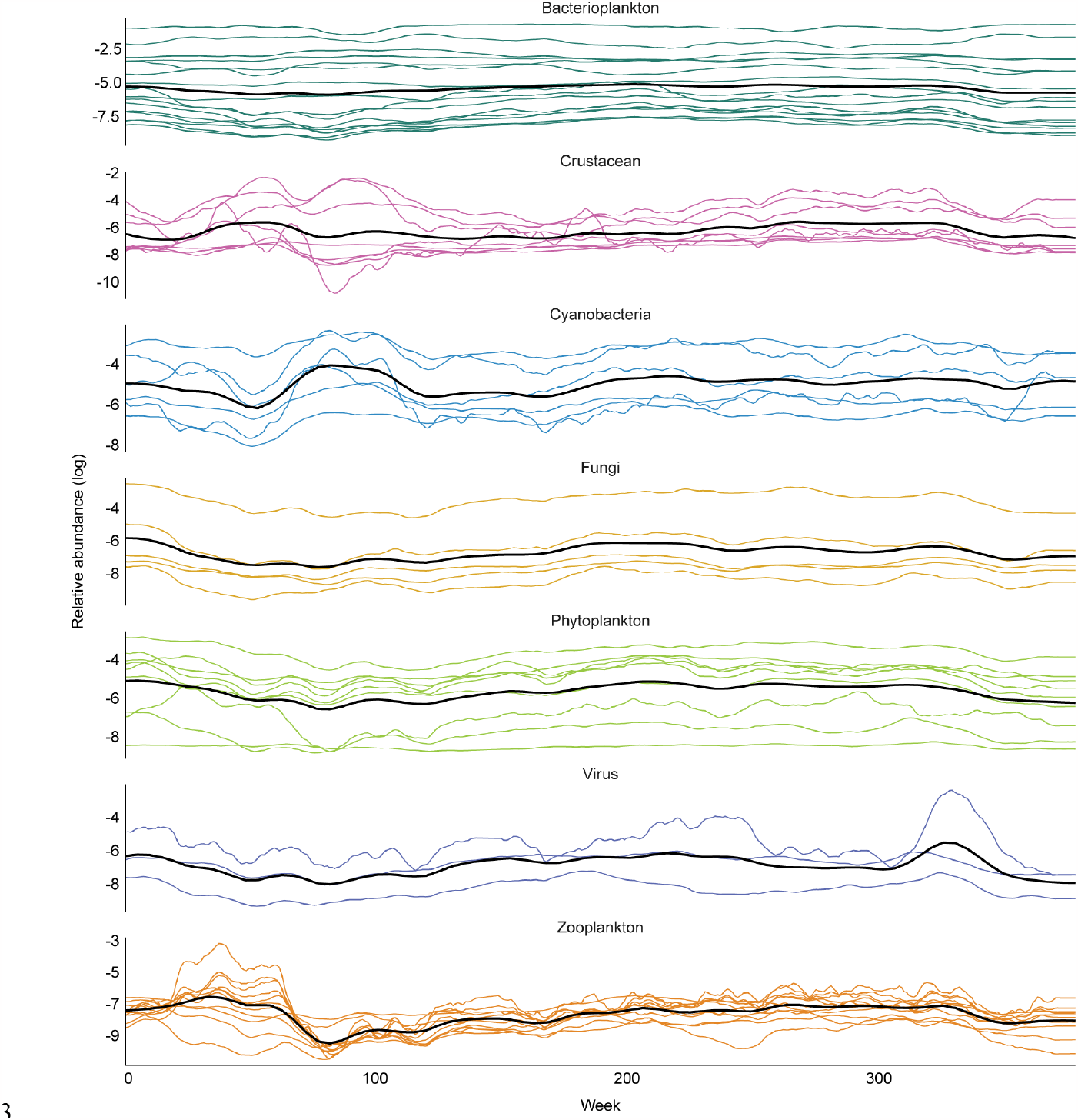
Individual relative abundance trends for genera in cluster 13. Line plots showing the relative abundances (log scale) across the time series from fitting individual models for each genus in cluster 13. Here, the genera are partitioned according to type of organism and have thus not been clustered by similarity in relative abundances. The thick black line for each group of organism indicates the mean relative abundance for that group.

## Supplementary data file descriptions

### Data S1

Particle mass originating from different distances from the aerosol sampling station for each week (sheet 1) and yearly averages as well as proportion of particle mass originating from all cardinal directions (sheet 2).

### Data S2

Taxonomic composition of the Kraken 2 database and the total sequence (in basepairs) used as input.

### Data S3

List of observed genera in Torne lappmark according to the Swedish Species Observation System.

### Data S4

Pseudolabeled genera used to train the gradient boosting classifier. “tax_id” denotes the NCBI taxonomic identification code assigned to the reads by Kraken 2 and “genus” is the corresponding name; “type” indicates if a genus was considered as a true or false positive; “set” identifies those used in model training or reserved for model testing; and columns 5-419 contain feature data and are described in the Supplementary Materials.

### Data S5

Weekly relative proportions of the 2,739 positively-classified taxa and their cluster memberships. Column “pp” denotes the predictive probability of being a true positive.

### Data S6

Summary of the taxonomic composition of the 17 clusters identified through hierarchical clustering of pairwise covariance in log-ratios. Taxonomic ranks from domain through genus that comprise ≥ 5% of a given cluster are enumerated, along with their mean relative abundance.

Taxonomy follows the NCBI taxonomic database.

### Data S7

Estimated median and 95% non-parametric confidence intervals for per-genus differences in γ-diversity contributions between 1974-1988 and 1994-2008. Negative values indicate a larger contribution in 1994-2008. *P-*values were adjusted with the Benjamini-Hochberg procedure (5% FDR). Cluster membership and NCBI taxonomy are provided for convenience.

### Data S8

Climatic regressor matrix used in time series models. Variable abbreviations correspond to table s4.

### Data S9

Summary of Bayesian state space model fit and convergence diagnostics for production runs. Models are grouped by sheet, where ‘abundances’ refers to cluster abundance models using the full time series data, ‘catchment’ refers to abundances truncated to match the time period of the particle dispersion models, ‘diversity’ contains α-, β-, γ-diversity of order *q* = 1, 2, and 3 for each of the the ‘total’, ‘no14’ and ‘eukaryotic’ fractions of the eDNA community; and ‘birds’ contains the summary results for nine genera with contemporaneous survey data. The trend and regressor matrix specification comprising the model are indicated, and the expected log pointwise predictive densities (ELPD), its standard error (ELPD.SE), along with model prediction and residual standard deviations and r^2^. Residual diagnostics include the maximum residual autocorrelation (acf.max) and its lag (acf.max.lag), the *F* variance ratio, and Kolomogorov-Smirnov’s *d* (KS.d). Effective sample sizes (ESS), the Geweke statistic, and Raftery and Lewis’s diagnostic (RL) are given for each parameter. The ELPD difference (ELPD.diff) and the standard error of this difference (ELPD.diff.se) is reported between a given model and the highest-scoring model in a comparison.

### Data S10

Supplemental time series model results for eDNA temporal cluster abundances and community diversity metrics.

### Data S11

Marginal inclusion probabilities and median coefficient estimates with 95% credible intervals for each regressor. Results are shown for climatic regression models that were supported over alternative specifications by differences in expected log pointwise predictive densities (ELPD).

## Footnotes

1 https://github.com/danisven/kraken2

2 https://github.com/danisven/StringMeUp

3 https://github.com/DerrickWood/kraken2/wiki/Manual#confidence-scoring

4 date: 2 January 2020

5 date: 11 December 2019

6 ftp://ftp.ncbi.nlm.nih.gov/blast/db/

7 https://www.ncbi.nlm.nih.gov/Traces/wgs/

8 https://trace.ncbi.nlm.nih.gov/Traces/sra/sra.cgi?view=software

9 a historic administrative division, roughly extending 100 km north, east, and west and 15 km south of the aerosol sampling station

10 Artportalen, a repository for biological surveys in Sweden and quality-reviewed community observations: https://artportalen.se/

11 Artportalen, a repository for biological surveys in Sweden and quality-reviewed community observations: https://artportalen.se/

12 NCBI Bioproject accession number PRJNA767205

13 GenBank accession number: GCA_004027795.1

14 World Meteorological Organization (WMO) number: SWE00140904

15 World Meteorological Organization (WMO) number: SWE00140904

16 density-preserving manifold approximation and projection

17 hierarchical density-based spatial clustering of applications with noise

18 Svensk Fågeltaxering; http://www.fageltaxering.lu.se

19 https://www.slu.se/institutioner/akvatiska-resurser/databaser/elfiskeregistret

20 Länsstyrelsen Norrbotten

21 Översiktlig skogsinventering; a national inventory of privately-owned property > 20 hectares conducted between 1982 and 1993

22 Riksskogstaxeringen (*125*)

23 a historic region used for statistical reporting comprising the two northernmost provinces of Sweden

24 calculated as the sum of hectares ‘cleaned’ (röjning in Swedish), thinned or felled and the total forest area (skogsmark) including alpine regions in 2020 for northern Norrland; data available from the Swedish National Forest Inventory (*126*)

25 ståndortsindex

26 the solid over-bark volume from stump to the top of the bole; skogskubikmeter

27 GenBank accession number: MT872530.1

