## Supplementary material for "Airborne eDNA captures three decades of ecosystem biodiversity": Data_S6

**Cluster 1 ($n$ = 202; $\phi_s$ = 0.40, $\sigma^2$ = 0.22)**

- KINGDOM: Fungi, 95.55%
- PHYLUM: Basidiomycota, 94.28%
    - CLASS: Agaricomycetes, 86.00%
    - ORDER: Agaricales, 10.14%
        - FAMILY: Tricholomataceae, 5.96%
    - ORDER: Atheliales, 37.78%
        - FAMILY: Atheliaceae, 37.78%
        - GENUS: *Fibularhizoctonia*, 37.72%
    - ORDER: Boletales, 28.23%
        - FAMILY: Suillaceae, 22.15%
        - GENUS: *Suillus*, 22.15%
    - ORDER: Russulales, 5.07%
        - FAMILY: Russulaceae, 5.03%
    - CLASS: Pucciniomycetes, 8.27%
    - ORDER: Pucciniales, 8.27%
        - FAMILY: Melampsoraceae, 7.17%
        - GENUS: *Melampsora*, 7.17%

**Cluster 2 ($n$ = 390; $\phi_s$ = 0.54, $\sigma^2$ = 0.19)**

- KINGDOM: Fungi, 94.11%
    - PHYLUM: Ascomycota, 67.44%
        - CLASS: Dothideomycetes, 11.54%
            - ORDER: Capnodiales, 5.18%
            - ORDER: Pleosporales, 5.19%
        - CLASS: Leotiomycetes, 39.59%
            - ORDER: Helotiales, 38.01%
                - FAMILY: Lachnaceae, 17.26%
                    - GENUS: *Lachnellula*, 17.11%
                - FAMILY: Sclerotiniaceae, 9.12%
                    - GENUS: *Botrytis*, 6.85%
        - CLASS: Sordariomycetes, 7.34%
    - PHYLUM: Basidiomycota, 26.66%
        - CLASS: Agaricomycetes, 25.78%
            - ORDER: Polyporales, 16.79%
                - FAMILY: Fomitopsidaceae, 6.28%
                    - GENUS: *Fomitopsis*, 6.26%
                - FAMILY: Polyporaceae, 9.79%
                - GENUS: *Trametes*, 9.65%

**Cluster 3 ($n$ = 37; $\phi_s$ = 0.38, $\sigma^2$ = 0.19)**

- KINGDOM: Viridiplantae, 100.00%
  - PHYLUM: Streptophyta, 100.00%
    - CLASS: Bryopsida, 53.30%
      - ORDER: Bryales, 6.23%
        - FAMILY: Mniaceae, 5.61%
      - ORDER: Hypnales, 41.23%
        - FAMILY: Hylocomiaceae, 38.74%
          - GENUS: *Pleurozium*, 38.74%
    - CLASS: Polytrichopsida, 46.58%
      - ORDER: Polytrichales, 46.58%
        - FAMILY: Polytrichaceae, 46.58%
          - GENUS: *Pogonatum*, 26.89%
          - GENUS: *Polytrichum*, 18.96%

**Cluster 4 ($n$ = 109; $\phi_s$ = 0.62, $\sigma^2$ = 0.23)**

- KINGDOM: Fungi, 21.61%
  - PHYLUM: Basidiomycota, 14.15%
    - CLASS: Pucciniomycetes, 12.36%
      - ORDER: Pucciniales, 12.36%
        - FAMILY: Cronartiaceae, 12.36%
          - GENUS: *Cornartium*, 12.19%
- KINGDOM: Viridiplantae, 76.81%
  - PHYLUM: Streptophyta, 76.81%
    - CLASS: Magnoliopsida, 74.71%
      - ORDER: Poales, 73.92%
        - FAMILY: Poaceae, 73.92%
          - GENUS: *Dactylis*, 36.42%
          - GENUS: *Triticum*, 15.67%
          - GENUS: *Hordeum*, 14.52%

**Cluster 5 ($n$ = 79; $\phi_s$ = 0.62, $\sigma^2$ = 0.23)**

- KINGDOM: Viridiplantae, 97.67%
    - PHYLUM: Streptophyta, 97.67%
        - CLASS: Magnoliopsida, 32.98%
            - ORDER: Fagales, 27.83%
                - FAMILY: Betulaceae, 27.43%
                    - GENUS: *Betula*, 27.04%
        - CLASS: Pinopsida, 61.18%
            - ORDER: Pinales, 61.18%
                - FAMILY: Pinaceae, 61.18%
                    - GENUS: *Picea*, 61.18%

**Cluster 6 ($n$ = 72; $\phi_s$ = 0.67, $\sigma^2$ = 0.24)**

KINGDOM: Viridiplantae, 97.27%

PHYLUM: Streptophyta, 97.27%

CLASS: Pinopsida, 97.04%

ORDER: Pinales, 97.04%

FAMILY: Pinaceae, 97.04%

GENUS: *Pinus*, 75.68%

GENUS: *Larix*, 21.36%

**<u>Cluster 7 ($n$ = 100; $\phi_s$ = 0.67, $\sigma^2$ = 0.18)</u>**

- KINGDOM: Unclassified, 28.33%
    - Phlyum: Discosea, 7.66%
        - CLASS: Unclassified, 7.66%
            - ORDER: Longamoebia, 7.66%
                - FAMILY: Acanthamoebidae, 7.66%
                    - GENUS: *Acanthamoeba*, 7.66%
    - PHYLUM: Evosea, 5.49%
        - CLASS: Eumycetozoa, 5.49%
            - ORDER: Unclassified, 5.49%
                - FAMILY: Unclassified, 5.49%
                    - GENUS: *Synstelium*, 5.59%
    - PHYLUM: Unclassified, 7.39%
- KINGDOM: Fungi, 52.18%
    - PHYLUM: Ascomycota, 38.58%
        - CLASS: Saccharomycetes, 36.66%
            - ORDER: Saccharomycetales, 36.66%
                - FAMILY: Debaryomycetaceae, 8.95%
                - FAMILY: Metschnikowiaceae, 10.20%
                    - GENUS: *Metschnikowia*, 8.47%
                - FAMILY: Saccharomycetaceae, 5.17%
    - PHYLUM: Mucoromycota, 9.62%
        - CLASS: Mucoromycetes, 7.29%
            - ORDER: Mucorales, 7.29%
- KINGDOM: Metazoa, 8.10%
    - PHYLUM: Tardigrada, 5.11%
        - CLASS: Eutardigrada, 5.11%
            - ORDER: Parachela, 5.11%
                - FAMILY: Hypsibiidae, 5.11%
                    - GENUS: *Hypsibius*, 5.11%
- KINGDOM: Viridiplantae, 11.11%
    - PHYLUM: Chlorophyta, 10.05%
        - CLASS: Chlorophyceae, 5.38%

**Cluster 8 ($n$ = 132; $\phi_s$ = 0.62, $\sigma^2$ = 0.17)**

- KINGDOM: Bacteria, 8.27%
  - PHYLUM: Proteobacteria, 7.77%
    - CLASS: Alphaproteobacteria, 7.59%
      - ORDER: Rickettsiales, 7.59%
- KINGDOM: Metazoa, 91.67%
  - PHYLUM: Arthropoda, 91.39%
    - CLASS: Insecta, 91.36%
      - ORDER: Diptera, 89.48%
        - FAMILY: Cecidomyiidae, 36.55%
          - GENUS: *Contarinia*, 23.77%
          - GENUS: *Mayetiola*, 11.38%
        - FAMILY: Chironomidae, 30.67%
          - GENUS: *Belgica*, 17.65%
          - GENUS: *Chironomus*, 10.75%
        - FAMILY: Trichoceridae, 8.88%
        - GENUS: *Trichocera*, 8.88%

**Cluster 9 ($n$ = 343; $\phi_s$ = 0.37, $\sigma^2$ = 0.18)**

- KINGDOM: Metazoa, 96.32%
  - PHYLUM: Arthropoda, 91.63%
    - CLASS: Insecta, 90.26%
      - ORDER: Lepidoptera, 70.18%
        - FAMILY: Geometridae, 61.33%
          - GENUS: *Operophtera*, 60.03%

**Cluster 10 ($n$ = 46; $\phi_s$ = 0.60, $\sigma^2$ = 0.16)**

- KINGDOM: Bacteria, 88.66%
  - PHYLUM: Proteobacteria, 86.55%
    - CLASS: Gammaproteobacteria, 79.99%
      - ORDER: Enterobacterales, 73.83%
        - FAMILY: Enterobacteriaceae, 10.65%
          - GENUS: Candidatus Hamiltonella, 10.65%
        - FAMILY: Erwiniaceae, 18.42%
          - GENUS: *Buchnera*, 18.42%
        - FAMILY: Morganellaceae, 14.75%
          - GENUS: *Arsenophonus*, 7.23%
          - GENUS: *Providencia*, 5.90%
        - FAMILY: Pectobacteriaceae, 30.01%
          - GENUS: *Sodalis*, 30.01%
- KINGDOM: Metazoa, 11.30%
  - PHYLUM: Arthropoda, 11.30%
    - CLASS: Insecta, 10.34%
      - ORDER: Diptera, 10.34%
        - FAMILY: Cecidomyiidae, 10.34%
          - GENUS: *Arthrocnodax*, 10.34%

**Cluster 11 ($n$ = 32; $\phi_s$ = 0.39, $\sigma^2$ = 0.14)**

- KINGDOM: Bacteria, 100.00%
  - PHYLUM: Proteobacteria, 100%
    - CLASS: Gammaproteobacteria, 100.00%
      - ORDER: Enterobacterales, 98.63%
        - FAMILY: Erwiniaceae, 36.58%
          - GENUS: *Erwinia*, 32.30%
        - FAMILY: Yersiniaceae, 51.34%
          - GENUS: *Rahnella*, 30.51%
          - GENUS: *Serratia*, 14.49%

**Cluster 12 ($n$ = 70; $\phi_s$ = 0.67, $\sigma^2$ = 0.20)**

- KINGDOM: Fungi, 89.91%
  - PHYLUM: Ascomycota: 89.73%
    - CLASS: Lecanoromycetes, 89.63%
      - ORDER: Lecanorales, 83.27%
        - FAMILY: Parmeliaceae, 83.25%
          - GENUS: *Pseudevernia*, 53.04%
          - GENUS: *Evernia*, 27.15%
      - ORDER: Umbilicariales, 6.33%
        - FAMILY: Umbilicariaceae, 6.32%
          - GENUS: *Umbilicaria*, 6.32%
- KINGDOM: Viridiplantae, 7.54%

**Cluster 13 ($n$ = 58; $\phi_s$ 0.79, $\sigma^2$ = 0.27)**

- KINGDOM: Bacteria, 85.66%
  - PHYLUM: Actinobacteria, 30.43%
    - CLASS: Actinobacteria, 30.43%
      - ORDER: Candidatus Nanopelagicales, 30.43%
        - FAMILY: Candidatus Nanopelagicaceae, 30.43%
          - GENUS: Candidatus Planktophila, 23.94%
          - GENUS: Candidatus Nanopelagicus, 6.50%
  - Phlyum: Cyanobacteria, 6.88%
    - CLASS: Unclassified, 6.55%
      - ORDER: Synechococcales, 6.88%
  - PHYLUM: Proteobacteria, 48.08%
    - CLASS: Betaproteobacteria, 47.89%
      - ORDER: Burkholderiales, 47.89%
        - FAMILY: Comamonadaceae, 41.07%
          - GENUS, *Limnohabitans*, 38.36%
- KINGDOM: Metazoa, 8.69%
  - PHYLUM: Arthropoda, 8.65%
    - CLASS: Hexanauplia, 8.65%
      - ORDER: Cyclopoida, 8.65%
        - FAMILY: Cyclopoida, 8.24%

**Cluster 14 (*n* = 208; $\phi_s$ 0.62, $\sigma^2$ = 0.18)**

- KINGDOM: Bacteria, 68.84%
    - PHYLUM: Actinobacteria: 39.74%
        - CLASS: Actinobacteria: 39.51%
            - ORDER: Actinomycetales, 6.78%
                - FAMILY: Actinomycetaceae, 6.78%
                    - GENUS: *Actinomyces,* 6.50%
            - ORDER: Corynebacteriales, 13.89%
                - FAMILY: Corynebacteriaceae, 12.36%
                    - GENUS: *Corynebacterium*, 12.36%
            - ORDER: Micrococcales, 9.09%
                - FAMILY: Micrococcaceae, 7.62%
            - ORDER: Propionibacteriales, 9.69%
                - FAMILY: Propionibacteriaceae, 9.69%
                    - GENUS: *Cutibacterium*, 9.14%
    - PHYLUM: Firmicutes, 16.94%
        - CLASS: Bacilli, 13.83%
            - ORDER: Lactobacillales, 9.91%
                - FAMILY: Streptococcaceae, 7.51%
                    - GENUS: *Streptococcus*, 7.26%
    - PHYLUM: Proteobacteria, 8.49%
        - CLASS: Gammaproteobacteria, 6.04%
- KINGDOM: Fungi, 7.20%
    - PHYLUM: Basidomycota, 7.08%
        - CLASS: Malasseziomycetes, 7.08%
            - ORDER: Malasseziales, 7.08%
                - FAMILY: Malasseziaceae, 7.08%
                    - GENUS: *Malassezia*, 7.08%
- KINGDOM: Metazoa, 23.12%
    - PHYLUM: Chordata, 23.11%
        - CLASS: Mammalia, 23.11%
            - ORDER: Artiodactyla, 9.77%
                - FAMILY: Suidae, 9.77%
                    - GENUS: *Sus*, 9.77%
            - ORDER: Carnivora, 8.93%
                - FAMILY: Canidae, 8.93%
                    - GENUS: *Canis*, 8.93%

### Cluster 15 (*n* = 32; $\phi_s$ 0.34, $\sigma^2$ = 0.13)

- KINGDOM: Bacteria, 99.91%
  - PHYLUM: Actinobacteria, 42.17%
    - CLASS: Actinobacteria, 41.15%
      - ORDER: Corynebacteriales, 31.96%
        - FAMILY: Dietziaceae, 31.96%
          - GENUS: *Dietzia*, 31.96%
      - ORDER: Micrococcales, 9.19%
        - FAMILY: Microbacteriaceae, 8.91%
          - GENUS: *Microcella*, 8.75%
  - PHYLUM: Proteobacteria
    - CLASS: Alphaproteobacteria, 7.38%
      - ORDER: Rhizobiales, 5.01%
    - CLASS: Gammaproteobacteria
      - ORDER: Oceanospirillales, 41.31%
        - FAMILY: Halomonadaceae, 41.31%
          - GENUS: *Halmonas*, 41.31%

**<u>Cluster 16 ($n$ = 180; $\phi_s$ = 0.75, $\sigma^2$ = 0.16)</u>**

- KINGDOM: Bacteria, 98.29%
  - PHYLUM: Bacteroidetes, 36.70%
    - CLASS: Flavobacteriia, 26.63%
      - ORDER: Flavobacteriales, 26.63%
        - FAMILY: Flavobacteriaceae, 26.62%
          - GENUS: *Chryseobacterium*, 18.66%
          - GENUS: *Flavobacterium*, 7.20%
    - CLASS: Sphingobacteriia, 9.93%
      - ORDER: Sphingobacteriales, 9.93%
        - FAMILY: Sphingobacteriaceae, 9.93%
          - GENUS: *Pedobacter*, 9.92%
  - PHYLUM: Firmicutes, 28.95%
    - CLASS: Bacilli, 25.45%
      - ORDER: Bacillales, 16.90%
        - FAMILY: Paenibacillaceae, 11.11%
          - GENUS: *Paenibacillus*, 10.49%
      - ORDER: Lactobacillales, 8.55%
        - FAMILY: Carnobacteriaceae, 5.45%
          - GENUS: *Carnobacterium*, 5.43%
  - PHYLUM: Proteobacteria, 30.30%
    - CLASS: Betaproteobacteria, 28.69%
      - ORDER: Burkholderiales, 28.50%
        - FAMILY: Burkholderiaceae, 25.74%
          - GENUS: *Burkholderia*, 8.92%
          - GENUS: *Paraburkholderia*, 8.17%
          - GENUS: *Ralstonia*, 5.51%

**Cluster 17 ($n$ = 649; $\phi_s$ = 0.47, $\sigma^2$ = 0.16)**

- KINGDOM: Bacteria, 99.83%
    - PHYLUM: Actinobacteria, 18.23%
        - CLASS: Actinobacteria, 18.23%
            - ORDER: Micrococcales, 6.86%
    - PHYLUM: Bacteroidetes, 6.23%
        - CLASS: Cytophagia, 6.04%
            - ORDER: Cytophagales, 6.04%
                - FAMILY: Hymenobacteraceae, 5.76%
                    - GENUS: *Hymenobacter*, 5.72%
    - PHYLUM: Proteobacteria, 72.68%
        - CLASS: Alphaproteobacteria, 26.84%
            - ORDER: Rhizobiales, 8.73%
                - FAMILY: Methylobacteriaceae, 5.01%
            - ORDER: Sphingomonadales, 14.28%
                - FAMILY: Sphingomonadaceae, 13.94%
                    - GENUS: *Sphingomonas*, 12.82%
        - CLASS: Betaproteobacteria, 22.88%
            - ORDER: Burkholderiales, 22.64%
                - FAMILY: Oxalobacteraceae, 16.75%
                    - GENUS: *Massilia*, 14.90%
        - CLASS: Gammaproteobacteria, 22.78%
            - ORDER: Pseudomonadales, 22.04%
                - FAMILY: Pseudomonadaceae, 22.04%
                    - GENUS: *Pseudomonas*, 22.03%
