## Supplementary material for "Airborne eDNA captures three decades of ecosystem biodiversity": Data_S10

Science
AAAS

### Supplementary Results for

#### Airborne eDNA captures three decades of ecosystem biodiversity

Alexis R. Sullivan§, Edvin Karlsson§, Daniel Svensson, Björn Brindefalk, Jose Antonio Villegas, Amanda Mikko, Daniel Bellieny, Abu Bakar Siddique, Anna-Mia Johansson, Håkan Grahn, David Sundell, Anita Norman, Per-Anders Esseen, Andreas Sjödin, Navinder J Singh, Tomas Brodin, Mats Forsman and Per Stenberg

§: These authors contributed equally.

**The PDF file includes:**

Supplementary Results: Bayesian state-space time series models of cluster abundances

Figs. SR1 to SR31

#### Bayesian state-space time series models of cluster abundances

We used Bayesian state-space models to decompose the abundances of the temporal eDNA clusters into components predicted by latent trends, abiotic conditions, and observation error. As trend models, we compared time-dynamic versions of the local linear trend (LLT), which reduces to a random walk (or local level, LL) if the latent slope approaches zero, and the integrated random walk (IRW), which effectively models the second-differences of the data (Supplementary Materials Section 8). We paired these two trend models with three different regressor matrices comprising: 1) six trigonometric variables with a fixed wavelength, phase, and amplitude to represent seasonal periodicity ('base'; Supplementary Materials Equation 11), 2) these plus 25 variables summarizing weekly variation in the aerosol station catchment area ('particle'; Supplementary Materials Section 1.3), and 3) the six trigonometric variables plus 75 climatic variables that represent weekly to multiannual change in temperature, precipitation, and water-balance parameters ('climatic'; Supplementary Materials Section 7). We used leave-future-out cross-validation to estimate the expected log pointwise predictive density (ELPD) of trend-regressor combinations, with the expectation that better-specified and more general models should make more accurate predictions (Supplementary Materials Section 8.2.4). If models could not be distinguished by ΔELPD, we considered the model with fewer parameters and/or a smaller regressor matrix to be 'best' of the candidates. Catchment-related variables could only be calculated from 1980 (Supplementary Materials Section 1.3), so we first constructed models using data from 1980-2008 to compare all three regressor matrices, and then from the full time series to compare the 'climatic' and 'base' matrices.

*Catchment area regression*

Regression covariables related to weekly changes in the aerosol station catchment area did not clearly improve predictions for any cluster (data S9). For three clusters (8, 10, and 17), models using the particle matrix tied for 'best' designation ($\Delta ELPD/se(\Delta ELPD) < 2$) (data S9). However, none of the particle variables in these models had high inclusion probabilities (maximum marginal inclusion $p(\zeta_k = 1) = 0.20$; cluster17), and none were included with $\geq 0.10$ frequency in the cluster10 model.

*Trend components*

Ten of the seventeen temporal clusters were best predicted by an IRW trend model (fig. SR1; data S9). Abundances with IRW-like dynamics are best (or most parsimoniously) predicted by a slowly-changing trend, potentially because zero-mean stochastic changes are relatively small or because the observations do not contain enough information to distinguish them from observation error. The seven clusters best predicted by an LLT model ranged from cases with a zero slope over the majority of the time series, indicating mostly random-walk (LL) dynamics, to those with sustained non-zero slopes (fig. SR2). Of this latter group, clusters 8 (Diptera: Cecidomyiidae, Chironomidae, and Trichoceridae) and 15 (bacteria similar to Gammaproteobacteria: *Halmonas* and Actinobacteria: *Dietzia*; data S6) appear most likely to be shaped by both a level and slope component (fig. SR2).

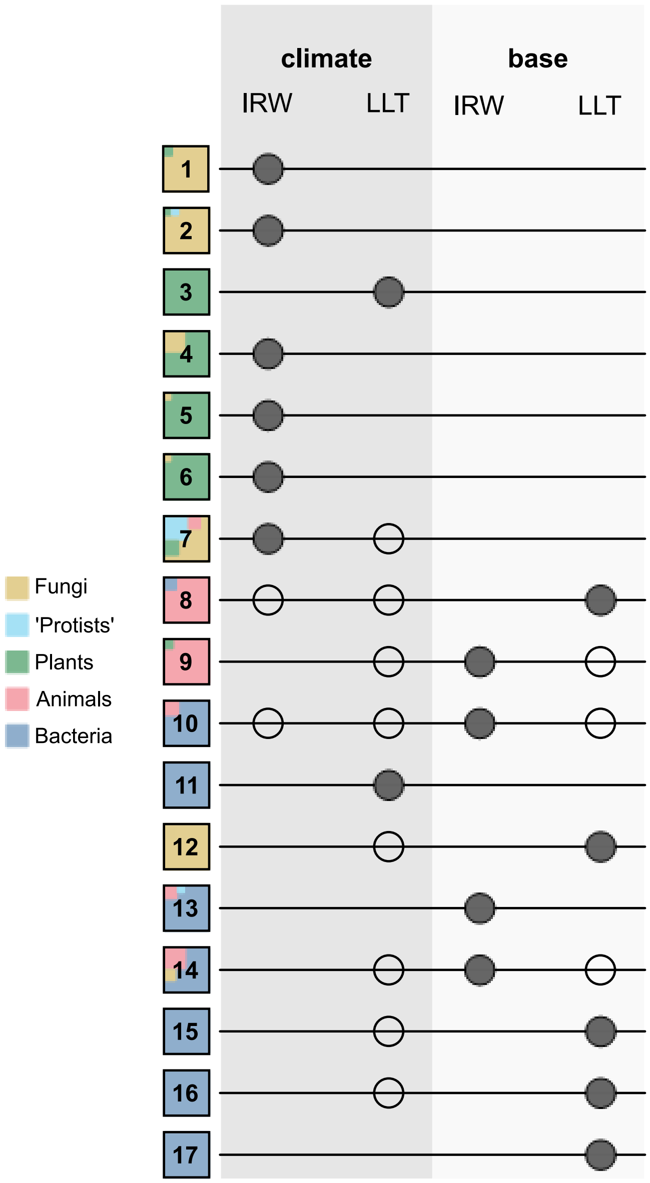


**Fig. SR1. Summary of the trend and regression components in the best-fitting models for each temporal eDNA cluster.** Clusters are numbered 1 through 17 and the nested boxes indicate their kingdom-rank relative abundance composition. Shaded circles denote the best or most parsimonious model and open circles are specifications with equal ΔELPD support.

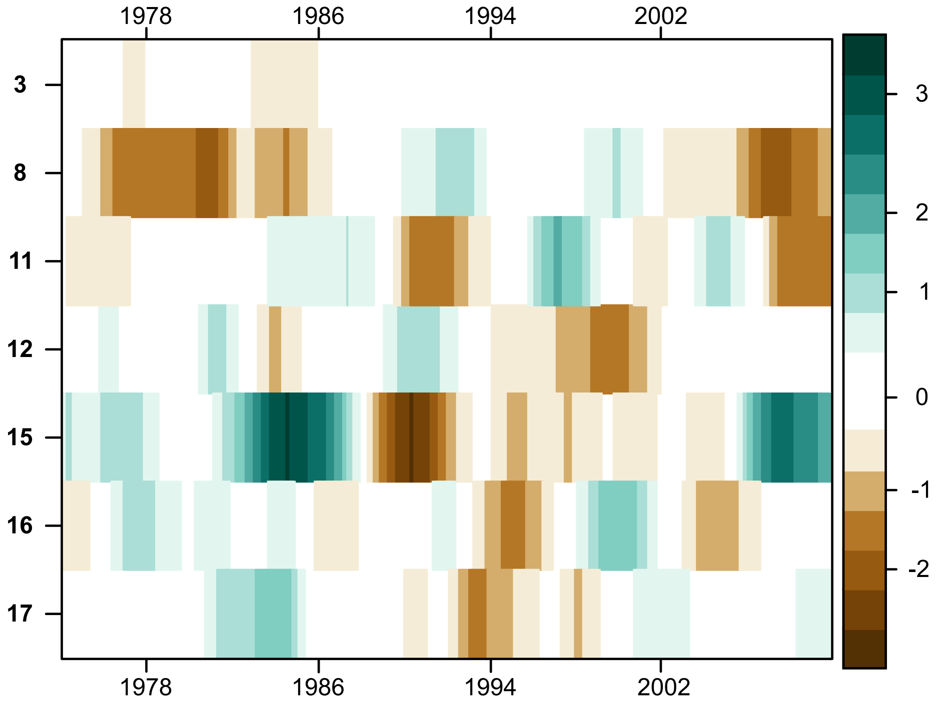


**Fig. SR2. Posterior $\log_2$ odds favoring a non-zero slope for the seven temporal eDNA clusters best described by the local linear trend (LLT) model.** Negative numbers and brown shading indicate a declining trend, whereas positive numbers and green shading indicate an increasing trend. A $\log_2$ odds of -1 means a decrease is twice as likely as an increase, or equivalently, *ca*. 67% of the posterior distribution supports a negative slope at time *t*.

*Climatic regressions*

Forecasts for eight clusters were improved by the climatic variables (fig. SR1; data S9). Best-fitting models that included the climatic regressors also tended to use the IRW trend model, which may suggest the mean-zero stochasticity expected under LL-like dynamics can be largely explained by climatic parameters.

We designed the climatic regressors to maximize non-redundant information, which can make biological interpretation more challenging. For example, the North Atlantic Oscillation (NAO) influences temperature and precipitation in northern Europe, particularly during the winter, but mechanisms linking its index (NAOI) and changes in eDNA abundances may not be obvious. To this end, we used density-preserving manifold approximation and projection (densUMAP) to reduce the full set of time-lagged climatic variables to seven latent axes of variation, corresponding broadly to: 1) precipitation, 2) water storage, 3) winter conditions, 4) warming trend, 5) seasonal transitions, 6) evapotranspiration, and the 7) Atlantic Multidecadal Oscillation (Supplementary Materials Section 7.4). This analysis enabled us to, for example, more specifically connect the variability of the NAOI (NAOI_sd ) to short-lag changes in frost and diurnal temperature range ('seasonal transitions') while running means of the NAOI closely covary with snowpack depth ('winter conditions'). We use these seven categories to summarize the regression results in fig. SR3 and marginal inclusion probabilities and coefficient estimates are shown in data S11.

Variables related to seasonal transitions were the most likely to be included in the regression models most (marginal inclusion probability $p(\zeta_k = 1) > 0.50$), followed by those related to precipitation and evapotranspiration (fig. SR3). Seasonal transitions predicted the abundances of all four plant-dominated clusters, along with the basidomycote cluster (fig. SR3). Mean precipitation and indices of extreme precipitation were included in three of the four fungal clusters (data S11), with lichens being the exception (fig. SR3). Evapotranspiration-related variables were also included in both plant and fungal cluster models, in addition to the sole bacterial cluster (comprising Enterobacteriales similar to *Erwinia*, *Rahnella*, and *Serratia*; data S6, S9), albeit with $p(\zeta_k = 1) = 0.47$. In general, regression components explained seasonal to cyclical variation, but we did not find clear evidence supporting climatic variables as predictors of long-term trends. Posterior draws of the trend and regression components of best-performing model for each cluster are shown in figs. SR4-20.

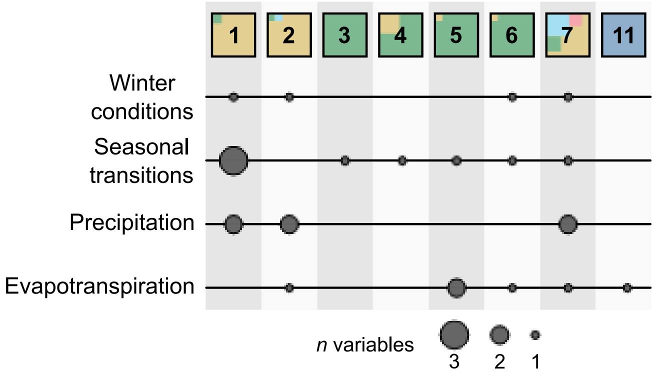


**Fig. SR3. Summary of climatic variables with marginal inclusion probabilities > 0.50† in the best-fitting eDNA abundance models.** Clusters are indicated by numbered boxes, colored by their kingdom-rank relative abundance composition as in Fig. SR1. The size of the circle indicates the number of variables on each climatic axis for a given model. † with the exception of Cluster 11, where p(ζ = 1) = 0.47 for the displayed variable.

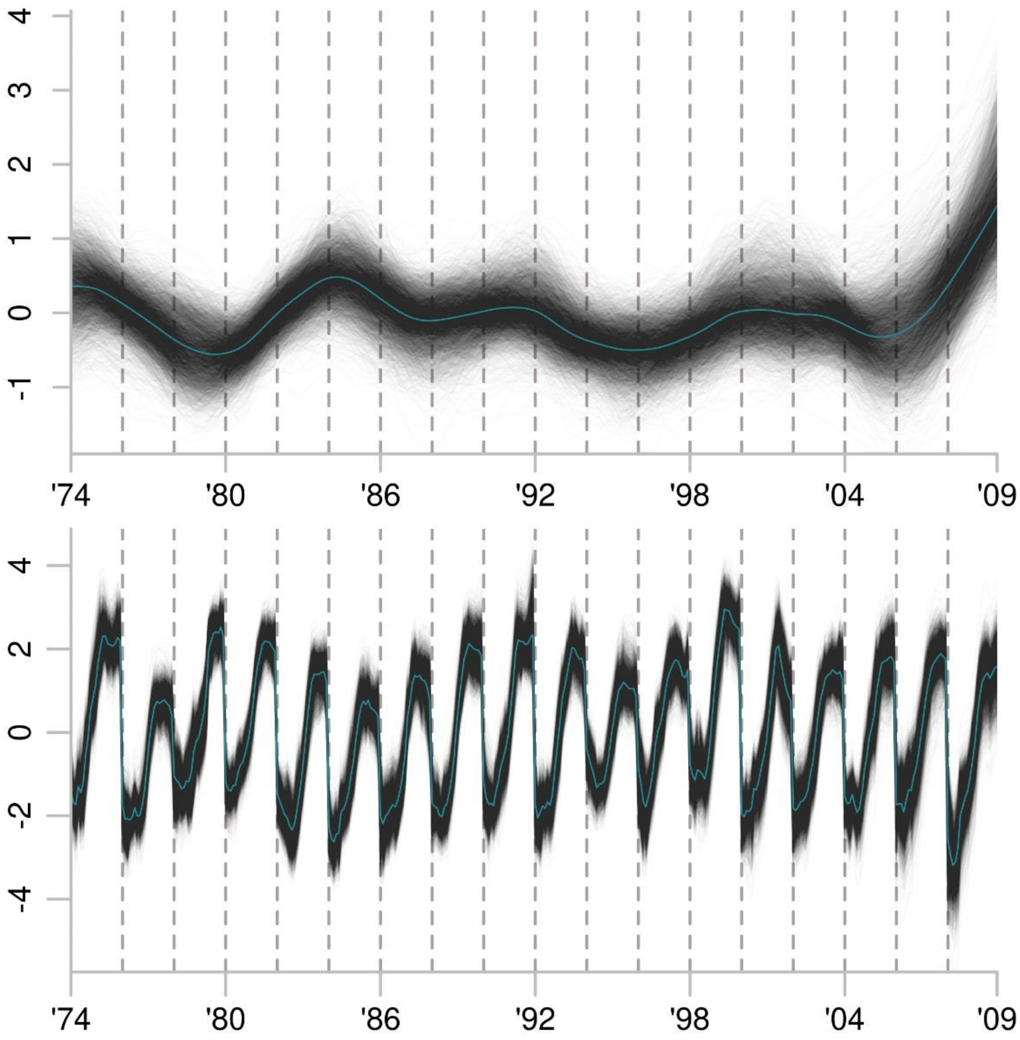


**Fig. SR4. Cluster1 best-fitting model: IRW-climate.** Greyscale lines show 5,000 posterior draws from the trend state component in the top panel and the regression component in bottom panel show. Teal lines indicate the median estimate. *Y*-axes are in pivot log-ratio coordinates. Years on the *x*-axes are even-numbered years from 1974 through 2008.

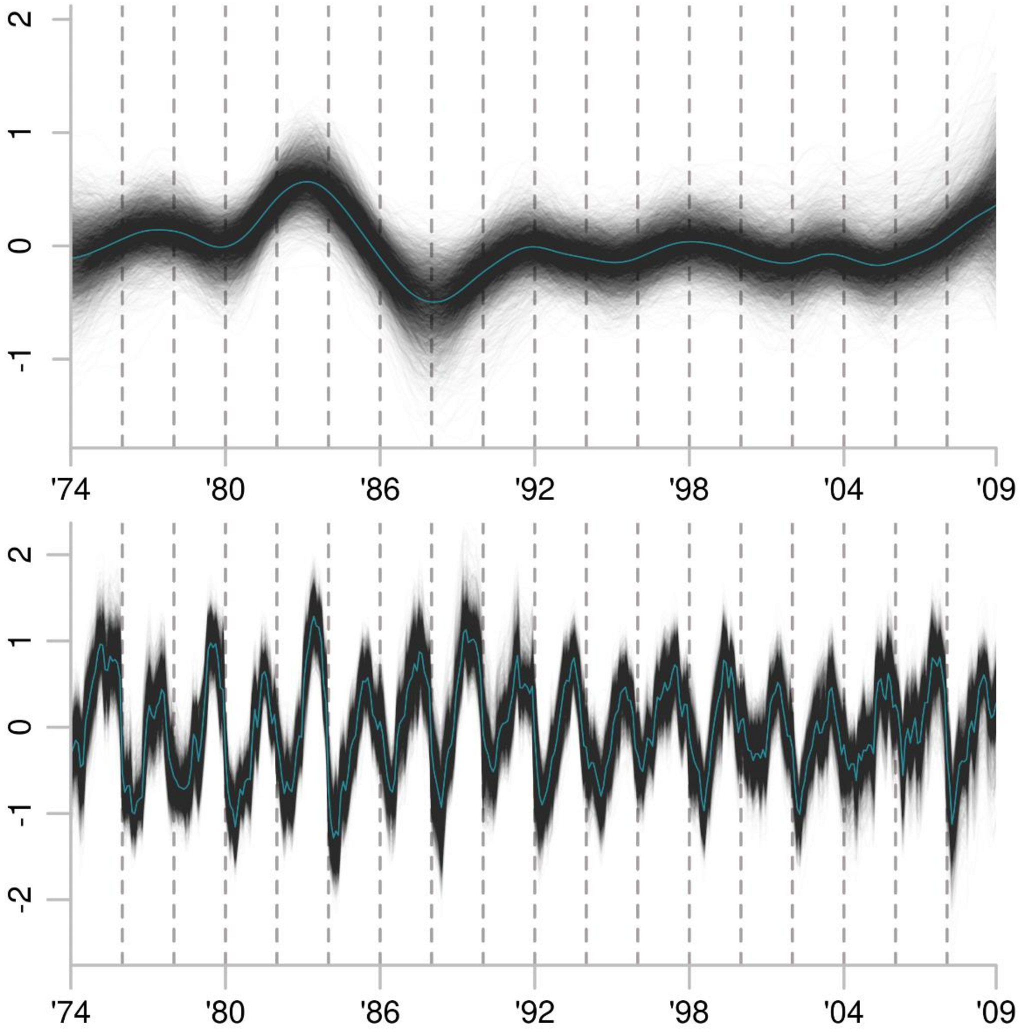


**Fig. SR5. Cluster2 best-fitting model: IRW-climate.** Greyscale lines show 5,000 posterior draws from the trend state component in the top panel and the regression component in bottom panel show. Teal lines indicate the median estimate. *Y*-axes are in pivot log-ratio coordinates. The *x*-axes show weekly estimates from the even-numbered years from 1974 through 2008.

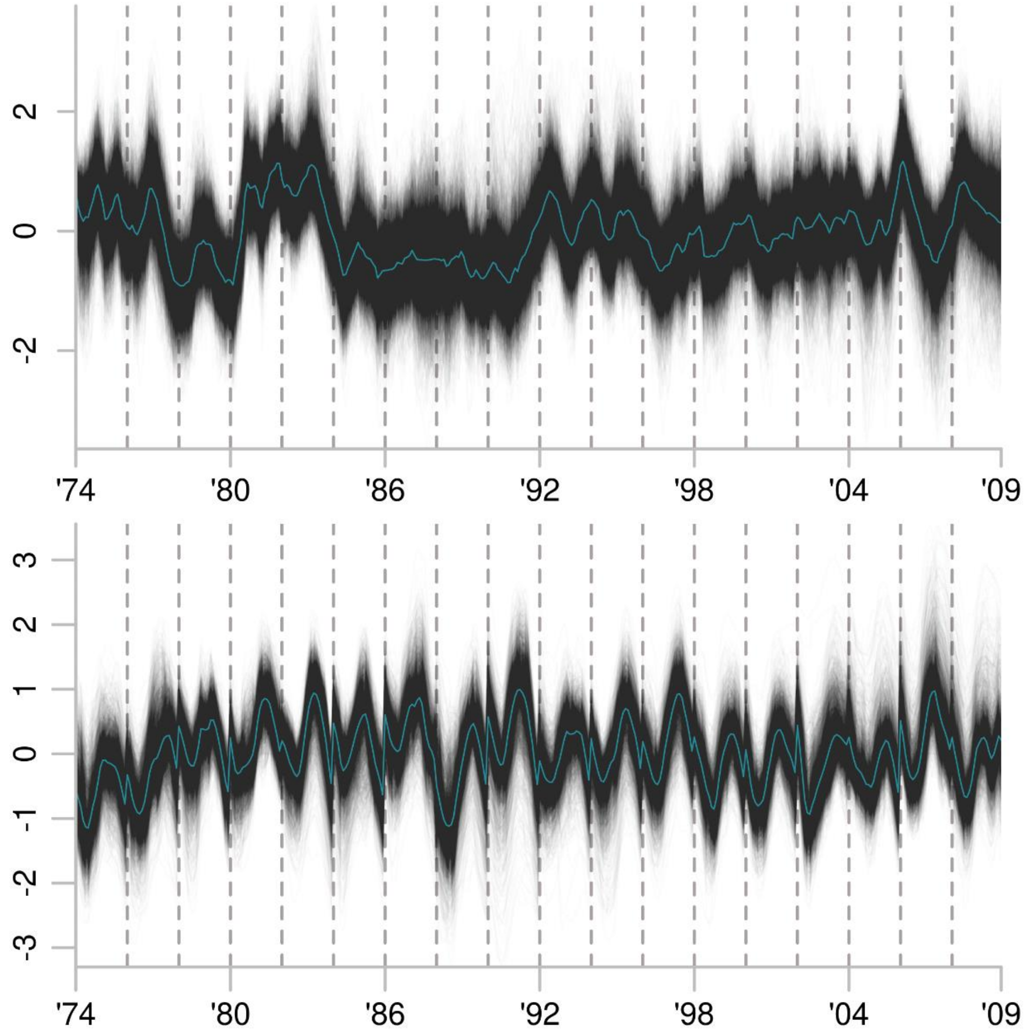


**Fig. SR6. Cluster3. best-fitting model: LLT-climate.** Greyscale lines show 5,000 posterior draws from the trend state component in the top panel and the regression component in bottom panel show. Teal lines indicate the median estimate. *Y*-axes are in pivot log-ratio coordinates. The *x*-axes show weekly estimates from the even-numbered years from 1974 through 2008.

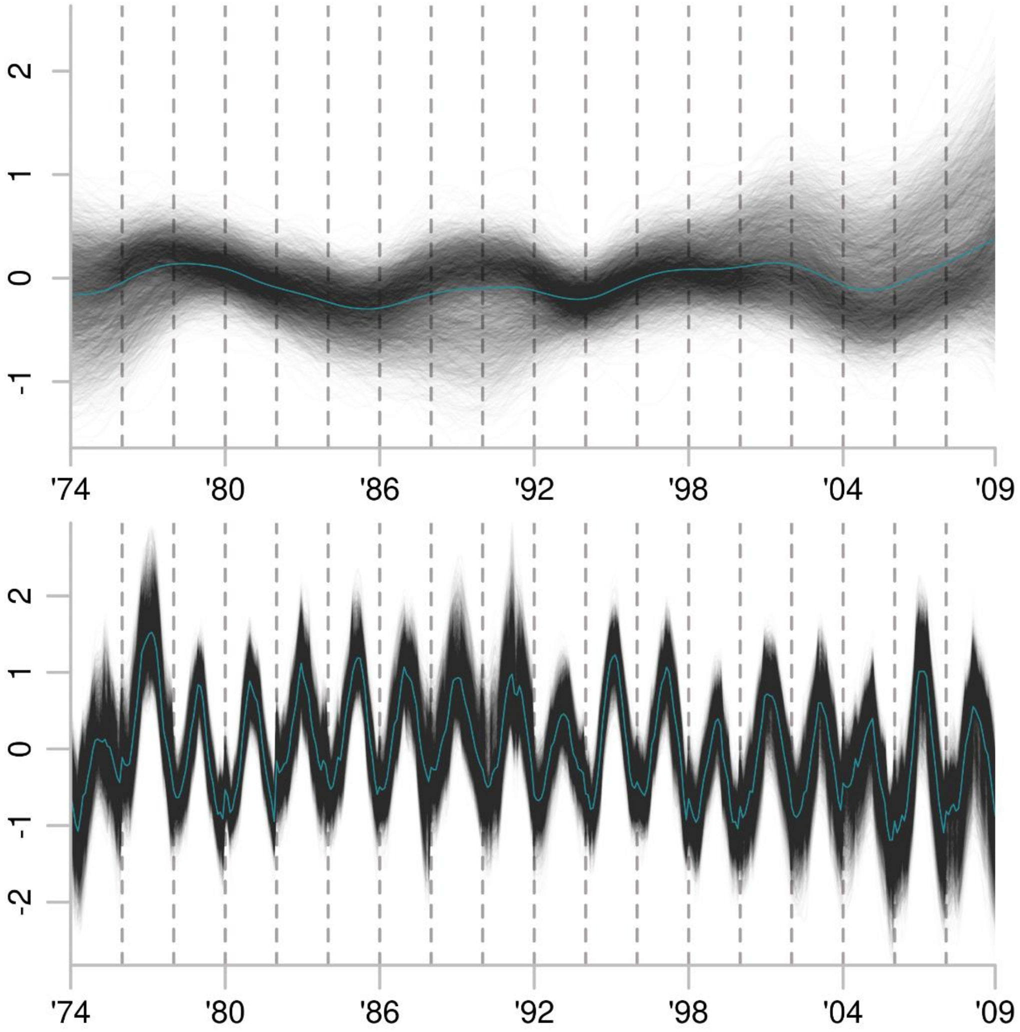


**Fig. SR7. Cluster4 best-fitting model: IRW-climate.** Greyscale lines show 5,000 posterior draws from the trend state component in the top panel and the regression component in bottom panel show. Teal lines indicate the median estimate. *Y*-axes are in pivot log-ratio coordinates. The *x*-axes show weekly estimates from the even-numbered years from 1974 through 2008.

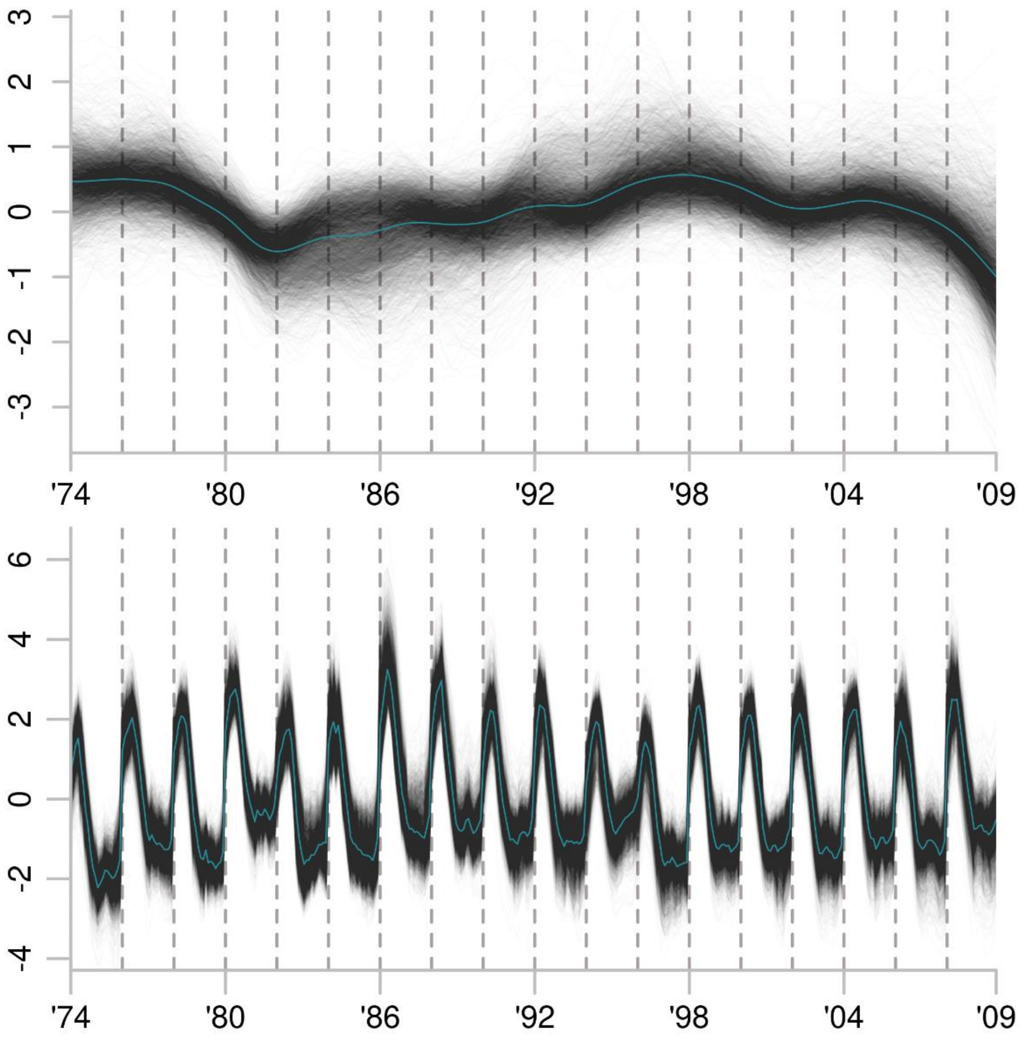


**Fig. SR8. Cluster5 best-fitting model: IRW-climate**. Greyscale lines show 5,000 posterior draws from the trend state component in the top panel and the regression component in bottom panel show. Teal lines indicate the median estimate. *Y*-axes are in pivot log-ratio coordinates. The *x*-axes show weekly estimates from the even-numbered years from 1974 through 2008.

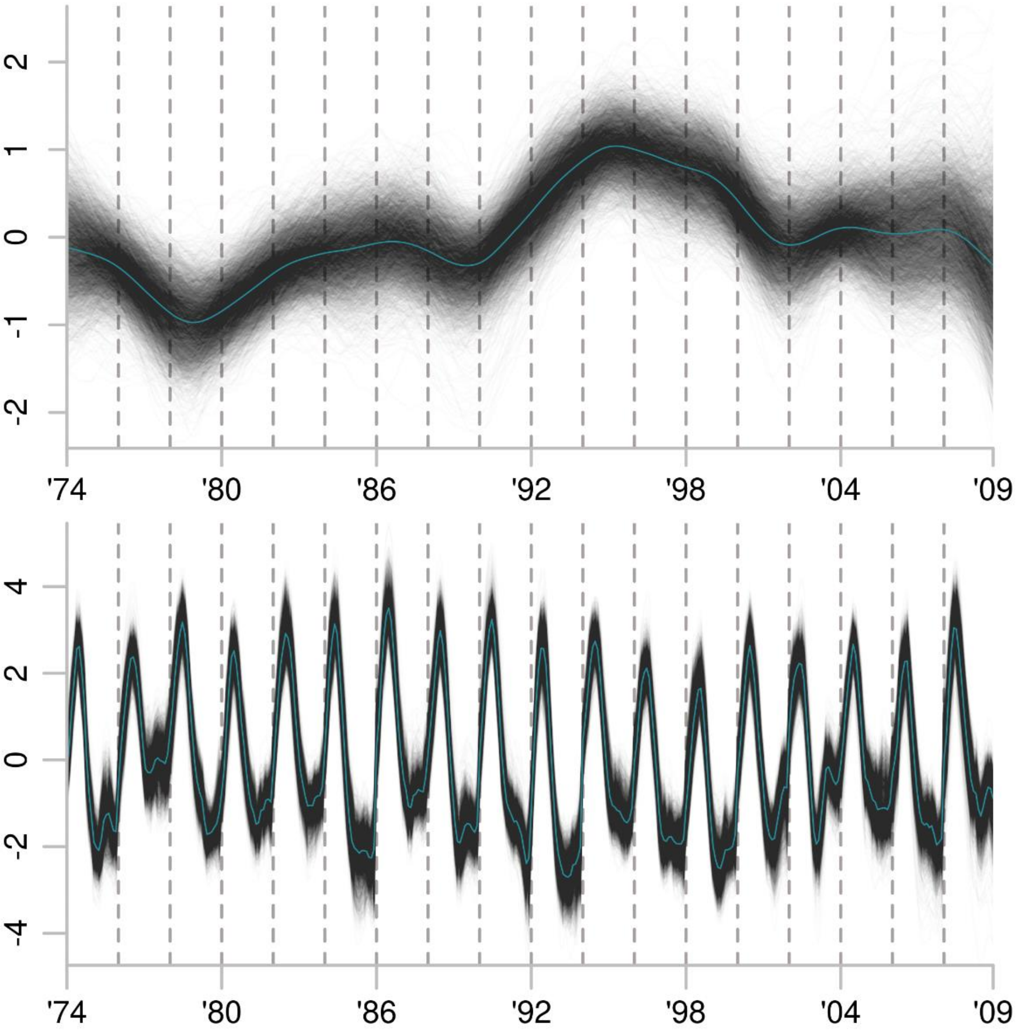


**Fig. SR9. Cluster6 best-fitting model: IRW-climate.** Greyscale show 5,000 posterior draws from the trend state component in the top panel and the regression component in bottom panel show. Teal lines indicate the median estimate. *Y*-axes are in pivot log-ratio coordinates. The *x*-axes show weekly estimates from the even-numbered years from 1974 through 2008.

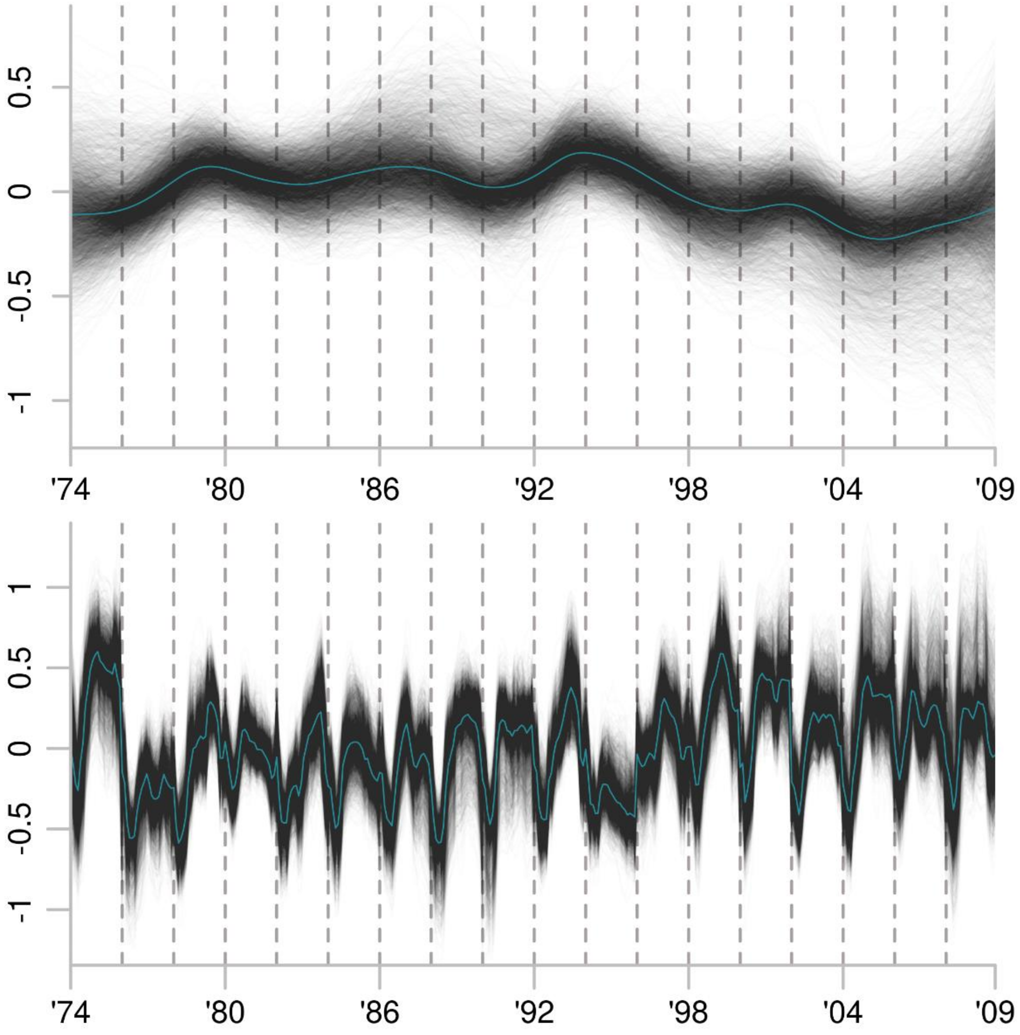


**Fig. SR10. Cluster7 best-fitting model: IRW-climate.** Greyscale show 5,000 posterior draws from the trend state component in the top panel and the regression component in bottom panel show. Teal lines indicate the median estimate. *Y*-axes are in pivot log-ratio coordinates. The *x*-axes show weekly estimates from the even-numbered years from 1974 through 2008.

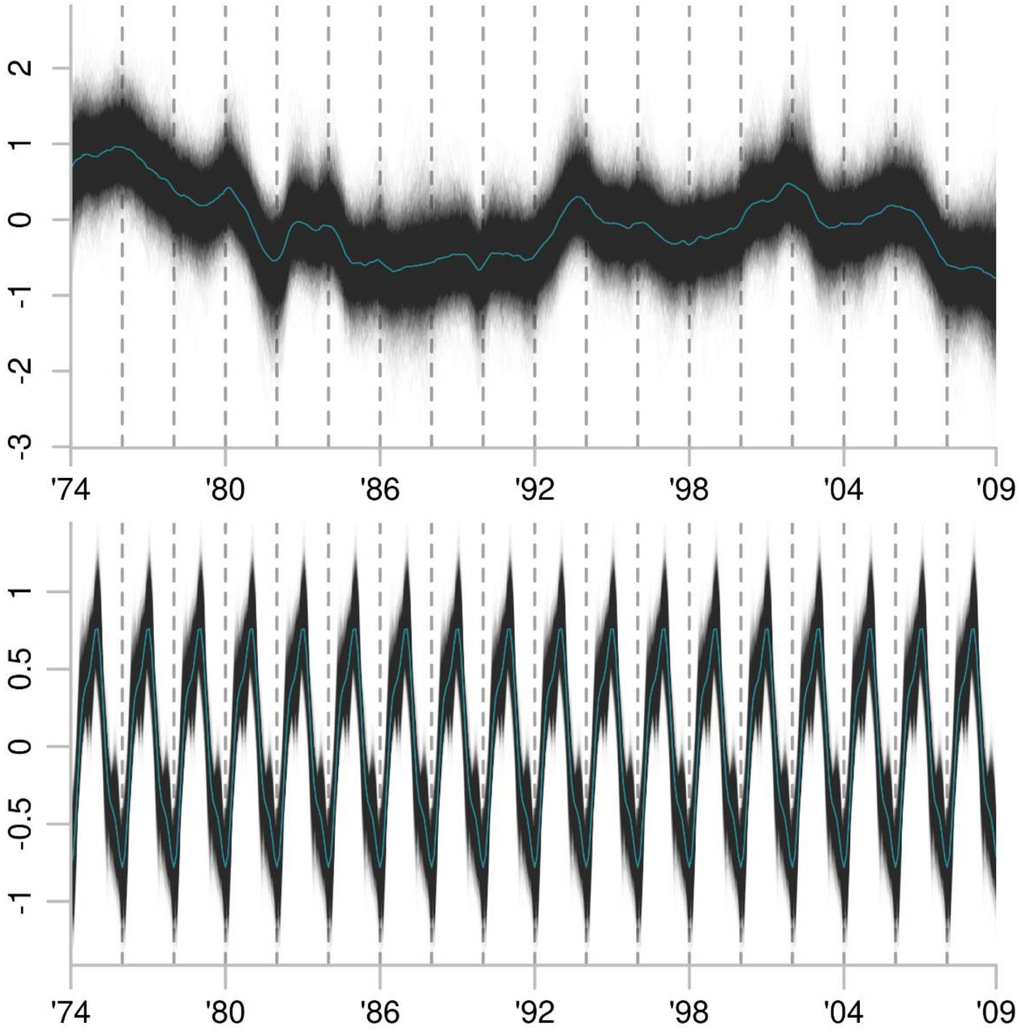


**Fig. SR11. Cluster8 best-fitting model: LLT-base.** Greyscale show 5,000 posterior draws from the trend state component in the top panel and the regression component in bottom panel show. Teal lines indicate the median estimate. *Y*-axes are in pivot log-ratio coordinates. The *x*-axes show weekly estimates from the even-numbered years from 1974 through 2008.

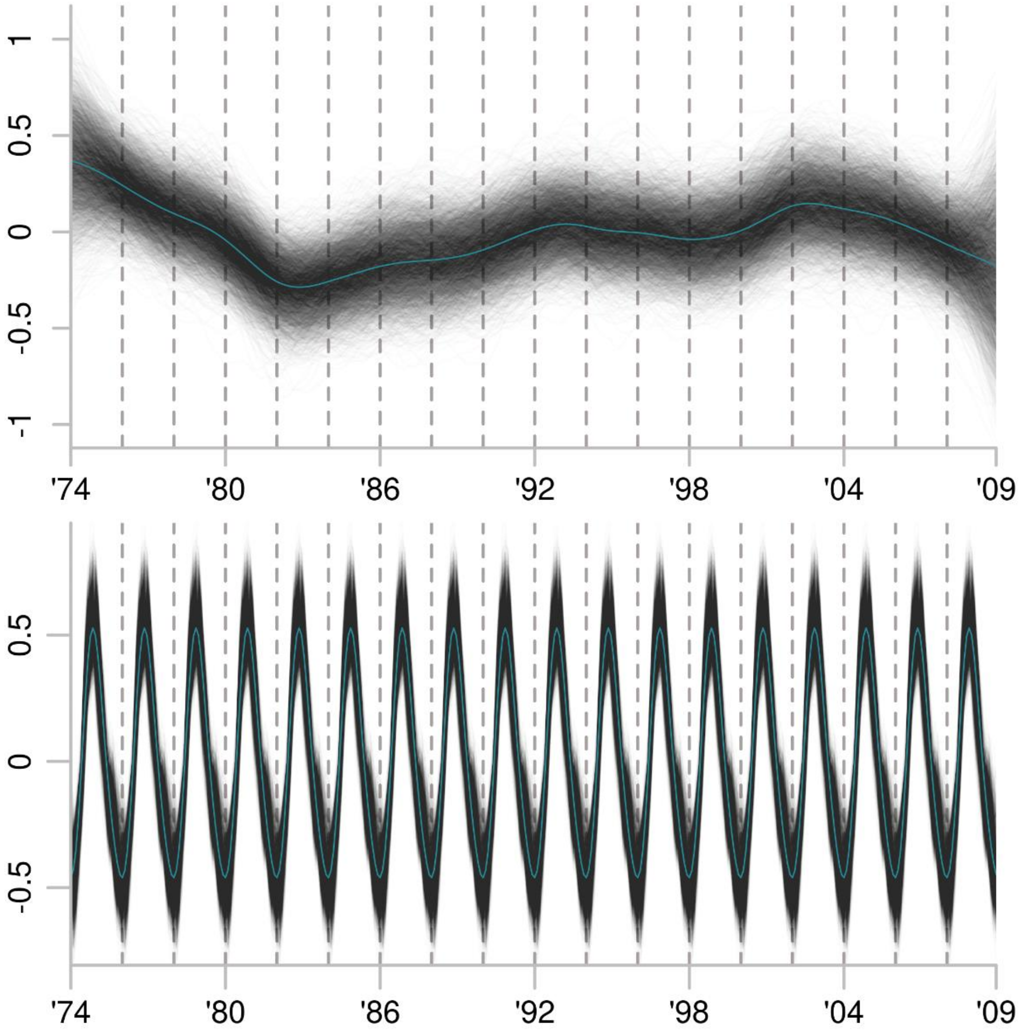


**Fig. SR12. Cluster9 best-fitting model: IRW-base.** Greyscale show 5,000 posterior draws from the trend state component in the top panel and the regression component in bottom panel show. Teal lines indicate the median estimate. *Y*-axes are in pivot log-ratio coordinates. The *x*-axes show weekly estimates from the even-numbered years from 1974 through 2008.

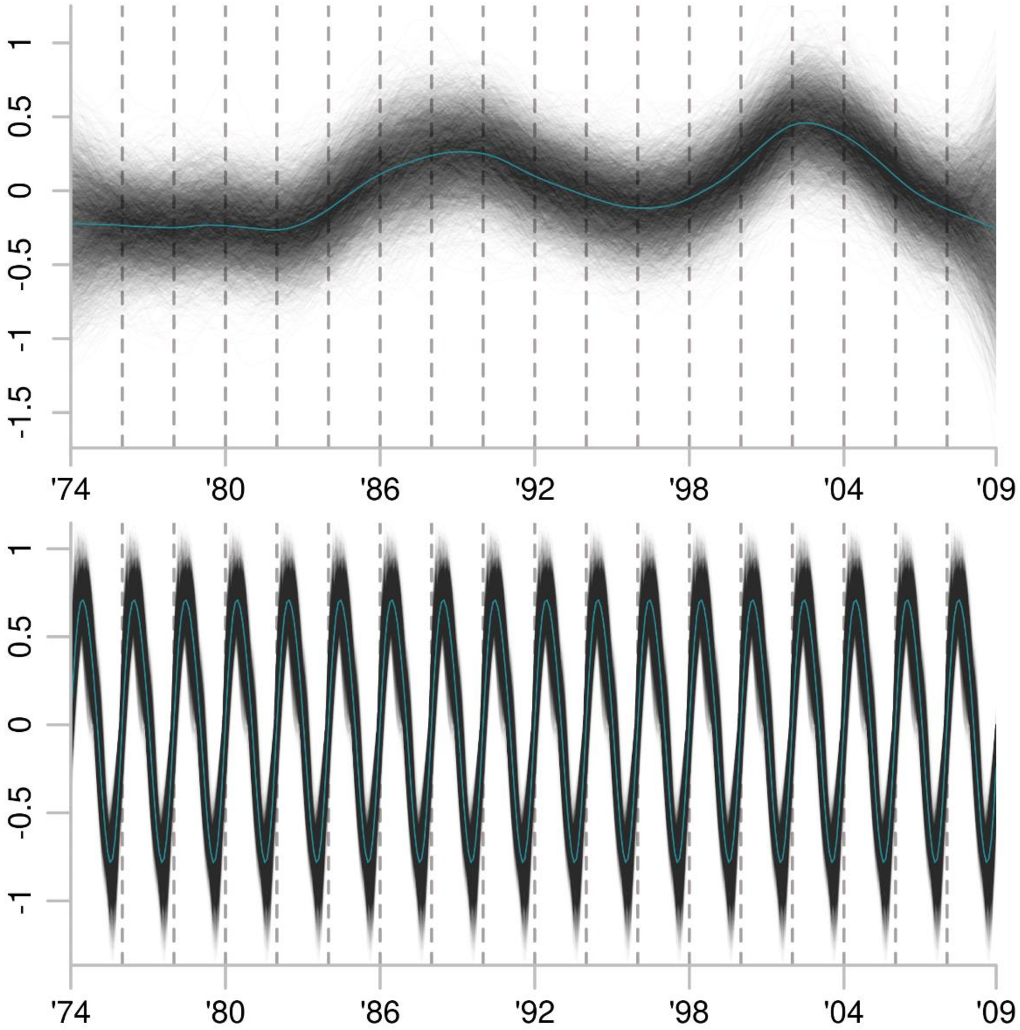


**Fig. SR13. Cluster10 best-fitting model: IRW-base.** Greyscale show 5,000 posterior draws from the trend state component in the top panel and the regression component in bottom panel show. Teal lines indicate the median estimate. *Y*-axes are in pivot log-ratio coordinates. The *x*-axes show weekly estimates from the even-numbered years from 1974 through 2008.

.

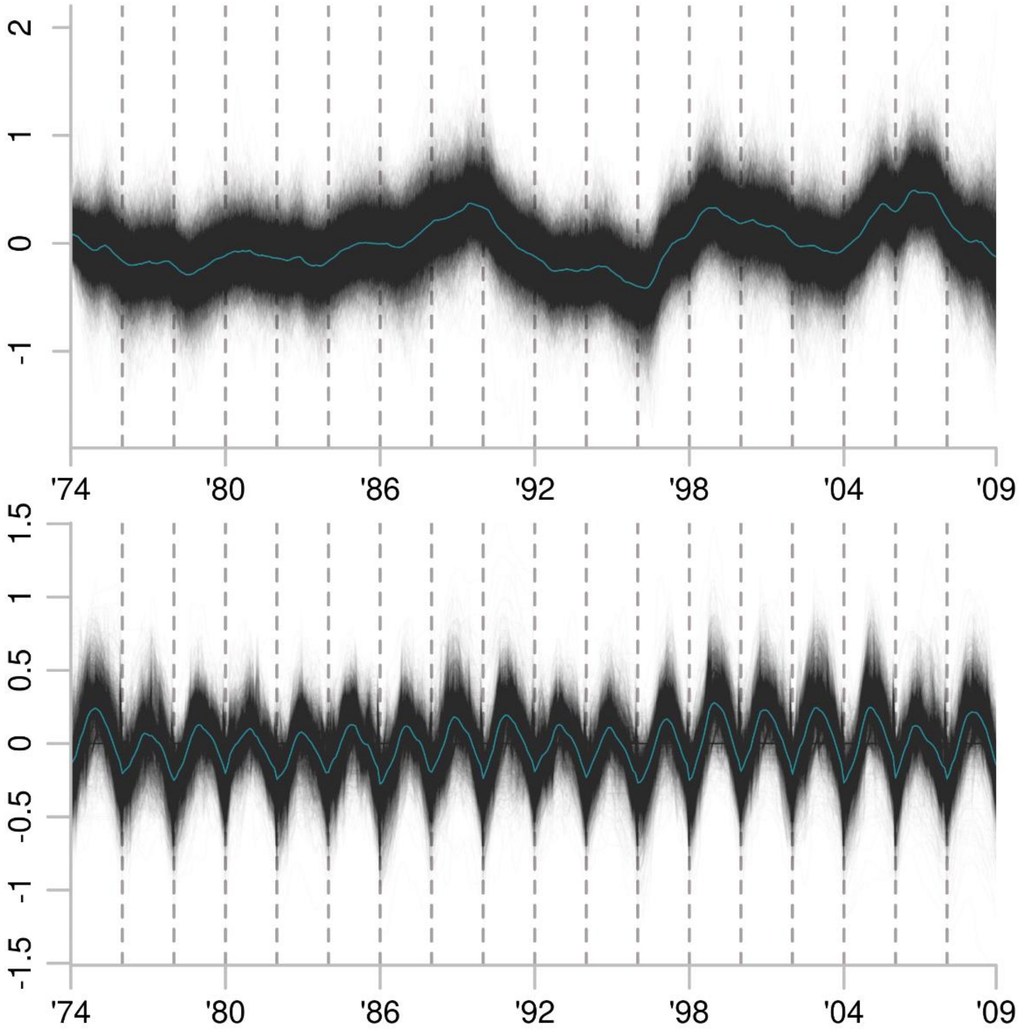


**Fig. SR14. Cluster11 best-fitting model: LLT-climate.** Greyscale show 5,000 posterior draws from the trend state component in the top panel and the regression component in bottom panel show. Teal lines indicate the median estimate. *Y*-axes are in pivot log-ratio coordinates. The *x*-axes show weekly estimates from the even-numbered years from 1974 through 2008.

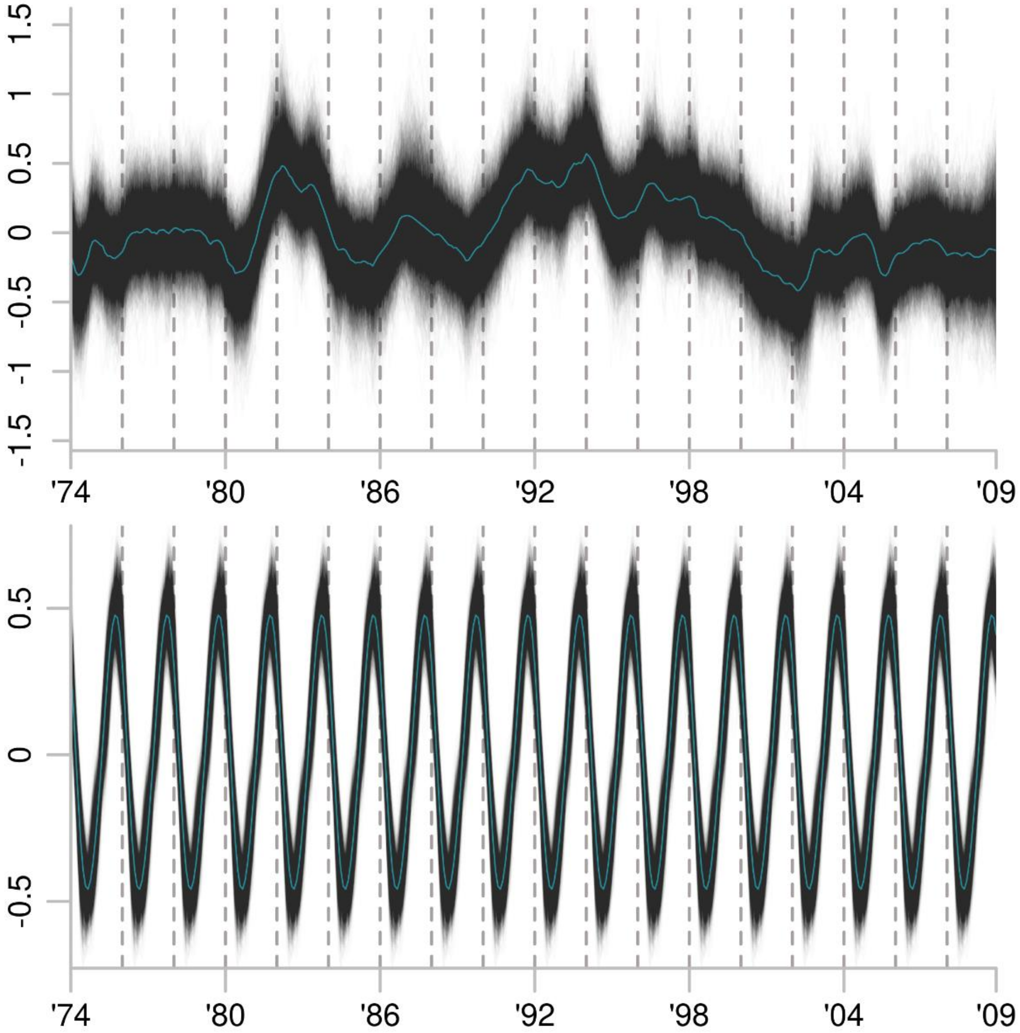


**Fig. SR15*. *Cluster12 best-fitting model: LLT-base.** Greyscale show 5,000 posterior draws from the trend state component in the top panel and the regression component in bottom panel show. Teal lines indicate the median estimate. *Y*-axes are in pivot log-ratio coordinates The *x*-axes show weekly estimates from the even-numbered years from 1974 through 2008.

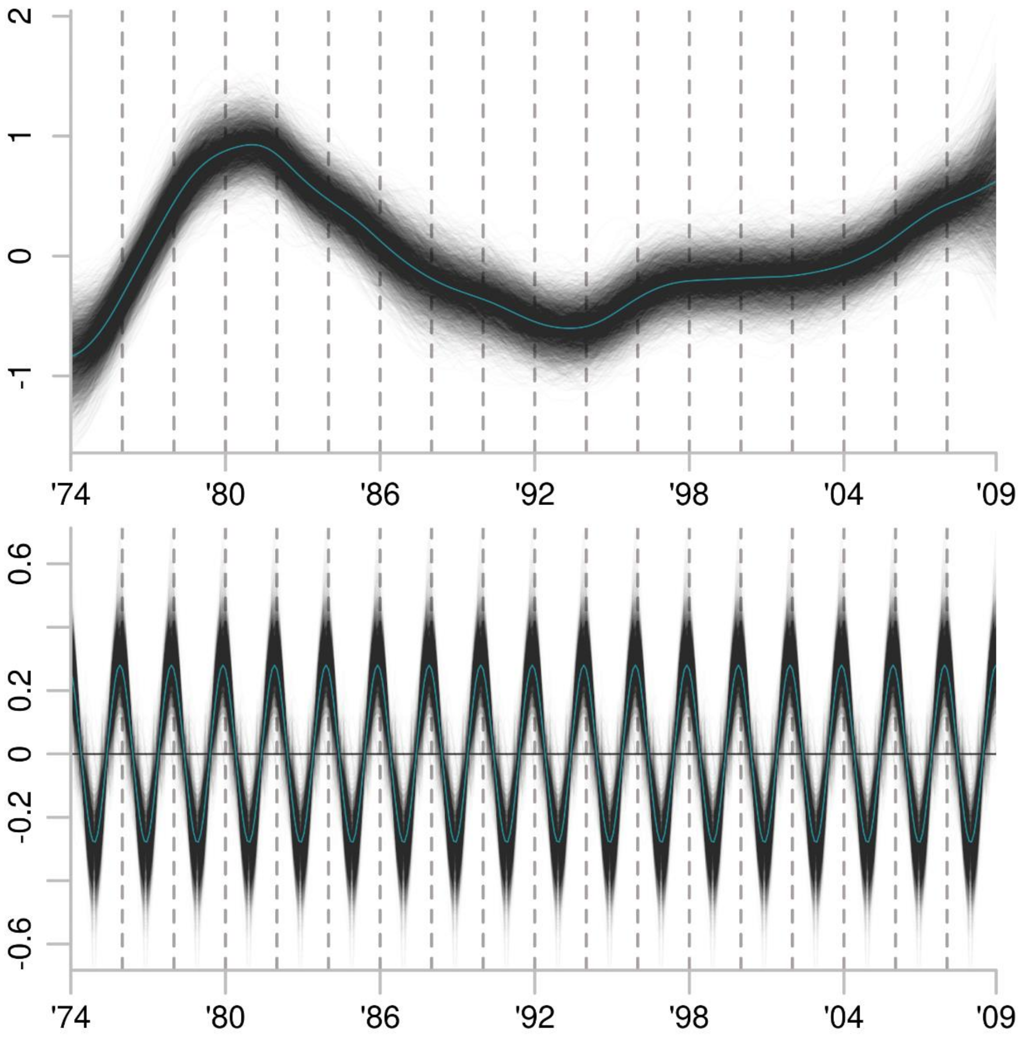


**Fig. SR16. Cluster13 best-fitting model: IRW-base.** Greyscale show 5,000 posterior draws from the trend state component in the top panel and the regression component in bottom panel show. Teal lines indicate the median estimate. *Y*-axes are in pivot log-ratio coordinates. The *x*-axes show weekly estimates from the even-numbered years from 1974 through 2008.

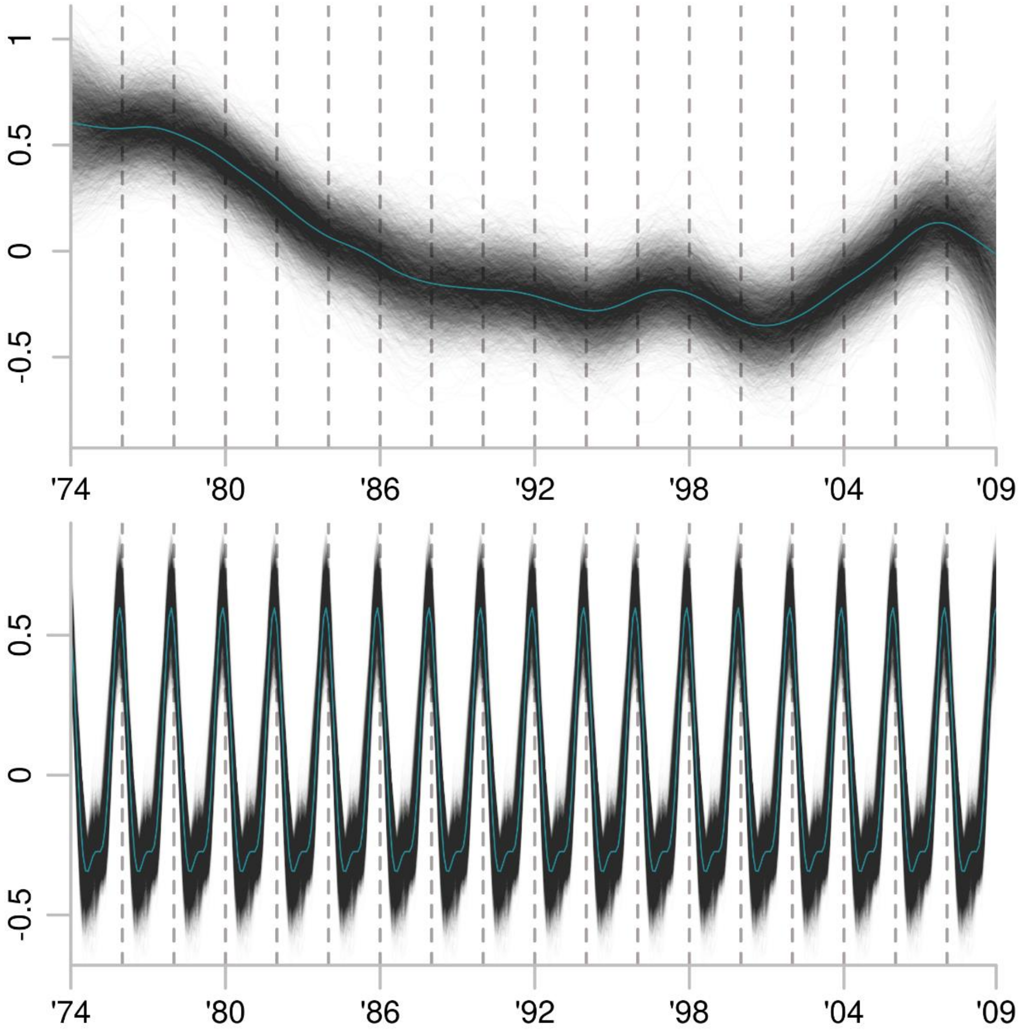


**Fig. SR17. Cluster14 best-fitting model: IRW-base.** Greyscale show 5,000 posterior draws from the trend state component in the top panel and the regression component in bottom panel show. Teal lines indicate the median estimate. *Y*-axes are in pivot log-ratio coordinates. The *x*-axes show weekly estimates from the even-numbered years from 1974 through 2008.

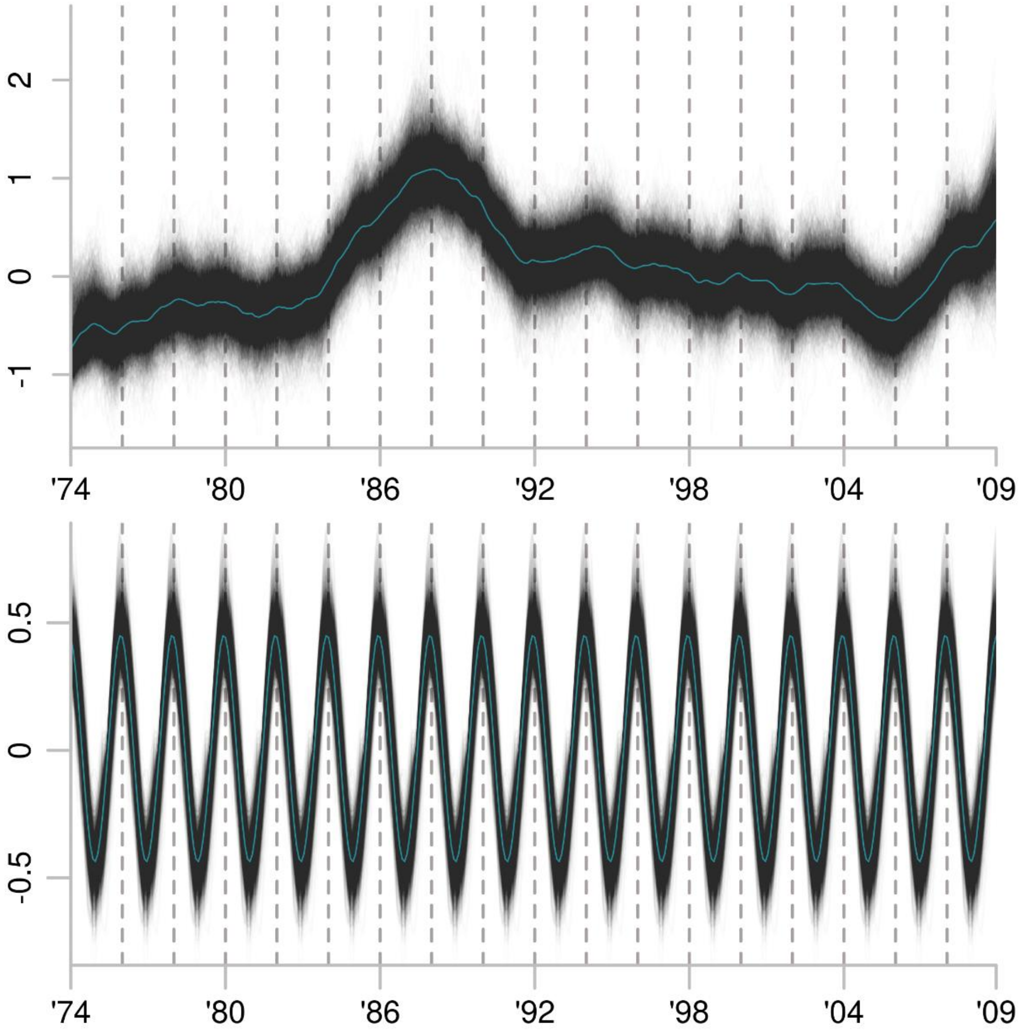


**Fig. SR18. Cluster15 best-fitting model: LLT-base.** Greyscale show 5,000 posterior draws from the trend state component in the top panel and the regression component in bottom panel show. Teal lines indicate the median estimate. *Y*-axes are in pivot log-ratio coordinates. The *x*-axes show weekly estimates from the even-numbered years from 1974 through 2008.

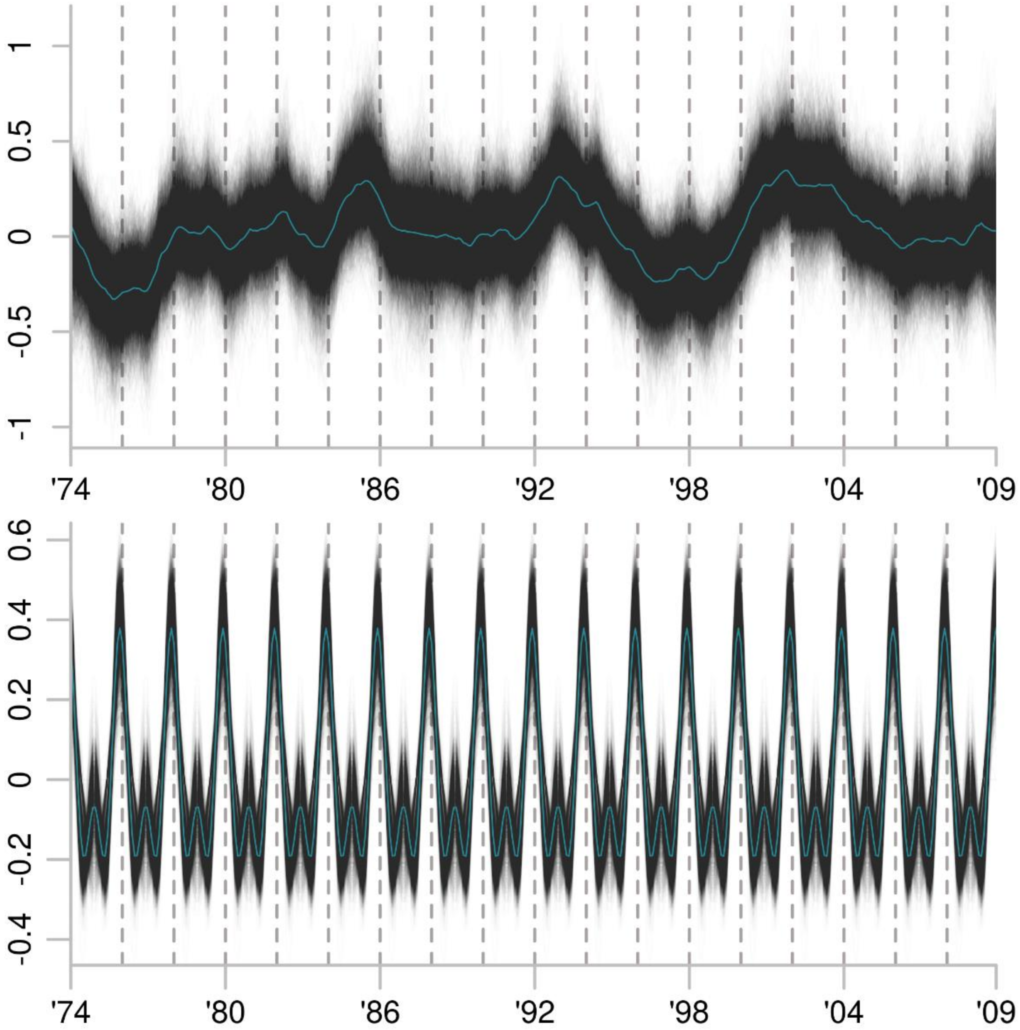


**Fig. SR19. Cluster16 best-fitting model: LLT-base.** Greyscale show 5,000 posterior draws from the trend state component in the top panel and the regression component in bottom panel show. Teal lines indicate the median estimate. *Y*-axes are in pivot log-ratio coordinates. The *x*-axes show weekly estimates from the even-numbered years from 1974 through 2008.

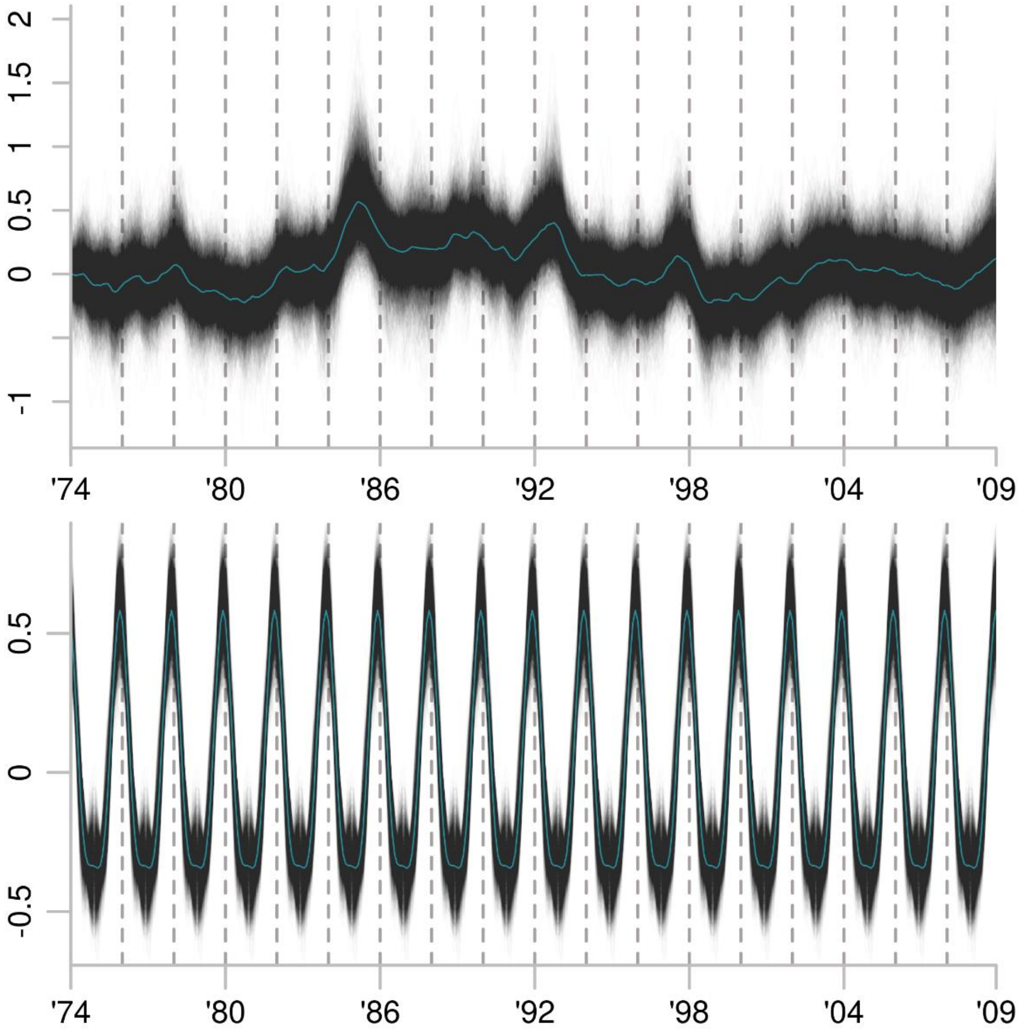


**Fig. SR20. Cluster17 best-fitting model: LLT-base.** Greyscale show 5,000 posterior draws from the trend state component in the top panel and the regression component in bottom panel show. Teal lines indicate the median estimate. *Y*-axes are in pivot log-ratio coordinates. The *x*-axes show weekly estimates from the even-numbered years from 1974 through 2008.

**Bayesian state-space time series models of α-, β-, and γ-diversity**

We estimated diversity components for three groupings of the eDNA community: 1) the total 2,739-genera dataset, 2) the 1,558 eukaryotic genera, and 3) the total dataset with the exclusion of the 208 genera in cluster14, which comprised genera associated with animal microbiomes, including humans and domestic animals (data S5, S6). For each subset, we calculated α-, β-, and γ-diversity of order $q$ = 1, 2, and 3. Higher orders increasingly emphasize the contributions of abundant taxa, and $q$ = 1 weighs taxa proportionally to their relative abundances.

The best-performing model included the climatic regressor matrix for all 27 diversity metrics (data S9). Twenty-one used the IRW trend model and the LLT and IRW trends were indistinguishable based on ΔELPD in the remaining six (data S9). We considered the IRW trend to be the ‘best’ model for these as well because the LLT model contains an additional free parameter.

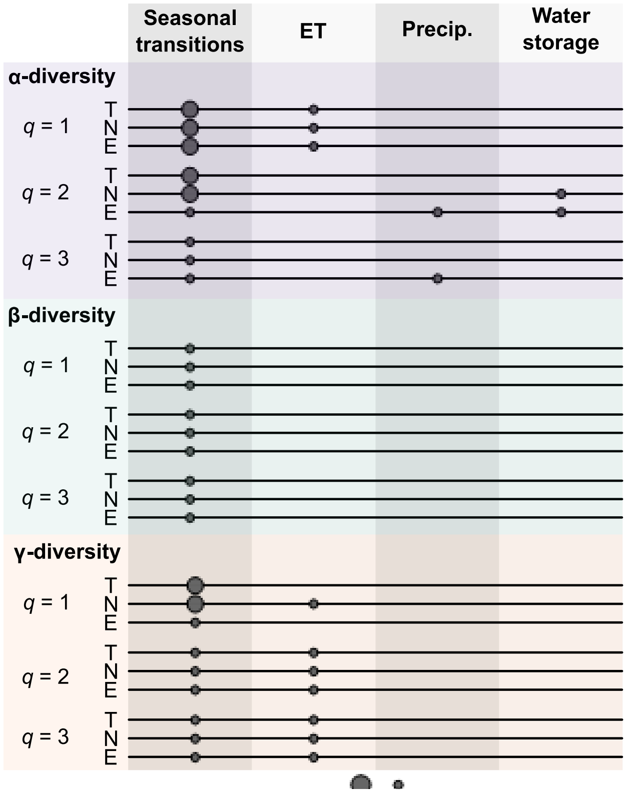


**Fig. SR21. Summary of climatic variables with marginal inclusion probabilities > 0.50† in the best-fitting diversity models.** ’T’ denotes the diversity of the total dataset, ‘N’ the fraction exclusive of cluster14 genera, and ‘E’ the diversity of eukaryotes. Circle sizes indicate the number of variables on each climatic axis in a given model.
† with the exception of q = 1 β-diversity, where p(ζ = 1) > 0.40 for the displayed variables.

Although all diversity models included the climatic regressor matrix, few individual variables had high marginal posterior inclusion probabilities (fig. SR21). Variables with $p(\zeta_k = 1) > 0.50$, however, tended to be included more frequently in the diversity ($\hat{p}$ = 0.85) than the cluster abundance models ($\hat{p}$ = 0.69; data S11). As in the abundance models, seasonal transitions were the most consistent predictors of diversity trends but precipitation-related variables contributed relatively less predictive power (*cf.* figs. SR3 and SR21). Climatic regression predicted seasonal to cyclical variation but did not provide a clear explanation for any sustained changes in slope (figs. SR23-SR31).

Trends for a given diversity metric were similar across the three datasets (fig. SR22). Diversity metrics calculated for eukaryotic genera tended to change more slowly over time, with fewer of the rapid but transient periods of anomalous diversity detected for the total community. Eukaryotes also experienced smaller losses of β- and γ-diversity than the community as a whole (fig. SR22). This is most clearly evident in β-diversity of order $q = 2$ and 3, which declined by 18% and 26% from 1974-1988 to 2000-2008 in eukaryotes, compared to 33% and 38% in the total community, respectively (figs. SR 27-28).

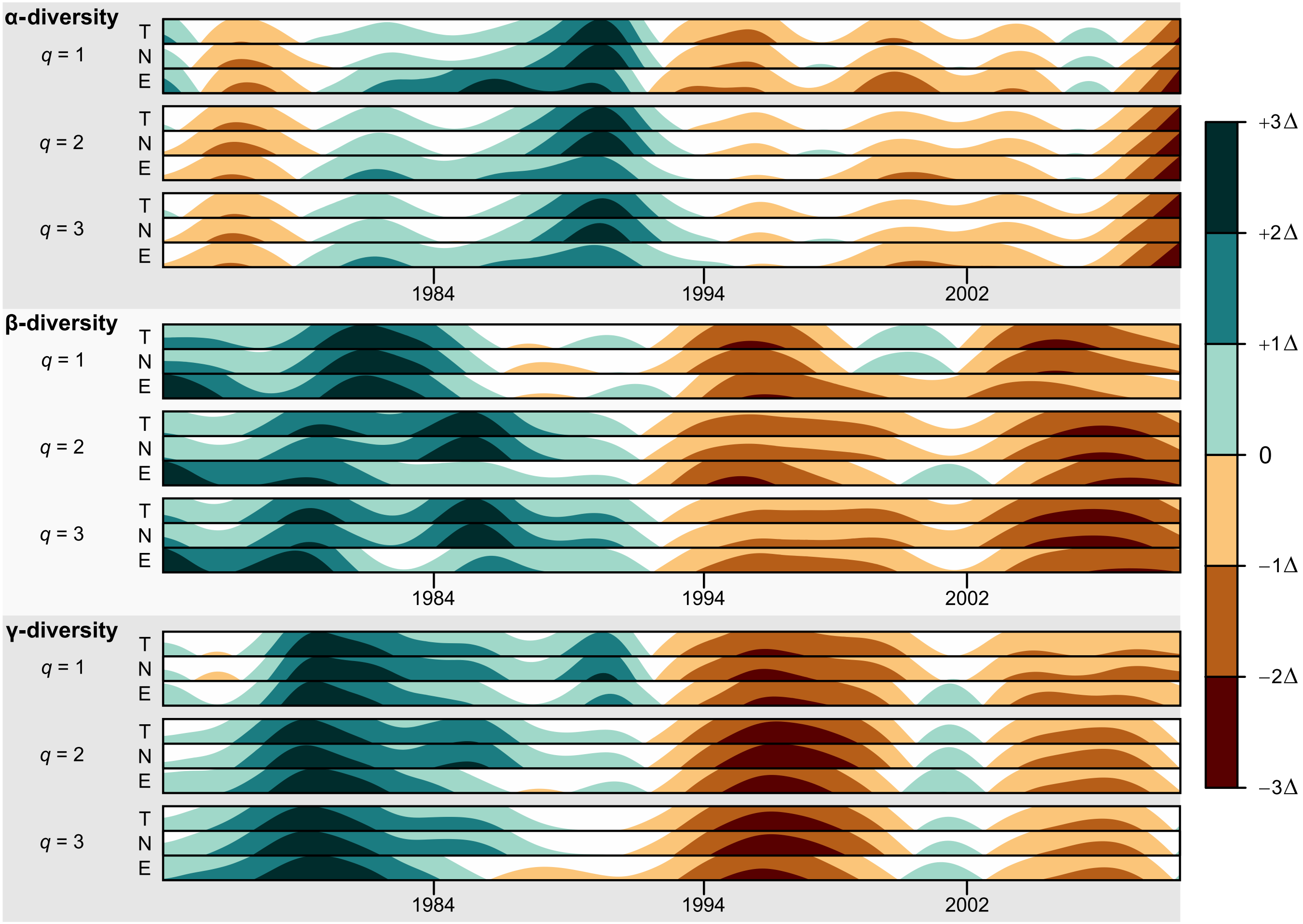


**Fig. SR22. Diversity trend estimates**. Each time series is centered, scaled, and compressed to a uniform *y*-axis so that the magnitudes of change at a given point in time are represented by color. Darker colors indicate larger positive (green) and negative (orange) deviations from the mean.

Different orders of the diversity components also resulted in similar trends (fig. SR22). However, higher-order ($q = 2, 3$) β-diversities indicated a larger and more rapid shift in community composition in the early 1990s (figs. SR22, SR26-28), while the opposite held true for γ-diversity (figs. S29-31). Higher-order ($q$ = 2, 3) γ-diversities also differed in the strength of the 2000-2002 local peak and by their strongly positive trajectory in 2008. In contrast, posterior estimates of first-order γ-diversities in 2008 almost equally supported a decline as an increase (figs. SR29-31). The larger shifts in community composition indicated by high-order β-diversities are consistent with the abundance changes found among dominant taxa, such as *Pinus* and *Pleurozium*. However, the differences among γ- diversity orders are more difficult to interpret because they occurred primarily during three years near the end of the time series. Trends for many of the eDNA clusters suggest a change in trajectory in 2008, particularly the highly abundant basidiomycote- and spruce-dominated clusters (figs. SR4, SR8; clusters 1 and 5, respectively), which would be reflected by γ-diversity. However, extending the time series is likely necessarily to determine if these 2008 patterns were a prelude to a similar shift in community diversity as in the early 1990s or were a part of a more transient event like in 2000-2002

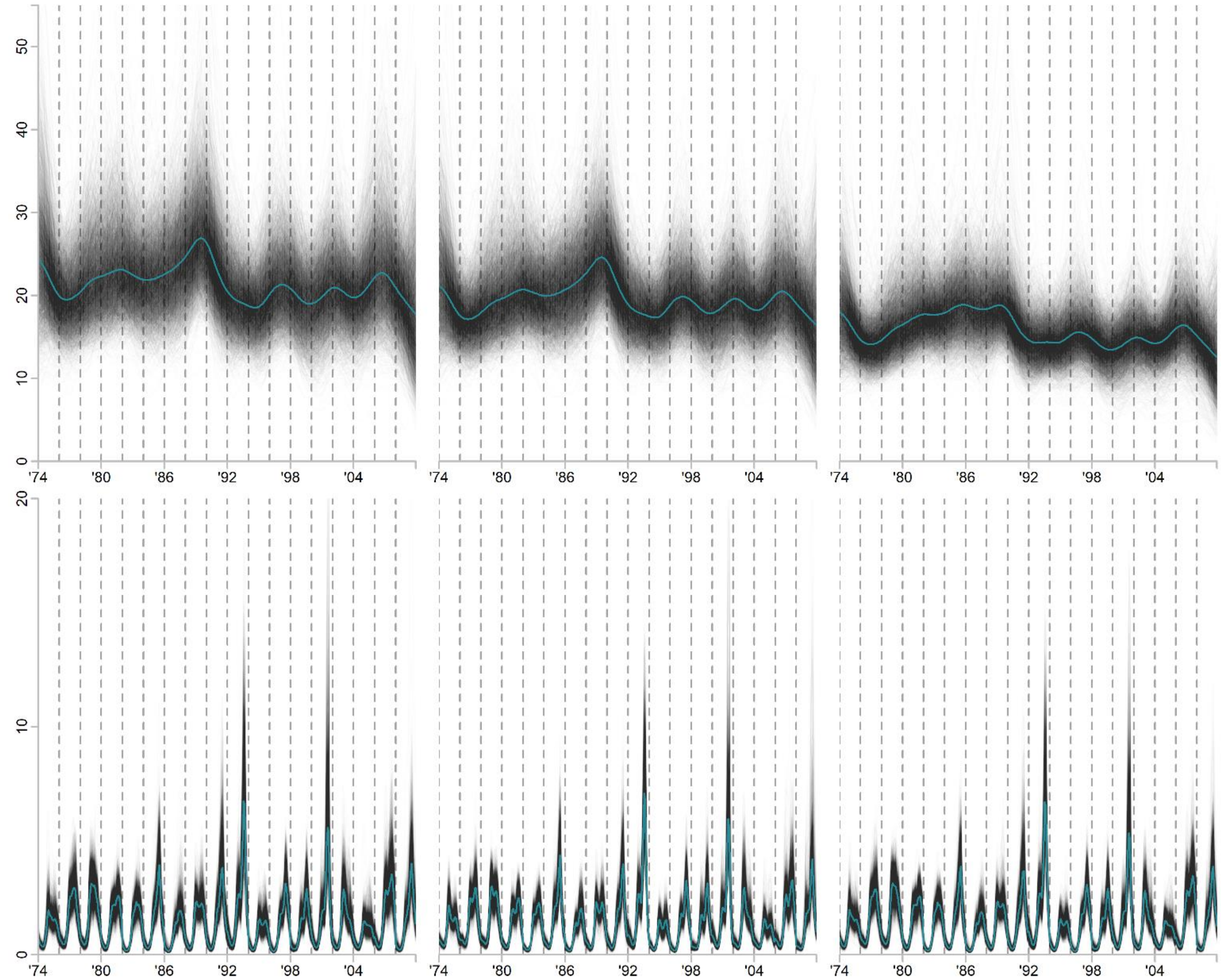


**Fig. SR23. IRW-climate models of α-diversity *q* = 1.** Columns from left to right show the model for 1) the total community, 2) the total community excluding the 208 genera in cluster14, and 3) only eukaryote genera. Greyscale lines show 5,000 posterior draws from the trend state component in the top panel and the regression component in bottom panel show. Teal lines indicate the median estimate. *Y*-axes are in units of effective number of genera. The *x*-axes show weekly estimates from the even-numbered years from 1974 through 2008.

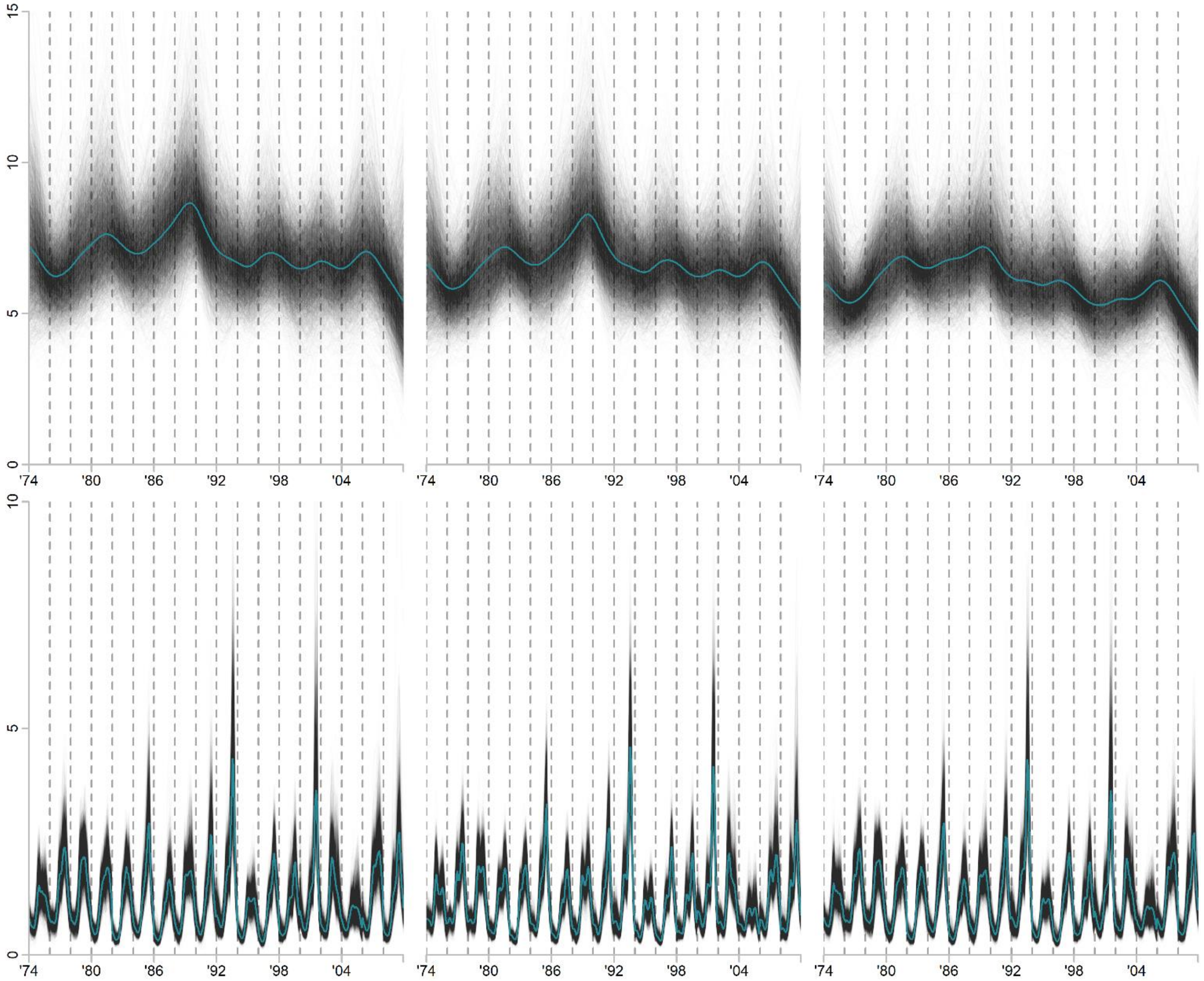


**Fig. SR24. IRW-climate models of α-diversity *q* = 2. .** Columns from left to right show the model for 1) the total community, 2) the total community excluding the 208 genera in cluster14, and 3) only eukaryote genera. Greyscale lines show 5,000 posterior draws from the trend state component in the top panel and the regression component in bottom panel show. Teal lines indicate the median estimate. *Y*-axes are in units of effective number of genera. The *x*-axes show weekly estimates from the even-numbered years from 1974 through 2008.

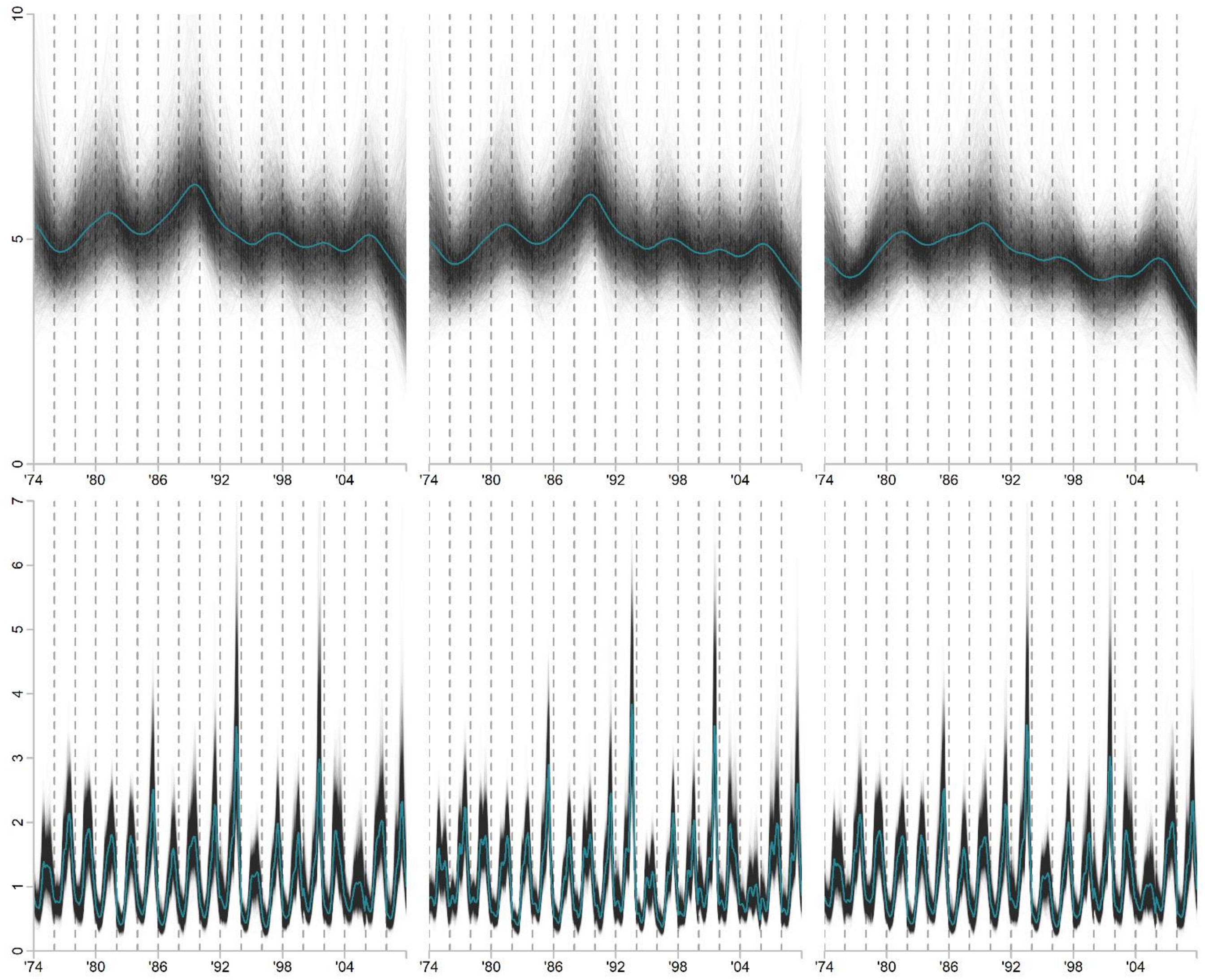


**Fig. SR25. IRW-climate models of α-diversity *q* = 3.** Columns from left to right show the model for 1) the total community, 2) the total community excluding the 208 genera in cluster14, and 3) only eukaryote genera. Greyscale lines show 5,000 posterior draws from the trend state component in the top panel and the regression component in bottom panel show. Teal lines indicate the median estimate. *Y*-axes are in units of effective number of genera. The *x*-axes show weekly estimates from the even-numbered years from 1974 through 2008 and are separated by dashed lines.

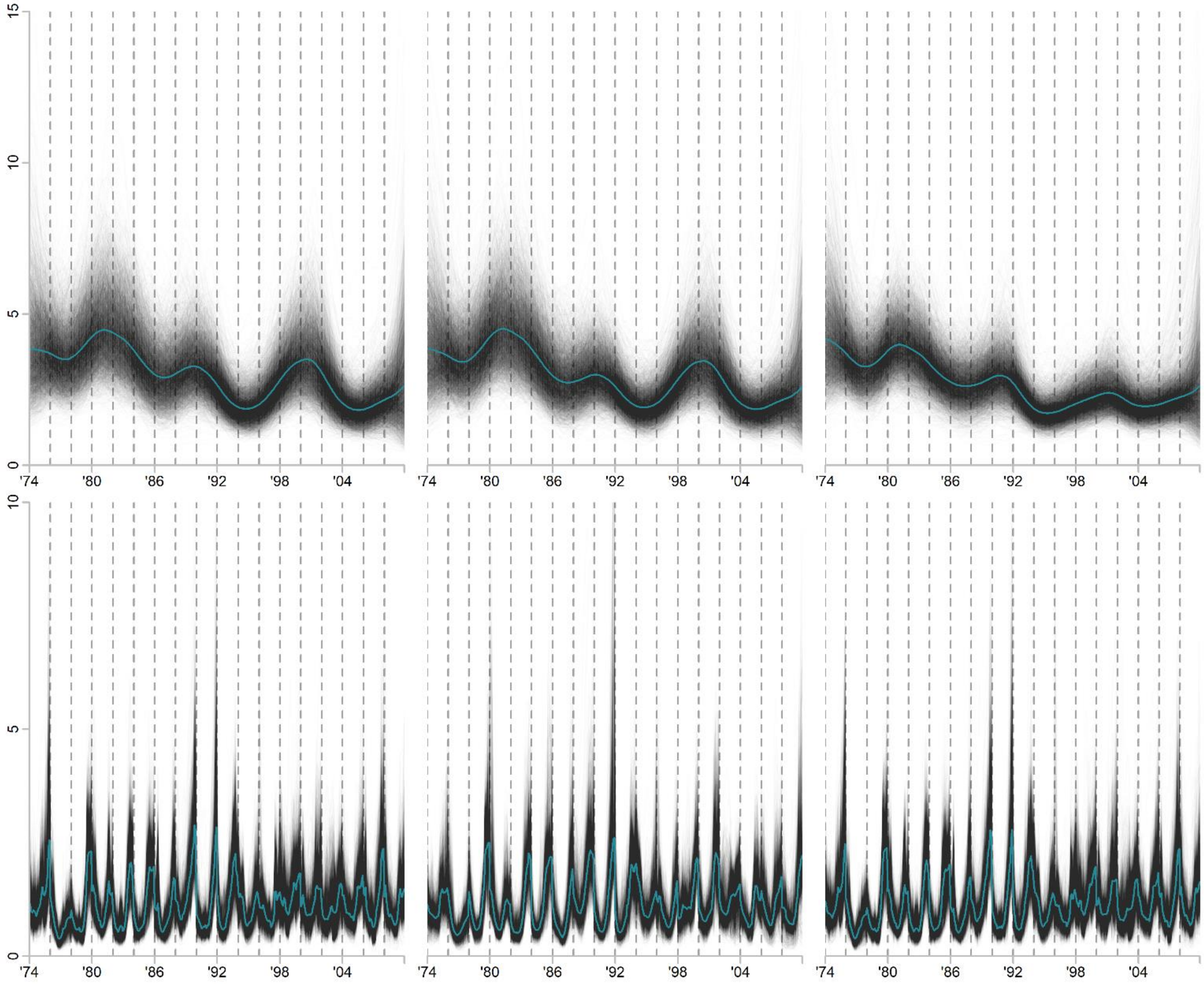


**Fig. SR26. IRW-climate models of β-diversity *q* = 1*.*** Columns from left to right show the model for 1) the total community, 2) the total community excluding the 208 genera in cluster14, and 3) only eukaryote genera. Greyscale lines show 5,000 posterior draws from the trend state component in the top panel and the regression component in bottom panel show. Teal lines indicate the median estimate. *Y*-axes are in units of effective number of subcommunities. The *x*-axes show weekly estimates from the even-numbered years from 1974 through 2008.

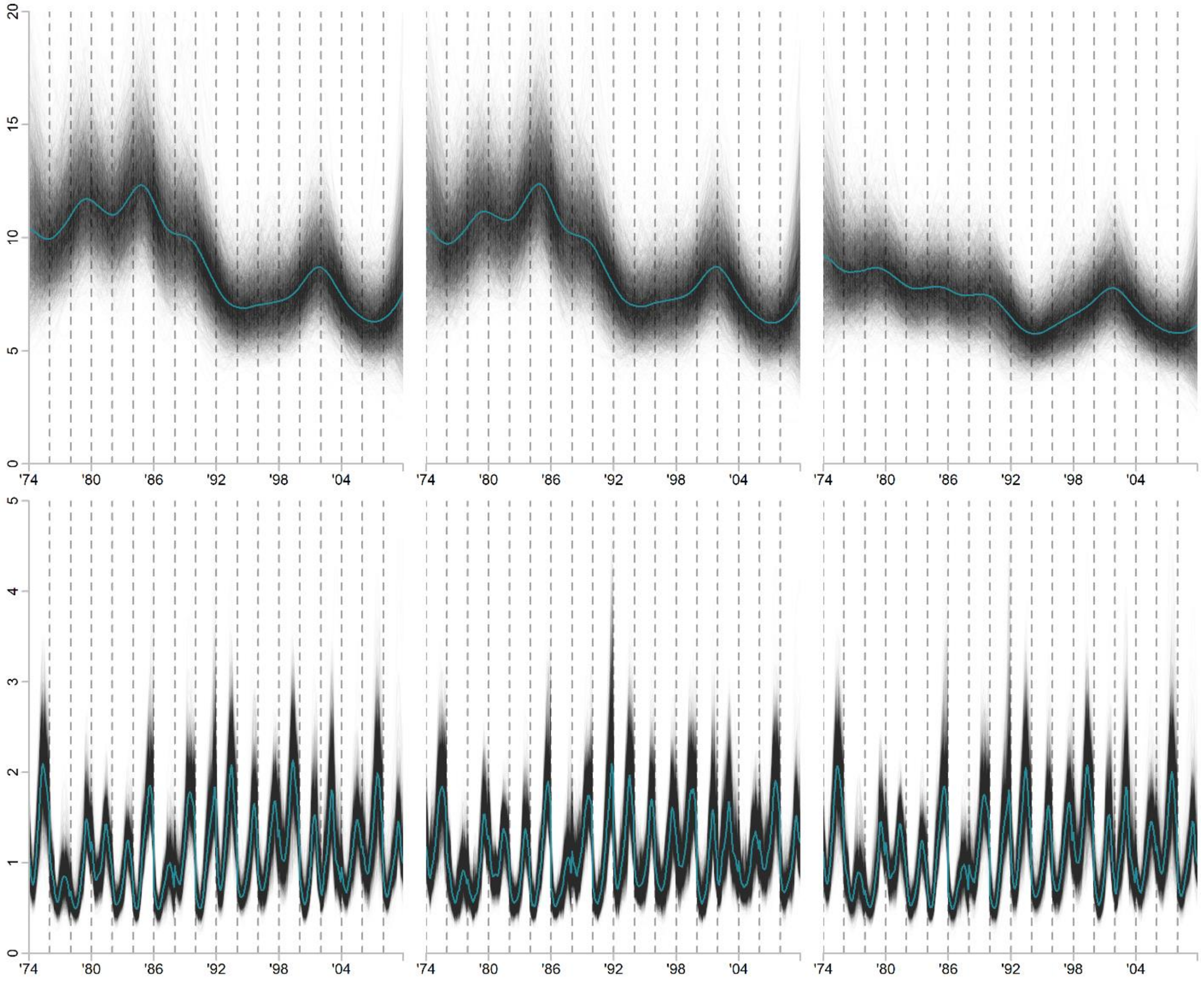


**Fig. SR27. IRW-climate models of β-diversity *q* = 2.** Columns from left to right show the model for 1) the total community, 2) the total community excluding the 208 genera in cluster14, and 3) only eukaryote genera. Greyscale lines show 5,000 posterior draws from the trend state component in the top panel and the regression component in bottom panel show. Teal lines indicate the median estimate. *Y*-axes are in units of effective number of subcommunities. The *x*-axes show weekly estimates from the even-numbered years from 1974 through 2008.

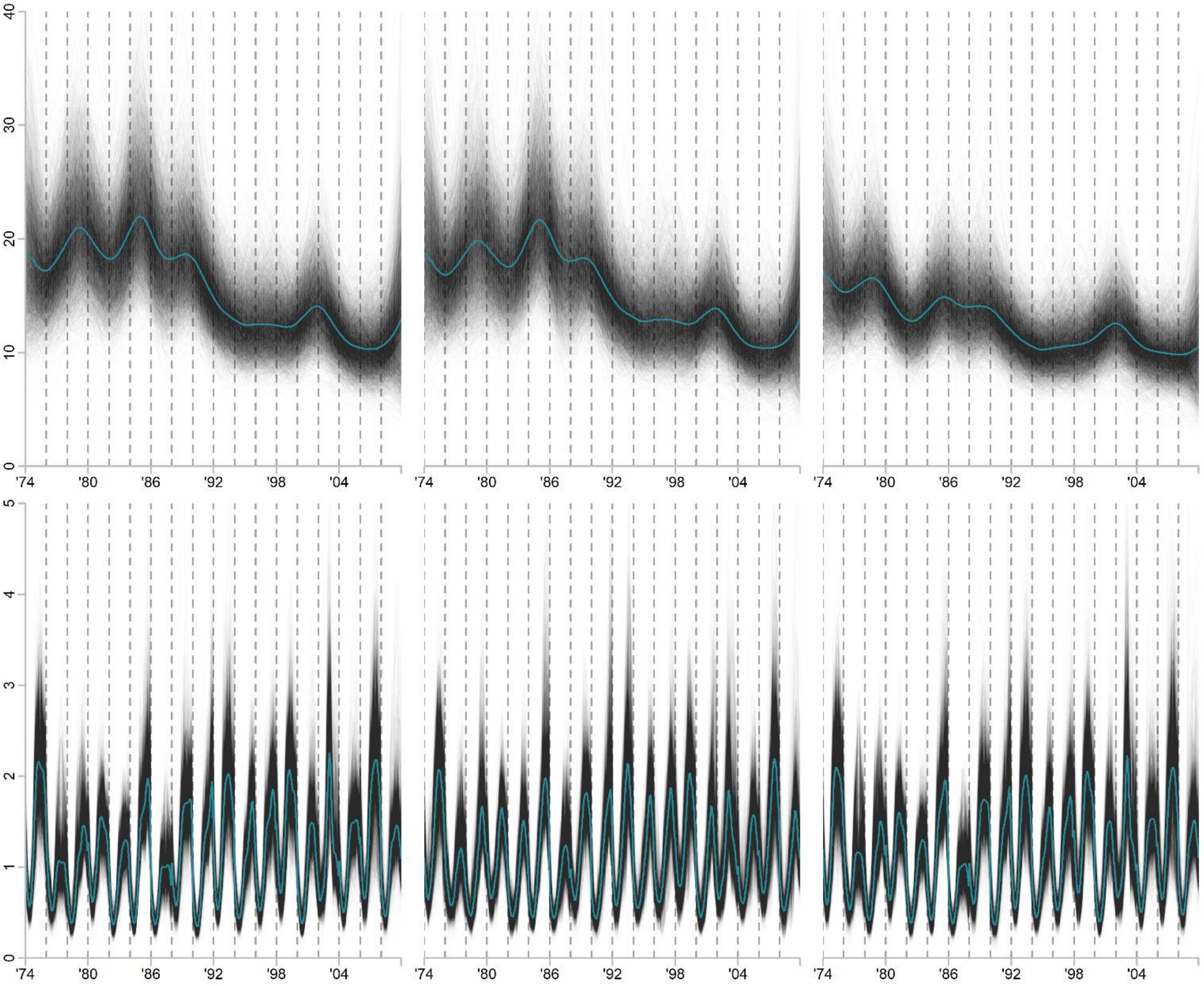


**Fig. SR28. IRW-climate models of β-diversity *q* = 3.** Columns from left to right show the model for 1) the total community, 2) the total community excluding the 208 genera in cluster14, and 3) only eukaryote genera. Greyscale lines show 5,000 posterior draws from the trend state component in the top panel and the regression component in bottom panel show. Teal lines indicate the median estimate. *Y*-axes are in units of effective number of subcommunities. The *x*-axes show weekly estimates from the even-numbered years from 1974 through 2008.

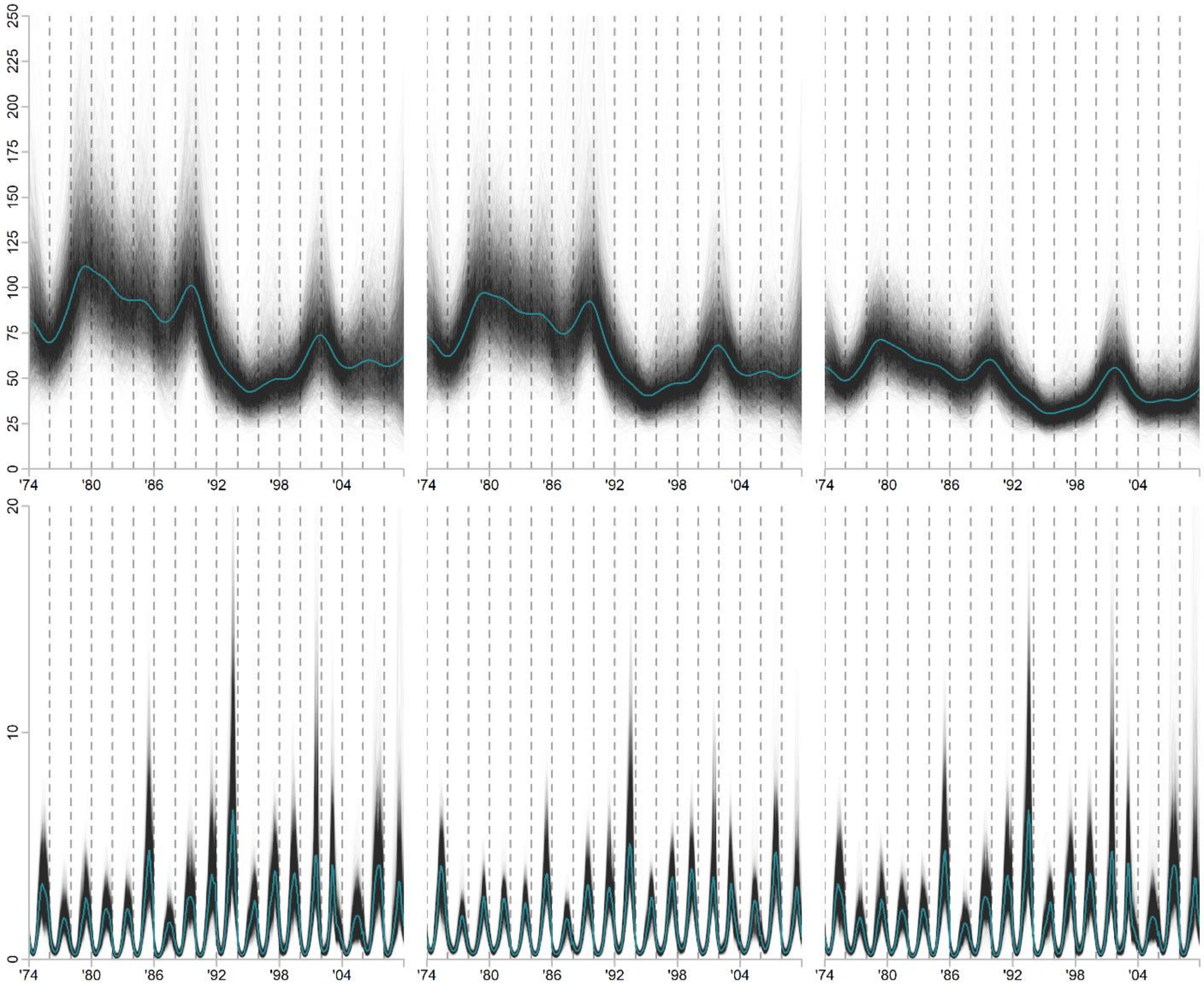


**Fig. SR29. IRW-climate models of γ-diversity *q* = 1.** Columns from left to right show the model for 1) the total community, 2) the total community excluding the 208 genera in cluster14, and 3) only eukaryote genera. Greyscale lines show 5,000 posterior draws from the trend state component in the top panel and the regression component in bottom panel show. Teal lines indicate the median estimate. *Y*-axes are in units of effective number of genera. The *x*-axes show weekly estimates from the even-numbered years from 1974 through 2008.

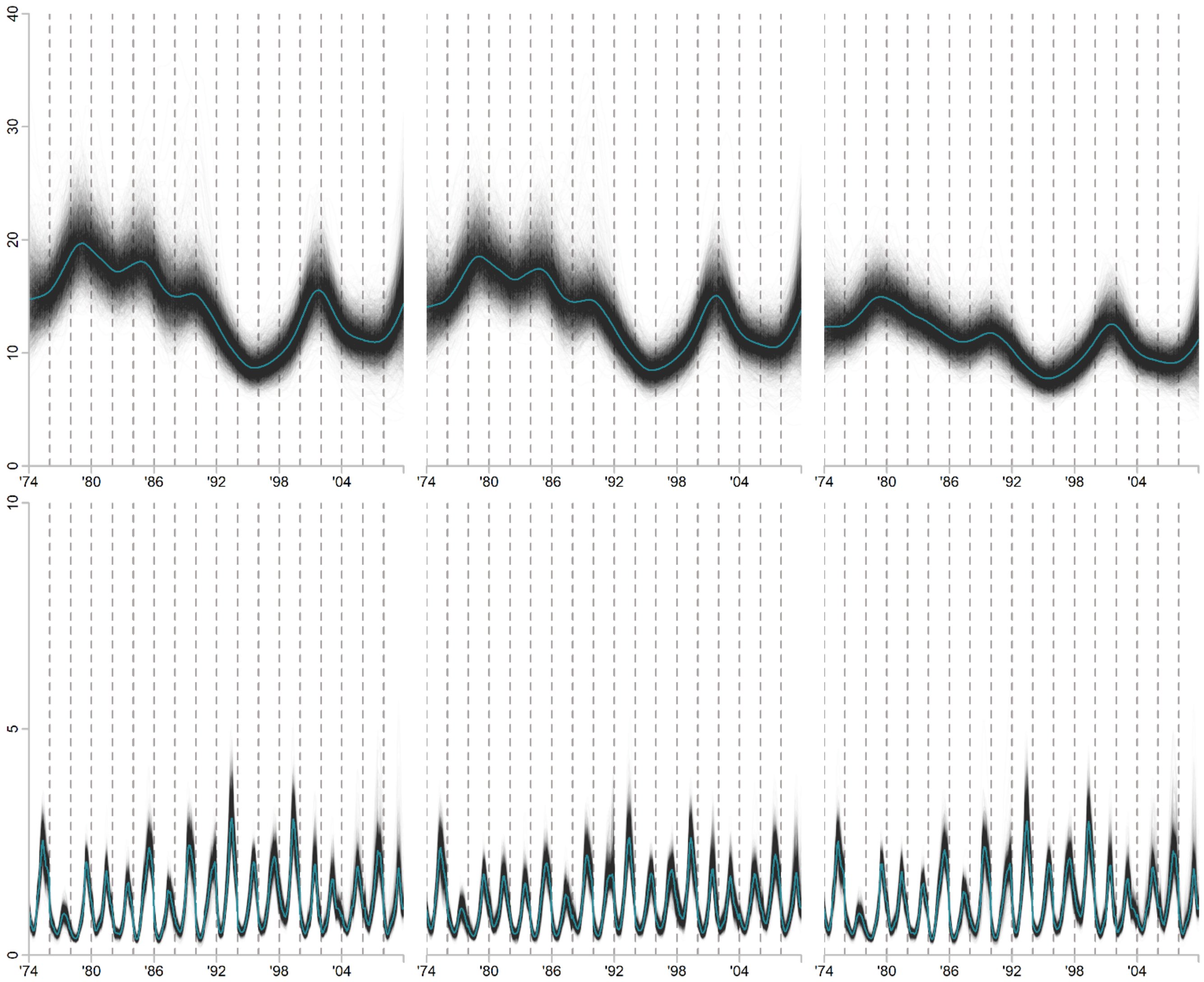


**Fig. SR30*. IRW-climate models of γ-diversity $q$ = 2.*** Columns from left to right show the model for 1) the total community, 2) the total community excluding the 208 genera in cluster14, and 3) only eukaryote genera. Greyscale lines show 5,000 posterior draws from the trend state component in the top panel and the regression component in bottom panel show. Teal lines indicate the median estimate. *Y*-axes are in units of effective number of genera. The *x*-axes show weekly estimates from the even-numbered years from 1974 through 2008.

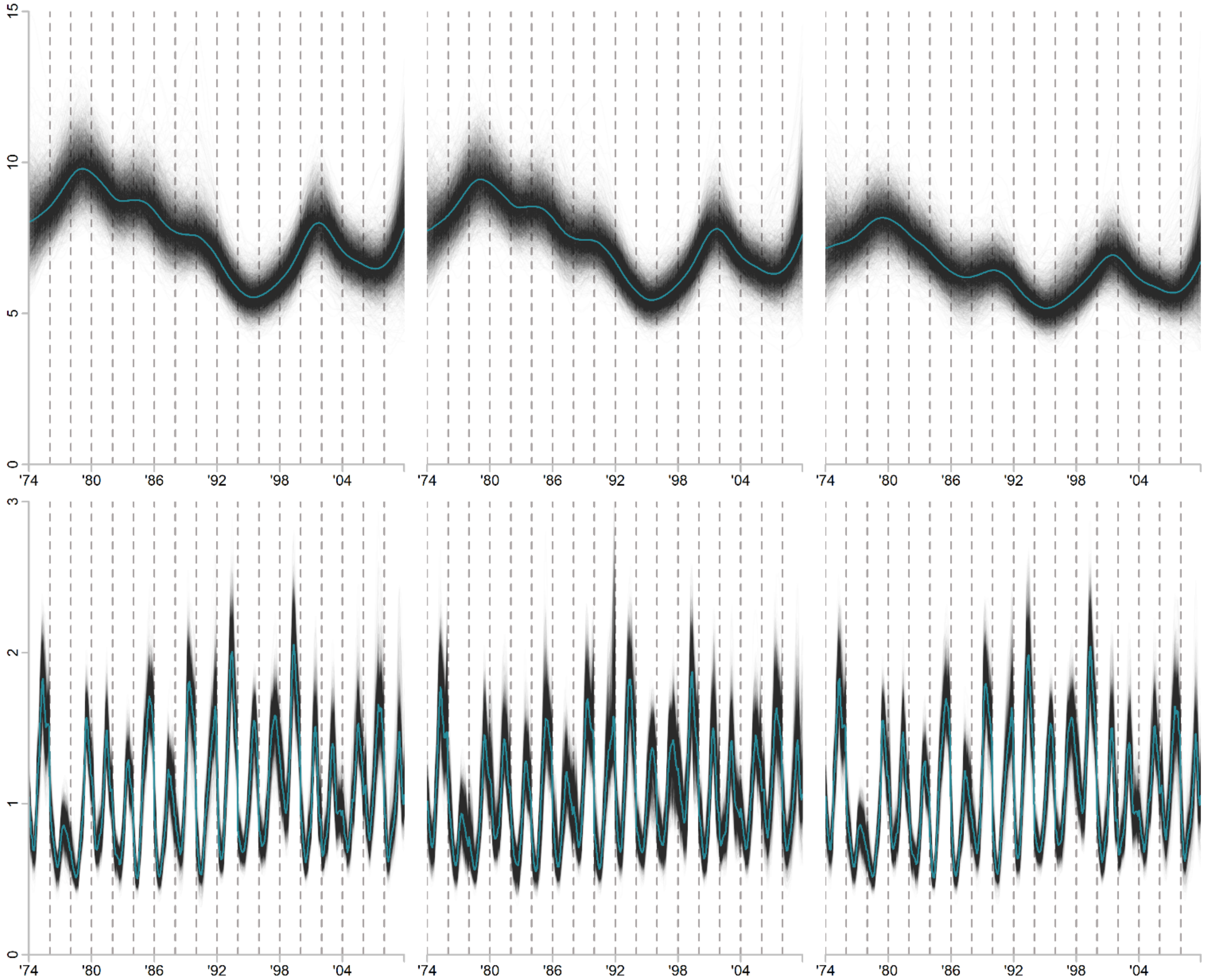


**Fig. SR31. IRW-climate models of γ-diversity *q* = 3.** Columns from left to right show the model for 1) the total community, 2) the total community excluding the 208 genera in cluster14, and 3) only eukaryote genera. Greyscale lines show 5,000 posterior draws from the trend state component in the top panel and the regression component in bottom panel show. Teal lines indicate the median estimate. *Y*-axes are in units of effective number of genera. The *x*-axes show weekly estimates from the even-numbered years from 1974 through 2008.
